# Prefrontal cortical dynorphin peptidergic transmission constrains threat-driven behavioral and network states

**DOI:** 10.1101/2024.01.08.574700

**Authors:** Huikun Wang, Rodolfo J. Flores, Hector E. Yarur, Aaron Limoges, Hector Bravo-Rivera, Sanne M. Casello, Niharika Loomba, Juan Enriquez-Traba, Miguel Arenivar, Queenie Wang, Robert Ganley, Charu Ramakrishnan, Lief E Fenno, Yoon Kim, Karl Deisseroth, Grace Or, Chunyang Dong, Mark A. Hoon, Lin Tian, Hugo A. Tejeda

## Abstract

Prefrontal cortical (PFC) circuits provide top-down control of threat reactivity. This includes ventromedial PFC (vmPFC) circuitry, which plays a role in suppressing fear-related behavioral states. Dynorphin (Dyn) has been implicated in mediating negative affect and mal-adaptive behaviors induced by severe threats and is expressed in limbic circuits, including the vmPFC. However, there is a critical knowledge gap in our understanding of how vmPFC Dyn-expressing neurons and Dyn transmission detect threats and regulate expression of defensive behaviors. Here, we demonstrate that Dyn cells are broadly activated by threats and release Dyn locally in the vmPFC to limit passive defensive behaviors. We further demonstrate that vmPFC Dyn-mediated signaling promotes a switch of vmPFC networks to a fear-related state. In conclusion, we reveal a previously unknown role of vmPFC Dyn neurons and Dyn neuropeptidergic transmission in suppressing defensive behaviors in response to threats via state-driven changes in vmPFC networks.

**Highlights:**

● vmPFC^Dyn^ neurons are activated by threats and threat-predictive cues
● Characterization of a genetically-encoded kappa-opioid receptor sensor
● vmPFC^Dyn^ neurons rapidly release Dyn in response to threats and their predictors
● vmPFC^Dyn^ signaling opposes threat-induced passive defensive behaviors
● Dyn signaling promotes threat-evoked state transitions in vmPFC networks

## Introduction

To ensure survival, organisms must respond to threats and mount appropriate defensive behaviors to minimize danger associated with threat and maximize other goal-directed behaviors essential for survival. The medial prefrontal cortex (mPFC) plays a critical role in mediating threat-induced defensive behaviors, with the ventral mPFC (vmPFC) playing a fundamental role in inhibiting fear-related responses to threats (Quirk et al., 2000, Sierra-Mercado et al., 2011, Moscarello and LeDoux, 2013, Do-Monte et al., 2015, Bravo-Rivera et al., 2015). Dysregulation of threat-processing and reactivity is also observed in many neuropsychiatric disorders (Penninx et al., 2021, Alexandra Kredlow et al., 2022).

The dynorphin (Dyn) / kappa-opioid receptor (KOR) system has been implicated in mediating stress-induced adaptive and mal-adaptive behaviors (Bruchas et al., 2010, Crowley et al., 2016, Tejeda et al., 2012, Massaly et al., 2016, Cahill et al., 2022). The Dyn / KOR system is highly expressed in limbic and cortical regions, including the mPFC, that support affective processing and motivation. However, most studies on the Dyn / KOR system have been aimed at elucidating the role of this system in regulating motivationally-charged behavior by focusing on subcortical limbic circuits, namely the ventral striatum, midbrain, and extended amygdala (Crowley et al., 2016, Al-Hasani et al., 2015, Shippenberg, 2009, Bals-Kubik et al., 1993, Castro and Bruchas, 2019, Baird et al., 2021, Massaly et al., 2019, Polter et al., 2017, Graziane et al., 2013, Bloodgood et al., 2021). As such, our understanding of how the Dyn / KOR system is embedded in mPFC networks and regulates threat-induced defensive behaviors in the face of threats is extremely limited. KOR activation in the dorsal mPFC (dmPFC) promotes aversion (Bals-Kubik et al., 1993, Tejeda et al., 2013), anxiety-like behavior (Tejeda et al., 2015), depressive-like behaviors (Wang et al., 2023), and disrupts working memory (Abraham et al., 2021). Conversely, activation of KORs in the vmPFC promotes anxiolytic behavior (Wall and Messier, 2000, Wall and Messier, 2002). This is consistent with literature demonstrating separable roles of dmPFC and vmPFC in the affective processing of threats (Alexandra Kredlow et al., 2022). Unfortunately, it is unclear whether Dyn-positive neurons in the vmPFC (vmPFC^Dyn^ neurons) undergo changes in activity in response to threats to influence defensive behavior. Moreover, due to technical limitations hindering our ability to detect neuropeptide release in freely-moving animals, it is also unclear whether mPFC^Dyn^ cells release opioid peptides to influence emotional behavior. Lastly, there is currently no information on how Dyn, and generally neuropeptides as a whole, shapes prefrontal cortical information processing during salient behavioral states, such as fear elicited by threats.

Here, we utilized an interdisciplinary approach to dissect the function of the vmPFC Dyn / KOR system in regulating passive defensive behaviors in response to threats. We employed *in-vivo* bulk and single-cell Ca^2+^ imaging to demonstrate that vmPFC^Dyn^ neuron activity is bidirectionally modulated with increases and decreases in activity with the presence and absence of acute threat, respectively. Moreover, we characterize a fluorescence-based KOR sensor to probe *in-vivo* dynamics of the vmPFC Dyn / KOR system and utilize this tool to reveal rapid Dyn release in response to threats and threat-predictive cues. We also demonstrate that inhibiting vmPFC-derived Dyn transmission promotes passive defensive behaviors in response to threats. Finally, we demonstrate that Dyn-mediated signaling promotes a shift of vmPFC network states post threat encounter using single cell Ca^2+^ imaging in mice lacking vmPFC Dyn-mediated signaling.

## Results

### Dynamics of vmPFC^Dyn^ neurons track threat exposure and threat-predictive cues

The Dyn / KOR system has been implicated in mediating negative affective states and the aversive components of stress (Bruchas et al., 2010, Crowley et al., 2016, Tejeda et al., 2012, Massaly et al., 2016, Cahill et al., 2022). Since the vmPFC is a brain region involved in modulating threat reactivity (Quirk et al., 2000, Sierra-Mercado et al., 2011, Do-Monte et al., 2015, Bravo-Rivera et al., 2015), we determined whether vmPFC^Dyn^ cells were activated during threat conditioning procedures utilizing fiber photometry in prodynorphin (PDyn)-Cre mice expressing Cre-dependent GCaMP7s in the vmPFC (Fig. 1A). In animals undergoing threat (fear) conditioning (Fig. 1B), vmPFC^Dyn^ neurons exhibited excitatory GCaMP7s responses to footshocks (Fig. 1D; Fig. S1B). During early trials, there was a slight increase in GCaMP activity to the tone. However, with repeated tone-shock pairings, Ca^2+^ activity increased during the presentation of the footshock-predictive tones (Fig. 1D,E; Fig.S1B,C). There were no tone-evoked responses in separate mice exposed to tones but no shocks (Fig. 1D,E). Interestingly, vmPFC^Dyn^ cells immediately adapted and stopped responding to the footshock-predictive tones when the shock was omitted during fear extinction procedures despite no change in freezing behavior (Fig. 1C-E; Fig. S1B,D). Collectively, these studies indicate that vmPFC^Dyn^ neurons respond to threats and threat-predictive tones and rapidly decrease their activity as threat outcomes change.

**Figure 1:**
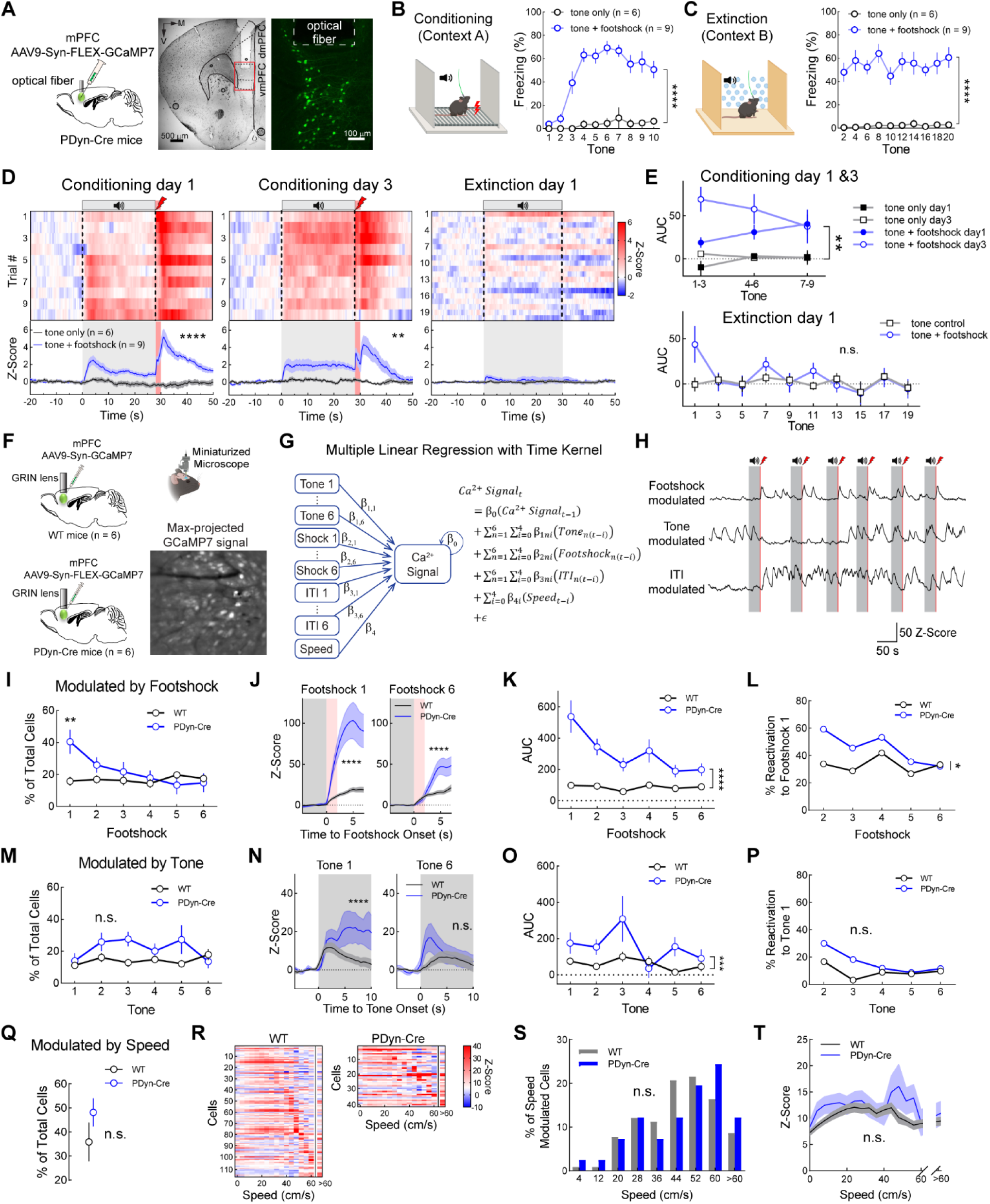
In-vivo monitoring of mPFC^Dyn^ cells during fear conditioning. A) Schematic viral expression and fiber photometry recording. Representative image of GCaMP7s expression and optical fiber tract in the vmPFC of a PDyn-Cre mouse. B) Threat conditioning paradigm in context A and freezing behavior in tone control and tone+footshock mice (Two-way ANOVA, group main effect \*\*\*\**p*<0.0001). C) Freezing during the threat recall / extinction in context B. Data is binned every 2 trials (Two-way ANOVA, group main effect \*\*\*\**p*<0.0001). D) Top: heat map of mean responses in mice undergoing associative threat conditioning and extinction. Bottom: time course of averaged GCaMP response during the whole session in mice exposed to tones alone (black) or to tones paired with footshocks (blue) (Two-way ANOVA, group main effect, \*\**p*=0.0054, \*\*\*\**p*<0.0001). E) Area under curve (AUC) of GCaMP response during conditioning and extinction. Data is binned every 3 trials for conditioning and every 2 trials for extinction (Conditioning day 1&3: Three-way ANOVA, \*\**p*=0.0016 between tone only vs tone + footshock group). F) Single-cell Ca^2+^ activity from global vmPFC neurons or vmPFC^Dyn^ neurons was recorded utilizing miniaturized microendoscopic imaging using GRIN lenses from WT and PDyn-Cre mice, respectively. G) Diagram and function of multiple linear regression (MLR) model with time kernel. H) Representative GCaMP activities from neurons that are significantly modulated by footshock (red area), tone (gray area), and ITI. I) Percentage of neurons significantly modulated by footshock (Two-way ANOVA, cell x trial interaction *p*=0.0053, \*\**p*=0.0019 with Bonferroni’s post-hoc test). J) Increased Ca^2+^ responses evoked by footshocks in footshock modulated neurons from PDyn-Cre mice relative to the global population (Two-way ANOVA, cell type main effect, \*\*\*\**p*<0.0001). K) AUC of Ca^2+^ responses during the footshock period (Two-way ANOVA, cell type main effect, \*\*\*\**p*<0.0001). L) The percentage of footshock encoding neurons during footshook 1 that were also significantly activated during all subsequent footshocks (paired t-test, \**p*=0.0496). M) Same as I, but for tones (Two-way ANOVA, cell main effect, *p*=0.0728). N) Same as J, but for tones (Two-way ANOVA, cell main effect, \*\*\*\**p*<0.0001). O) Same as K, but for tones (Two-way ANOVA, cell main effect, \*\*\**p*=0.0002). P) Same as L, but for tones (Paired t-test, \**p*=0.0880). Q) Percentage of neurons whose activity was predicted by speed (unpaired t-test, *p*=0.2364). R) Heatmaps of speed modulated cells sorted by maximal activity relative to speed from WT and PDyn-Cre mice. S) Distribution of speed-modulated neurons across distinct speed bins (Chi-square test, *p*=0.8251). T) Ca^2+^ responses as a function of speed in WT and PDyn Cre mice (RM two-way ANOVA,cell main effect *p*=0.1291). Data are expressed as the mean and error bars represent the SEM in this figure and all subsequent figures.

We next determined whether vmPFC^Dyn^ cells are preferentially activated by footshocks and shock-predictive tones relative to global vmPFC neurons. We performed single-cell imaging experiments from pan-neuronal GCaMP7-expressing cells in the vmPFC of WT mice and Cre-dependent GCaMP7-expressing cells in PDyn-Cre mice (Fig. 1F). A multiple linear regression model with a time kernel was used to identify behavioral- and task-related features that significantly predicted Ca^2+^ activity in individual neurons (Fig. 1G,H and methods). Using this model we observed that a larger fraction of vmPFC^Dyn^ neurons were responsive to footshocks in early trials (Fig. 1I), these neurons had larger Ca^2+^ responses across all trials (Fig. 1J,K), and were reactivated by footshocks with higher probability (Fig. 1L) than the global vmPFC population. No differences in the number of tone-encoding vmPFC^Dyn^ and global vmPFC cells or their probability of reactivation were observed (Fig. 1M,O,P), however vmPFC^Dyn^ neuron Ca^2+^ responses were increased primarily during early tone presentations (Fig. 1N). Lastly, the fraction and response magnitude of ITI-(Fig. S1F-I) or speed-encoding (Fig. 1Q-T) neurons from recordings obtained from WT and PDyn-Cre mice did not differ. Collectively, these results demonstrate that vmPFC^Dyn^ neurons are highly responsive to threats and their predictors relative to the global population of vmPFC neurons. vmPFC^Dyn^ neurons also rapidly adapt their activity as threat outcomes change.

### Anatomical and electrophysiological characterization of excitatory and inhibitory mPFC^Dyn^ neurons

We subsequently determined the distribution of PDyn mRNA-containing neurons within the mPFC utilizing RNAscope^®^ *in-situ* hybridization (Wang et al., 2012). mPFC^Dyn^ neurons are localized throughout various mPFC layers, with most cells in layer II/III expressing PDyn mRNA relative to the sparse population of mPFC^Dyn^ cells localized in deeper layers (Fig. 2A,B; Fig. S2B,C). To determine whether PDyn mRNA was differentially expressed in excitatory and inhibitory neurons, we examined co-localization of PDyn mRNA with VGluT1 and VGAT mRNA, markers of excitatory and inhibitory neurons, respectively (Fig. 2C,D). PDyn mRNA was primarily expressed in excitatory neurons and, to a lesser extent, inhibitory neurons in the dmPFC and vmPFC (Fig. 2D). PDyn mRNA expression from total excitatory and inhibitory neurons was similar in VGluT1- and VGAT-positive neurons (Fig. S2A). Layer II/III contained more excitatory mPFC^Dyn^ neurons, with both excitatory and inhibitory neurons present in deeper layers (Fig. S2B,C).

**Figure 2.**
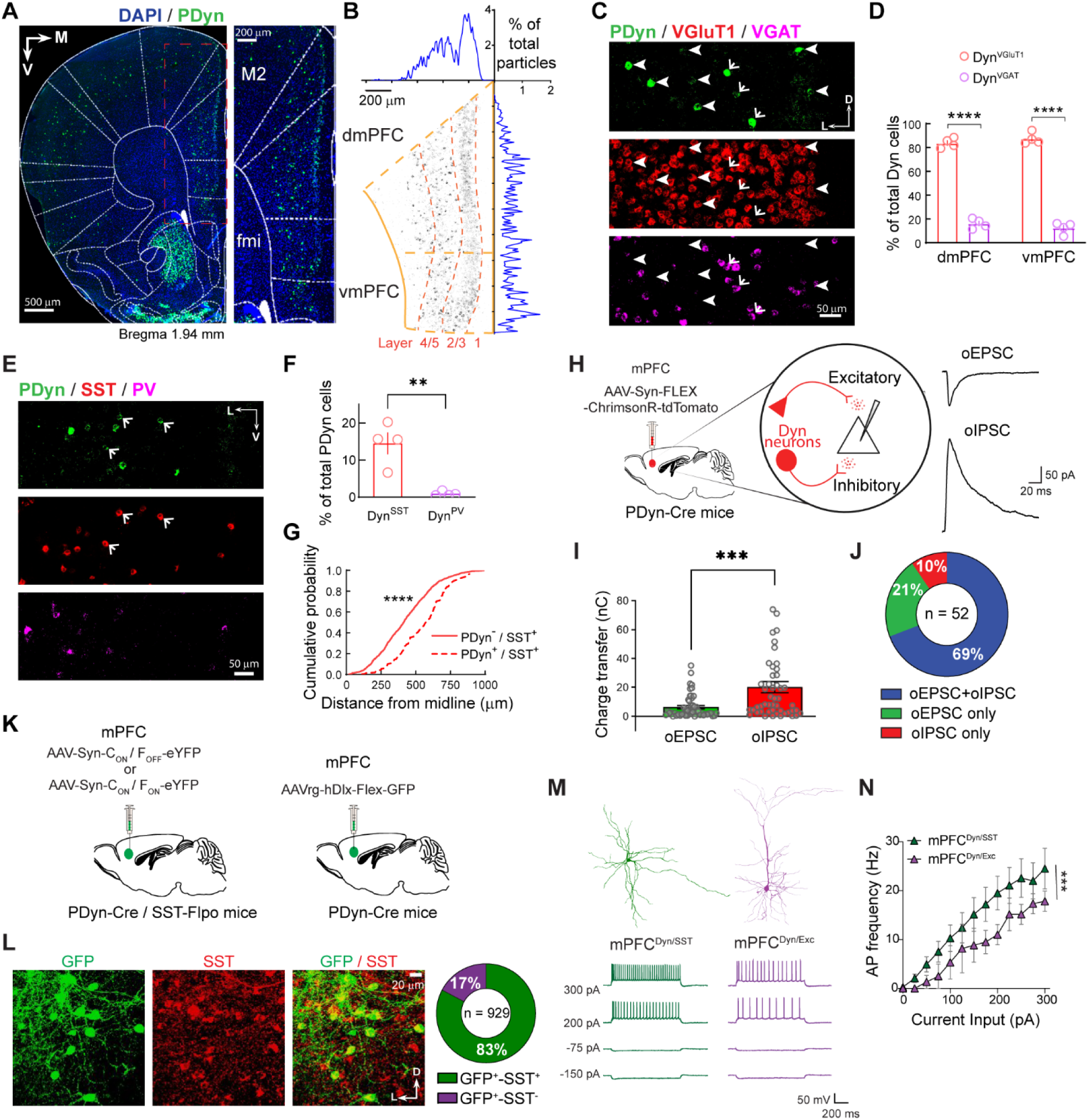
Anatomical characterization of mPFC dynorphin-expressing neurons. A) Representative image of PDyn mRNA (green) expression in the mPFC. B) Medial-lateral and dorso-ventral gradients of PDyn mRNA expression in the mPFC. Black dots are PDyn mRNA signals averaged from mPFC sections (bregma 1.94 mm) of 4 mice. C) Representative image of PDyn (green), VGluT1 (red) and VGAT (magenta) mRNA expression in mPFC. Arrows and arrowheads indicate the co-expression of PDyn in excitatory (VGluT1) and inhibitory (VGAT) neurons, respectively. D) PDyn mRNA is primarily expressed in excitatory neurons in both the dmPFC and vmPFC (Two-way ANOVA, cell-type main effect \*\*\*\**p*<0.0001). E) Representative image of PDyn (green), SST (red), and PV (magenta) mRNA expression in mPFC. Arrows indicate the co-expression of PDyn in SST neurons. F) Mean percentage of PDyn mRNA-positive cells co-expressing SST or PV mRNA (unpaired t-test, \*\**p*=0.0032). G) PDyn^+^ SST interneurons are localized in deeper layers relative to PDyn^-^ SST interneurons (Kolmogorov-Smirnov test, \*\*\*\**p*<0.0001). H) Optogenetic approach to identify how excitatory and inhibitory mPFC^Dyn^ cells make functional synaptic connections within mPFC circuits. I) Mean monosynaptic oEPSC and oIPSC charge transfer (unpaired t-test, \*\*\**p*=0.0007). J) Proportion of mPFC neurons receiving monosynaptic oEPSCs, oIPSCs, or dual oEPSCs/oIPSCs. K) INTRSECT and hDlx promoter approaches to label mPFC^Dyn/SST^ and mPFC^Dyn/Exc^ neurons. L) Representative images of SST immunoreactivity (red) on hDlx-Flex-GFP expression (green) mPFC cells. Pie chart shows the proportion of SST-positive and negative hDlx-GFP-positive neurons. M) Representative reconstruction and electrophysiological traces from biocytin-filled mPFC^Dyn/SST^ and mPFC^Dyn/Exc^ neurons. N) Action potentials (AP) in response to depolarizing current pulses in mPFC^Dyn/SST^ and mPFC^Dyn/Exc^ neurons (Two-way ANOVA, cell-type main effect \*\*\**p*=0.0002).

Inhibitory interneurons are comprised of a plethora of sub-populations, with parvalbumin (PV) and somatostatin (SST)-expressing neurons collectively constituting the majority of inhibitory interneurons. To further dissect the sub-population of PDyn mRNA-expressing inhibitory neurons, we determined whether PDyn mRNA colocalized with PV or SST mRNA (Fig. 2E,F; Fig. S2D). We found that PDyn mRNA was largely restricted to SST mRNA-expressing neurons and sparsely expressed in PV mRNA-expressing neurons, consistent with previous reports (Loh et al., 2017, Sohn et al., 2014, Smith et al., 2019). The percentage of Dyn neurons that are VGAT-positive (14.3%) is the same proportion as Dyn neurons that are SST-positive (14.5%), suggesting that SST-positive mPFC^Dyn^ interneurons (mPFC^Dyn/SST^) make the overwhelming majority of inhibitory neurons that express PDyn. mPFC^Dyn/SST^ neurons were differentially localized across layers of the mPFC relative to Dyn-negative SST-positive interneurons, suggesting mPFC^Dyn/SST^ neurons represent a subpopulation of SST-positive interneurons (Fig. 2G).

In dorsal and ventral striatum, Dyn is exclusively expressed in dopamine (DA) D1 receptor-expressing neurons (Castro and Bruchas, 2019). We observed that only a subset of mPFC^Dyn^ cells expressed Drd1a mRNA, suggesting that exclusive PDyn co-localization with DA D1 receptor observed in striatal circuits does not apply to mPFC circuits (Fig. S2E). Further, crossing PDyn-Cre mice with Ai14 tdTomato reporter mice resulted in ectopic expression of tdTomato mRNA in PDyn mRNA negative neurons (Fig. S2F), emphasizing the importance of using viral approaches to study PFC PDyn neurons.

Excitatory and inhibitory mPFC^Dyn^ neurons may establish connections within mPFC circuits necessary for fast neurotransmission and/or neuropeptide release. We confirmed that mPFC^Dyn^ synapses were widespread throughout cortical layers and differed from arborization patterns of mPFC^Dyn^ cells in PDyn-Cre mice injected with AAV-hSyn-FLEX-tdTomato-T2A-Synaptophysin-eGFP into the mPFC (Fig. S2G). Further, tdTomato-positive neurons were immuno-positive for PDyn (Fig. S2H), demonstrating specific viral expression in mPFC^Dyn^ neurons. To determine whether excitatory and inhibitory mPFC^Dyn^ cells make functional monosynaptic connections within the mPFC, we recorded optogenetically-evoked biophysically-isolated monosynaptic excitatory postsynaptic currents (oEPSCs) or inhibitory postsynaptic currents (oIPSCs) in layer V mPFC principal neurons in PDyn-Cre mice expressing Cre-dependent Chrimson-tdTomato in the mPFC (Fig. 2H-J; Fig. S2I). The majority of cells received monosynaptic excitatory and inhibitory connections from mPFC^Dyn^ neurons with similar onset latencies (Fig. 2J; Fig. S2I). Despite larger populations of Dyn neurons belonging to excitatory sub-types relative to inhibitory Dyn interneurons, inhibitory synaptic transmission onto principal neurons was more robust than excitatory transmission (Fig. 2I). *Ex-vivo* whole cell electrophysiology recordings from mPFC^Dyn^ neurons in PDyn-Cre mice injected with AAV-EF1α-DIO-eYFP in the mPFC revealed mPFC^Dyn^ cell morphology and electrophysiological properties were consistent with mPFC^Dyn^ neurons primarily consisting of excitatory pyramidal neurons and a smaller fraction belonging to non-pyramidal inhibitory neurons (Fig. S2J,K). To further delineate differences between mPFC^Dyn^ excitatory and inhibitory neurons, we utilized two complementary approaches to label inhibitory mPFC^Dyn^ neurons: 1) intersectional genetic and viral (INTRSECT) approach by injecting AAV-EF1α-C_ON_/F_ON_-eYFP in the mPFC of PDyn-Cre mice crossed with SST-Flpo mice (PDyn-Cre/SST-Flpo) or 2) injecting PDyn-Cre mice with AAV-hDlx-FLEX-eGFP, which yielded a high localization of SST immunoreactivity in fluorophore-positive mPFC neurons (Fig. 2K,L; Fig S2P). Excitatory mPFC^Dyn^ neurons (mPFC^Dyn/Exc^) were labeled using an INTRSECT approach employing AAV-EF1α-C_ON_/F_OFF_-eYFP to exclude expression from mPFC^Dyn/SST^ neurons in PDyn-Cre/SST-Flpo mice. Both approaches of mPFC^Dyn/SST^ neuronal labeling primarily labeled non-pyramidal neurons retaining dendritic arborization near the soma, whereas mPFC^Dyn/Exc^ neuronal populations primarily labeled pyramidal neurons (Fig. 2M; Fig. S2M). mPFC^Dyn/SST^ neurons were more hyperexcitable than their excitatory counterparts (Fig. 2M,N; Fig. S2M), consistent with less complex morphology in mPFC^Dyn/SST^ neurons (Fig. S2N,O). Collectively, these results demonstrate that Dyn is expressed in both excitatory and inhibitory neurons in the mPFC that are both integrated into local circuits. This contrasts with primary sensory regions where Dyn expression is restricted to interneurons (Sohn et al., 2014, Smith et al., 2019).

### Excitatory and inhibitory mPFC^Dyn^ neurons are recruited by threats and threat-predictive tones

To determine whether different subpopulations of vmPFC^Dyn^ neurons are recruited by threat and their predictive cues, we utilized INTRSECT approaches. First, to determine whether vmPFC^Dyn/SST^ neurons were activated by footshocks and associated cues, we injected AAV-EF1α-C_ON_/F_ON_-GCaMP6m into the vmPFC of PDyn-Cre/SST-Flpo mice (Fig. 3A). Similar to recordings from unspecified populations of vmPFC^Dyn^ cells (Fig. 1D), we initially observed responses to footshocks and to the tones that predict footshocks as associative learning occurred (Fig. 3B,J; Fig. S3A). We determined whether Dyn-lacking SST neurons differentially responded to threats and associated cues relative to Dyn-expressing counterparts by injecting PDyn-Cre/SST-Flpo mice with AAV-EF1α-C_OFF_/F_ON_-GCaMP6m (Fig. 3C). Dyn-negative SST neurons responded to footshocks with small tone-evoked responses (Fig. 3D,J; Fig. S3B). Lastly, we determined whether vmPFC^Dyn^ excitatory neurons were also recruited during threat conditioning procedures by injecting PDyn-Cre/SST-Flpo mice with AAV-EF1α-C_ON_/F_OFF_-GCaMP6m to exclude GCaMP expression from inhibitory neurons and restrict it to excitatory vmPFC^Dyn^ neurons (Fig. 3E). This INTRSECT strategy labels excitatory pyramidal neurons that differ from inhibitory SST Dyn neurons (Fig. S2L-O). However, it is possible this approach may also label sparse non-SST inhibitory neurons, such as Dyn PV-positive interneurons that make up approximately 2% of total PV interneurons and 0.5% of mPFC^Dyn^ neurons and would potentially contribute to the GCaMP signal (Fig. 2F, Fig. S2D). Excitatory vmPFC^Dyn^ neurons responded to footshocks and associated tones (Fig. 3F,J; Fig. S3C). To determine whether vmPFC^Dyn^ neuron activity tracked updating of threat outcomes, mice underwent extinction procedures wherein shocks were omitted at the end of the tone period. Interestingly, vmPFC^Dyn/SST^ neurons were immediately inhibited during extinction procedures (Fig 3G,J; Fig. S3A), an effect that was absent in Dyn-negative SST neurons (Fig 3H,J; Fig. S3B). Excitatory mPFC^Dyn^ neurons displayed a rapid switch to biphasic responses consisting of an initial activation followed by inhibition that persisted during the entire extinction session (Fig 3I,J; Fig. S3C). Thus, mPFC^Dyn^ neurons are strongly activated by threats and their predictors and reduce their activity as threats are extinguished (Fig. S3D). These results provide novel evidence that vmPFC^Dyn^ cells are responsive to threats, and their activity is positively correlated with internal states driven by threats but not freezing behavior (Fig. S3D,E).

**Figure 3:**
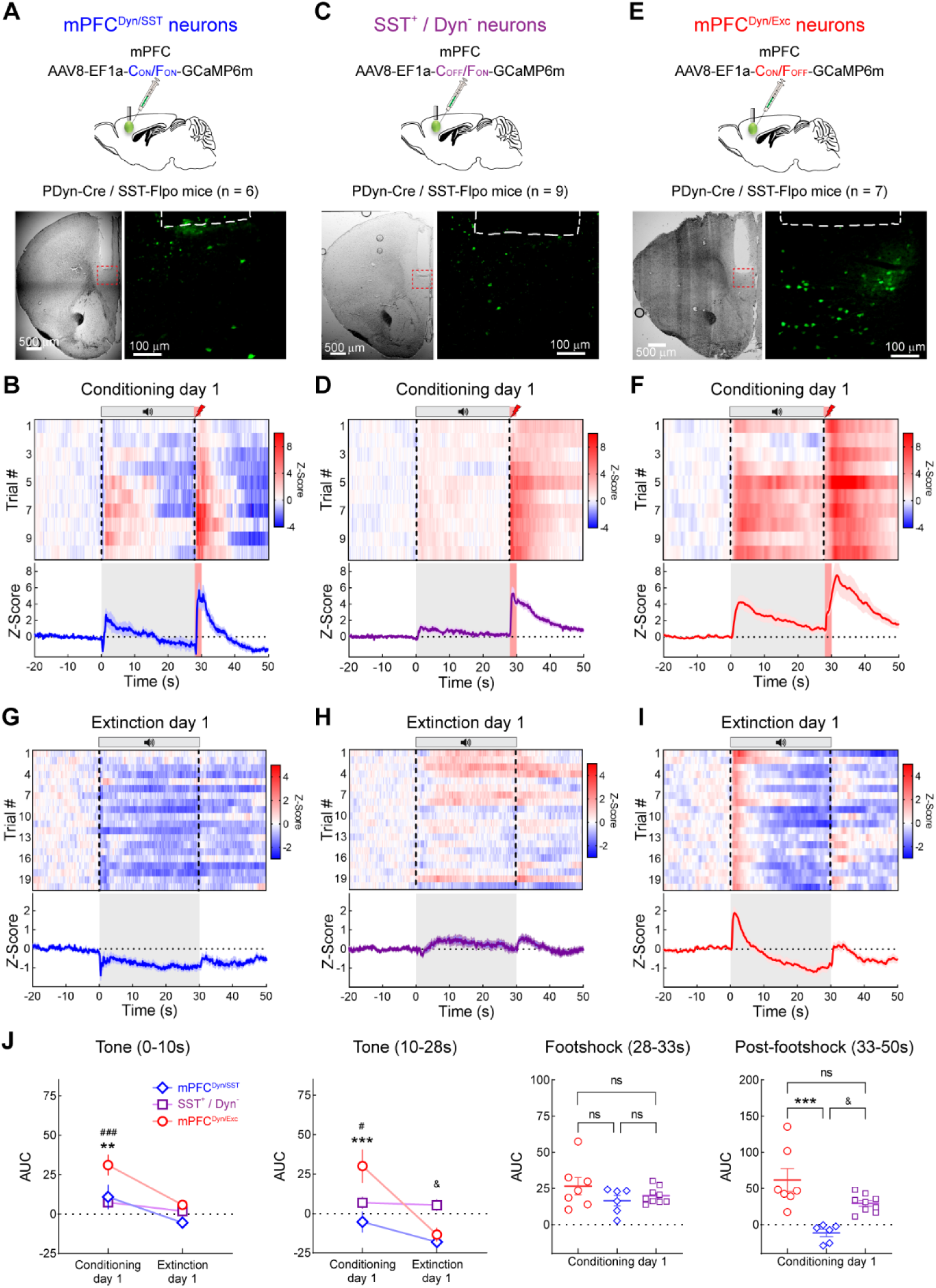
Excitatory and inhibitory vmPFC^Dyn^ cells track on-going threat states. A) Schematic depicting INTRSECT approach in PDyn-Cre / SST-Flpo mice to express GCaMP6m in mPFC^Dyn/SST^ cells. B) Mean heatmap across trials in mPFC^Dyn/SST^ neurons activity during threat conditioning procedures. Mean Z-Score of cue-shock evoked activity in mPFC^Dyn/SST^ neurons during threat across all trials. C) Experimental approach to monitor bulk Ca^2+^ activity in SST-positive Dyn-negative (SST^+^ / Dyn^-^) cells. D) As in B, but recording Ca^2+^ activity of SST^+^ / Dyn^-^ neurons during threat conditioning procedures. E) Experimental approach to monitor Ca^2+^ activity of mPFC^Dyn/Exc^ neurons. F) As in B, but recording Ca^2+^ activity of mPFC^Dyn/Exc^ neurons. G) Ca^2+^ dynamics in mPFC^Dyn/SST^ neurons during threat extinction. H) Ca^2+^ dynamics in SST^+^ / Dyn^-^ neurons during threat extinction. I) Ca^2+^ dynamics in mPFC^Dyn/Exc^ neurons during threat extinction. J) AUC of Ca^2+^ activity during early tone (0-10s, Two-way ANOVA, \*\**p*=0.0043 between mPFC^Dyn/SST^ and mPFC^Dyn/Exc^ neurons, ^###^*p*=0.0001 between SST^+^/Dyn^-^ and mPFC^Dyn/Exc^ neurons), late tone (10-28s, Two-way ANOVA, \*\*\**p*=0.0008 between mPFC^Dyn/SST^ and mPFC^Dyn/Exc^ neurons, ^#^*p*=0.0121 between SST^+^/Dyn^-^ and mPFC^Dyn/Exc^ neurons, ^&^*p*=0.0242 between mPFC^Dyn/SST^ and SST^+^/Dyn^-^ neurons), footshock (28-33s), and post footshock periods (33-50s, ANOVA, \*\*\**p*=0.0002 between mPFC^Dyn/SST^ and mPFC^Dyn/Exc^ neurons, ^&^*p*=0.0192 between mPFC^Dyn/SST^ and SST^+^/Dyn^-^ neurons).

### Threats and threat-predictive tones trigger rapid Dyn / KOR-mediated signaling

If vmPFC^Dyn^ neurons are releasing Dyn during footshock stress, then KOR activation should be observed. To this end, we utilized the KOR sensor kLight to monitor putative Dyn / KOR dynamics during threat conditioning procedures (Fig. 4A) (Patriarchi et al., 2018, Abraham et al., 2021). First, we characterized the kLight sensor in acute brain slices from WT mice injected with AAV-hSyn-kLight1.2a into the mPFC (Fig. 4B). Bath application of the KOR agonist U50,488 produced an increase in fluorescence, which was reversed by bath application of the competitive opioid receptor antagonist naloxone, or specific KOR antagonist nor-BNI (Fig. 4C; Fig. S4A). Similarly, Dyn A 1-17 bath application increased kLight fluorescence, while the non-opioid acting Dyn A 1-17 metabolite Dyn A 2-17 had no effect (Fig. 4D). kLight 0, a null control sensor, did not respond to U50,488 (Fig. 4C). Thus, kLight provides a measure of Dyn-induced KOR activation. We subsequently performed fiber photometry to validate this sensor *in-vivo* (Fig. 4E). Systemic administration of U50,488-induced increase of fluorescence was blocked by naloxone pre-treatment (Fig. 4F,G). Importantly, no changes in fluorescence were observed in mice injected with saline (Fig. 4G), as well as in mice expressing the control sensor kLight 0 injected with U50,488 (Fig. 4F).

**Figure 4:**
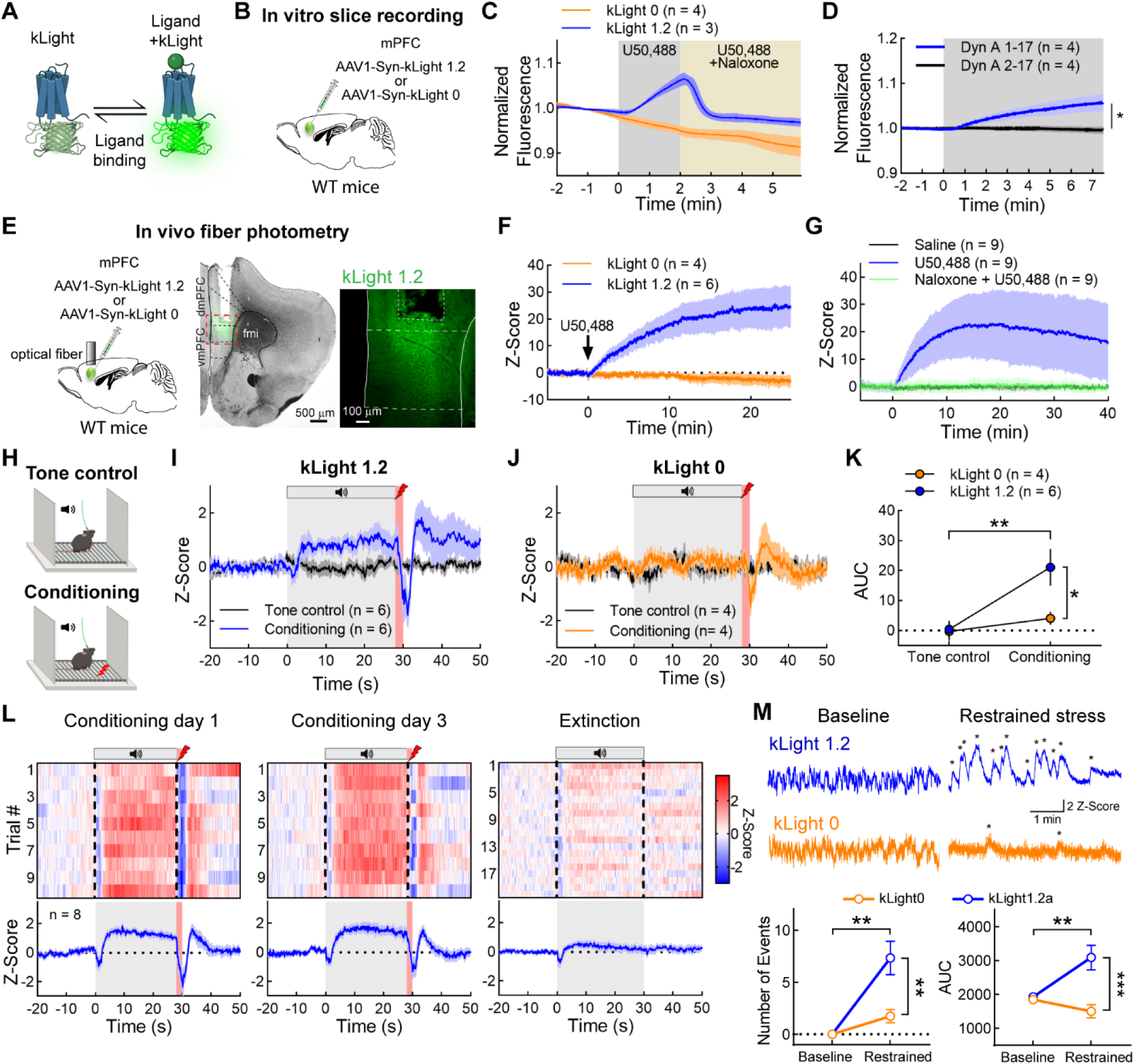
Threats and threat-predictive cues trigger rapid Dyn / KOR-mediated signaling. A) Schematic depicting kLight fluorescence in response to agonist activation. B) Schematic of AAV-hSyn-kLight1.2 or AAV-hSyn-SkLight0 expression in the mPFC of WT mice for in vitro slice recording. _C)_ U50,488 increases and naloxone inhibits fluorescence increases in slices expressing kLight1.2, but not kLight0. D) kLight1.2 responds to Dyn A 1-17, but not the opioid inactive metabolite Dyn A 2-17 (Two-way ANOVA, treatment main effect, \**p*=0.0161). E) Schematic and representative images depicting kLight expression in the vmPFC and implantation of optical fiber for fiber photometry recordings. F) Systemic KOR activation increases kLight1.2 fluorescence but not kLight0 in-vivo. G) Systemic KOR activation increases mPFC kLight1.2 fluorescence in-vivo in an opioid receptor-dependent manner. H) Schematic of tone control and threat conditioning sessions. I) kLight1.2 tracks shock-predictive cues during conditioning relative to sessions where mice are exposed to tones and no shocks. J) kLight0 does not fluoresce during threat conditioning procedures. K) AUC of kLight1.2 fluorescence in mice increased during threat conditioning relative to kLight0 (Two-way ANOVA with Bonferroni’s Post Hoc test, \*\**p*=0.0046 kLight 1.2a: Tone control vs. Conditioning, \**p*=0.034 Conditioning: kLight1.2 vs. kLight0). L) In separate mice kLight1.2 tracks threat-predictive cues across days and is not activated during threat extinction. M) kLight1.2 transients and AUC increase during restraint stress compared to kLight0 (Two-way ANOVA with Bonferroni’s Post Hoc test. Number of events: kLight 1.2a Baseline vs. Restrained (*p*=0.001), Restrained kLight1.2 vs. kLight0 (*p*=0.003); AUC: kLight 1.2a Baseline vs. Restrained (*p*=0.0087), Restrained: kLight1.2 vs. kLight0 (*p*=0.0005)).

We subsequently determined vmPFC Dyn / KOR dynamics during threat conditioning procedures (Fig. 4H). kLight fiber photometry recordings in the vmPFC revealed that mice initially displayed increased fluorescence to the footshock but minor responses to the footshock-predictive tone (Fig. 4I,L; Fig. S4B). With increased tone-shock pairings, kLight responses developed in response to the tone presentation, an effect that was not observed with the kLight 0 control sensor (Fig. 4I-K). A rapid dip in kLight 1.2 and 0 fluorescence was observed at the shock onset, which may reflect an artifact introduced by shock-induced escape responses. In a separate cohort of animals, conditioned kLight responses were absent during threat extinction procedures, consistent with our previous observations that vmPFC^Dyn^ neuron Ca^2+^ activity is rapidly attenuated or inhibited in response to shock omission (Fig. 4L; Fig. S4B,D). To determine whether kLight activity tracked other threat modalities, we determined the effect of restrained stress on kLight responses. Restrained stress caused an increase in transients in mice expressing kLight 1.2a in the vmPFC, while transients were rarely observed during baseline recordings (Fig 4M). This was not observed in mice expressing kLight 0. These results suggest threats activate vmPFC^Dyn^ neurons and lead to local KOR activation, presumably via Dyn released from vmPFC^Dyn^ neurons.

### vmPFC Dyn release limits threat-induced passive behavioral states

We subsequently determined whether vmPFC kLight responses, which provide an indirect index of KOR activation by an endogenous ligand, evoked by threats were mediated by vmPFC PDyn-derived neuropeptides. To this end, we injected WT mice with scrambled shRNA-tdTomato (control) or PDyn shRNA-tdTomato (shPDyn) in the vmPFC to knockdown vmPFC PDyn expression. In the same mice, we also injected AAV expressing kLight in the vmPFC to monitor kLight dynamics during threat conditioning and extinction (Fig. 5A; Fig. S5A). Here, we found a moderate time-dependent increase in kLight fluorescence evoked by footshock-associated tones in scrambled shRNA controls than in mice with vmPFC PDyn knockdown (Fig. 5B,D). Further, control mice displayed cue-evoked kLight responses on the first trial during threat extinction procedures that rapidly habituated after shock omission (Fig. 5C,D), but not in mice expressing PDyn shRNA in the vmPFC. Differences in threat-evoked kLight signaling were not due to differential kLight expression since both scrambled-and PDyn-shRNA expressing mice displayed similar U50,488-evoked increases in kLight fluorescence (Fig. 5E). These results are consistent with activation of vmPFC^Dyn^ neurons and endogenous Dyn release from vmPFC neurons during processing of on-going threats.

**Figure 5:**
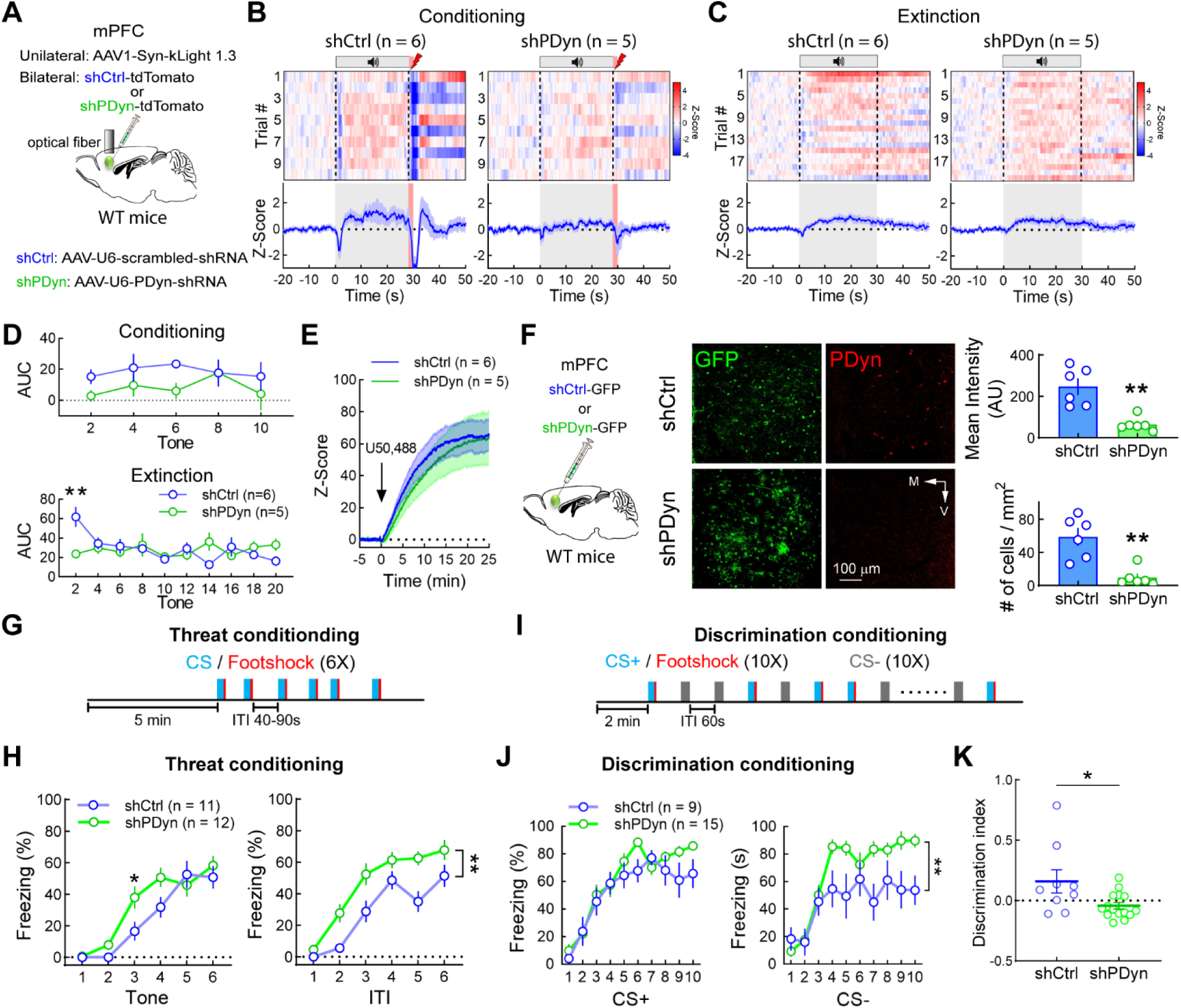
PDyn knockdown in the vmPFC limits passive threat driven states. A) Knockdown of PDyn expression using AAV expressing PDyn shRNA and expression of kLight1.3 in mPFC for fiber photometry recordings. B) Heatmaps and traces of kLight1.3 signals during conditioning in shCtrl and shPDyn mice (Two-way ANOVA, treatment (shRNA vs scramble shRNA) x Time interaction p<0.0001). C) Heatmaps and traces of kLight1.3 signals during extinction in shCtrl and shPDyn mice (Two-way ANOVA, treatment (shRNA vs scramble shRNA) x Time interaction p<0.0001). D) AUC analysis of kLight1.3 responses to tone during threat conditioning and recall/extinction. Data is binned every 2 trials (Two-way ANOVA, **p=0.0012 between tone 2 of shCtrl and shPDyn group). E) U50,488 similarly increases kLight fluorescence in control and PDyn shRNA-expressing mice. F) PDyn-immunoreactivity (red) in mice injected with scrambled shRNA-GFP (shCtrl) or PDyn shRNA-GFP (shPDyn) in the vmPFC and quantification of mean intensity and cell density (unpaired t-test, \*\**p*<0.01). G) Diagram of threat conditioning paradigm. H) Expression of PDyn shRNA potentiates freezing during the cue and intertrial interval (ITI) (Two-way ANOVA with Bonferroni’s Post Hoc test on tone, Trial x Treatment (shRNA vs control) interaction, \**p*<0.05; Treatment main effect for ITI, \*\**p*<0.01). I) Schematic of cue discrimination conditioning paradigm. J) In a cue discrimination threat-conditioning task, mice with vmPFC PDyn knockdown have increased freezing to the CS-relative to controls, but no difference to the CS+-evoked freezing (Two-way ANOVA with Bonferroni’s post-hoc test, shRNA main effect for CS-, \*\**p*=0.0187). K) Discrimination index ((Freezing_CS+_ - Freezing_CS-_)/(Freezing_CS+_ + Freezing_CS-_)) of control and PDyn knockdown mice (unpaired t-test, *p=0.0216)

To determine whether Dyn release by vmPFC^Dyn^ neurons modulates behavioral states driven by threats, we knocked down PDyn expression utilizing AAV expressing PDyn-shRNA-GFP (Fig. 5F). Mice expressing AAV-U6-PDyn-shRNA-GFP had decreased PDyn immunoreactivity in fibers and cell bodies in the vmPFC relative to scrambled shRNA controls (Fig. 5F). During threat conditioning, PDyn knockdown in the vmPFC robustly increased freezing during the intertrial interval (ITI) throughout the rest of the session and cue-elicited freezing during early tones (Fig. 5G,H). Freezing during the ITI reflects a persistent behavioral state generated by the footshock. During the ITI, threat appraisal is critical for establishing predictive external features (tone or context) of the footshock, evaluating appropriate defensive strategies given the environment, and balancing motor outputs that bias defensive behaviors towards passive states, including freezing, or active escape (Roelofs and Dayan, 2022). Increased freezing during the ITI (Fig. 5H) and early tones suggest that vmPFC Dyn-mediated signaling normally counteracts persistent freezing generated by acute threats. vmPFC PDyn shRNA mice did not display differences in freezing behavior during contextual and cued recall of threat memory (Fig. S5C), consistent with a rapid reduction in vmPFC^Dyn^ neuron activity and Dyn release (as assessed by kLight) during threat extinction (Fig. 1D,S3D, 4L, 5C). This implies that vmPFC Dyn cellular activity and peptidergic transmission may be critical for regulating a persistent threat-triggered behavioral state in the presence of on-going threats. Further, PDyn-loxP mice with intra-vmPFC infusions of AAV-Cre-GFP or AAV-eGFP displayed enhanced freezing relative to C57/Bl6J WT controls infused with AAV-Cre-GFP (Fig. S5B), suggesting that genotype differences in the PDyn-loxP mouse line drive differential threat-induced freezing and highlights the importance of both genotype and Cre-vector controls in conditional knockout studies. The vmPFC has been implicated in suppressing passive defensive behaviors (Moscarello and LeDoux, 2013, Bravo-Rivera et al., 2015, Do-Monte et al., 2015). We reasoned that vmPFC Dyn-mediated signaling may play a role in suppressing defensive states generated by threats. To this end, we tested mice in a cue discrimination task wherein a CS+ tone and CS-tone was co-terminated or not associated with a footshock during acquisition, respectively (Fig. 5I). In this task, during threat conditioning, control mice displayed lower levels of freezing to the CS-relative to the CS+ (Fig. 5J). However, we observed that vmPFC PDyn shRNA-expressing mice displayed elevated freezing response to the CS-relative to controls (Fig. 5J) and impaired ability to discriminate between CS+ and CS-tones (Fig. 5K). In accordance with a rapid ramping down in vmPFC^Dyn^ neuron activity and Dyn release during extinction (Fig. 1D,S3D,4L,5C), we observed that vmPFC Dyn knockdown did not impact freezing to the presentation of CS+ and CS-during the cued threat memory recall test when Dyn is not being released (Fig. S5D). These results suggest that vmPFC Dyn signaling facilitates the function of the vmPFC in inhibiting post-threat passive behavioral states.

Increased freezing in vmPFC PDyn shRNA mice during threat conditioning acquisition may stem from differences in locomotor activity, enhanced anxiety-like behavior, increased autonomic reactivity to threats, and/or increased working/short-term memory performance. However, no significant differences were observed in total locomotor activity in the center or periphery of an open field or during habituation periods in the acquisition and cued threat recall tests (Fig. S5E). No significant differences in anxiety-like behavior were observed in the light-dark box, elevated plus maze, and the open field test in mice with vmPFC PDyn knockdown (Fig. S5E-G). Moreover, threat-induced pupillometry responses were unaffected in PDyn shRNA-expressing mice, suggesting that increased freezing is not induced by changes in arousal (Fig. S5H). PDyn shRNA did not modify working and short-term memory on the spontaneous alternation task and the novel object recognition task (Fig. S5I,J), suggesting this effect was not due to differential short-term and working memory function, respectively. Lastly, vmPFC PDyn knockdown did not affect thermal and mechanical nociception, as well as CFA-induced hyperalgesia (Fig.S5K-M), suggesting that differences in freezing during acquisition of threat conditioning procedures are not mediated by differences in pain processing. Collectively, these studies demonstrate that vmPFC^Dyn^ cells, though diverse, are broadly tuned to detect threats. Dyn release in response to vmPFC^Dyn^ cell activation inhibits passive defensive behavioral states.

### vmPFC Dyn signaling modulates activity of threat-driven shifts in vmPFC neural dynamics

Dyn-mediated signaling may impact passive defensive behaviors upon acute threat exposure by regulating how vmPFC circuits process threats. We subsequently delineated how Dyn transmission may influence vmPFC circuits during threat exposure. To this end, we injected WT mice with GCaMP7-expressing virus and implanted a GRIN lens in the vmPFC to monitor vmPFC network activity and either AAV-U6-PDyn-shRNA-tdTomato in the vmPFC to knockdown PDyn expression (vmPFC^Dyn^ knockdown) or AAV-hSyn AAV-U6-scrambled-shRNA-tdTomato as a control (Fig. 6A). Examination of activity of all neurons recorded from controls and vmPFC knockdown mice revealed that the first footshock drove time-locked responses in subsets of vmPFC neurons recorded from control mice (Fig. 6B; cells are sorted by response magnitude to first footshock), but strikingly many neurons were subsequently activated and/or inhibited throughout the rest of the session (Fig. S6A). Session-wide threat-driven modulation of vmPFC neurons was not as pronounced in vmPFC^Dyn^ knockdown mice (Fig. 6B; Fig. S6A). Quantification and comparison of the distribution of mean Ca^2+^ activity (normalized to the baseline period of the task before the first cue) across all cells obtained from controls and vmPFC^Dyn^ knockdown mice (Fig. 6C-G) revealed that vmPFC^Dyn^ knockdown showed a narrowed distribution (closer to 0 in both negative and positive Z-Score regions) of Ca^2+^ activity across footshock and tone periods, as well as the ITI (Fig. 6D-G). This difference was not observed during baseline or the first tone presentation before it acquired salience through pairings with the footshock (Fig. 6C,E). These results demonstrate that upon exposure to footshocks (a direct threat), large subsets of vmPFC neurons are broadly activated and/or inhibited throughout the session, a process that is diminished in vmPFC^Dyn^ knockdown mice. It is not clear whether loss of threat-evoked session-wide modulation of activity with vmPFC^Dyn^ knockdown may influence encoding of distinct features of the task and the animals’ speed. Using a multiple linear regression encoder, we determined that the percentage and Ca^2+^ response of cells whose activity was significantly encoded by footshock (Fig. 6H-J), tone (Fig. 6K-M), ITI (Fig. 6N-P), or speed (Fig. 6Q-T) did not differ between controls and vmPFC^Dyn^ knockdown mice. These results suggest that attenuation of session-wide threat-driven shift in activity with vmPFC^Dyn^ knockdown is independent of specific encoding of task-related features and speed by vmPFC networks. Together, these data suggest that vmPFC^Dyn^ peptidergic transmission may regulate state transitions in vmPFC circuits at the population-level that persist beyond the acute threat.

**Figure 6:**
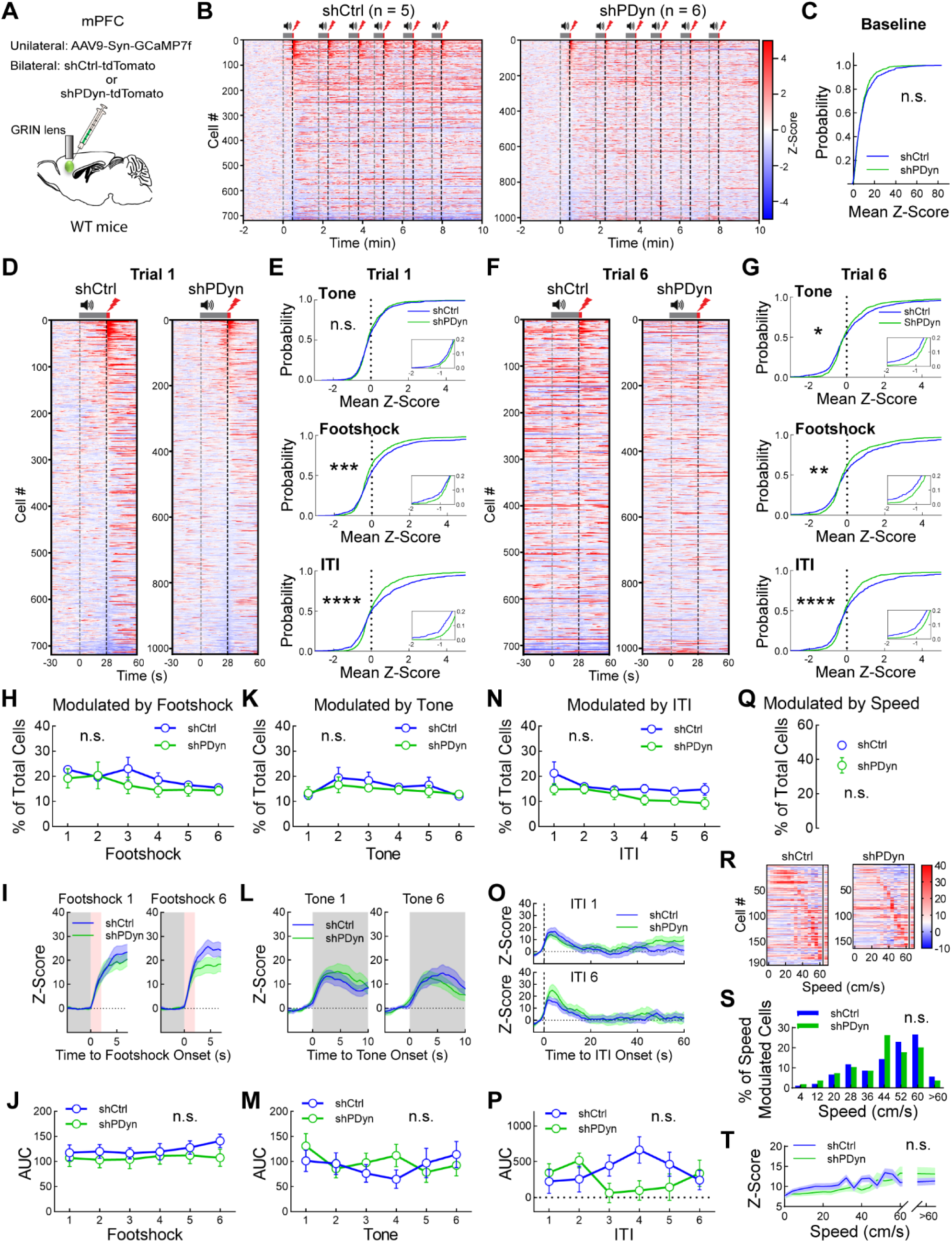
vmPFC Dyn signaling regulates activation and inhibition of vmPFC neurons during threat exposure. A) Experimental strategy to monitor single cell vmPFC Ca^2+^ dynamics using GCaMP in control mice injected with AAV-U6-scrambled shRNA-tdTomato or mice with PDyn knockdown using AAV-U6-PDyn shRNA-tdTomato. B) Population heatmap of Z-Scored GCaMP7 activity in control mice and mice with vmPFC PDyn knockdown across the whole session. Tone and shock presentations are depicted with speaker and shock symbols, respectively. Cells are sorted by Z-Scored activity during the five seconds starting from first shock onset. C) No significant difference of cumulative probability of mean Z-Score in control (blue) and PDyn-shRNA (green) mice during baseline periods. D) Heatmap of Z-Scored GCaMP7 activity in control mice and mice with vmPFC PDyn knockdown zoomed in on trial 1. E) Cumulative probability of mean Z-Score in control (blue) and PDyn-shRNA (green) mice during the Tone periods (top), shock periods (bottom), and ITI from the first trial. Insets are zoomed in between Mean Z-score −2 to −0.5. (Kolmogorov-Smirnov test, \*\*\**p*<0.001, \*\*\*\**p*<0.0001). F) Heatmap of Z-Scored GCaMP7 activity in control mice and mice with vmPFC PDyn knockdown zoomed in on trial 6. G) Same as C but for trial 6 (Kolmogorov-Smirnov test, \**p*<0.05, \*\**p*<0.01, \*\*\*\**p*<0.0001). H), K), N) Percentage of neurons modulated by footshocks (H), tones (K), ITI (N), and speed (Q) did not differ between control and PDyn shRNA mice.. I), L), O), T) Z-Scored GCaMP responses from cells whose activity was modulated by footshocks (I), tones (L), ITI (O), and speed (T) from control and PDyn shRNA mice. J), M), P) AUC of GCaMP responses from cells whose activity was modulated by footshocks (J), tones (M), ITI (P) from control and PDyn shRNA mice. R) Heatmaps of speed modulated cells sorted by maximal activity relative to speed from control and PDyn-shRNA mice. S) Distribution of speed-modulated neurons across distinct speed bins.

### PDyn transmission modulates vmPFC network dynamics

vmPFC^Dyn^ signaling may influence dynamics of vmPFC networks at the population level to influence processing beyond immediate threat experiences. To dissect the impact of vmPFC Dyn-mediated signaling on state transitions of vmPFC population dynamics induced by threat conditioning, we examined correlated activity of vmPFC networks on a “moment by moment” basis and across distinct trials (Fig. 7A; Fig. S6B). During baseline, vmPFC population dynamics did not correlate in both control and PDyn shRNA mice (Fig. 7A-D). We analyzed activity on a “moment by moment” basis within task epochs by plotting the mean Pearson correlation coefficients during the first 28 sec of the cue, 5 sec during and after the footshock, and during the ITI. The first presentation of the footshock increased correlated activity within the footshock period in control and PDyn shRNA groups (Fig. 7B). In controls, footshock delivery increased correlated activity that persisted throughout the rest of the session, including during the ITI and tone (Fig. 7C,D), consistent with a threat-driven state transition in vmPFC circuits. Specifically, correlated activity within the tone period increased with associative learning in control mice, suggesting that correlated vmPFC circuit activity tracks associative learning as well as the defensive/internal state that persists during the ITI. We subsequently determined whether vmPFC networks were correlated throughout the session. The correlation of activity state in vmPFC circuits during the first tone presentation (before it acquired incentive salience) and first footshock and subsequent trials decreased in control mice (Fig. 7E,F). These results suggest that in control mice, the population-level activity in vmPFC circuits may be differentially encoding the first and subsequent exposures to acute threats and footshock-predictive tones. For example, the first footshock has an element of surprise driven by unexpected nociception that is rapidly diminished with each subsequent footshock delivery and the first tone has not gained any predictive value. Interestingly, correlated activity during the first ITI after footshock and subsequent ITIs did not drop (Fig. 7G), suggesting that persistent correlated activity following tones and footshocks during the ITIs may be encoding a common vmPFC network state associated with threat processing, which may include top-down suppression of a passive defensive fear state during the ITI, a period where threat appraisal becomes particularly important. In contrast, mice with vmPFC^Dyn^ knockdown displayed similar increases in correlated activity in response to the first presentation of the tone and footshock (Fig. 7B,C). However, correlated activity during the ITI and subsequent tone presentations, but not footshock, was attenuated relative to control mice (Fig. 7B-D), suggesting that the enhanced activation of vmPFC neurons (Fig. 1), Dyn signaling (Fig. 4), and subsequent actions of Dyn on KORs in distinct vmPFC neurons and afferent terminals (Tejeda et al., 2013, Tejeda et al., 2015, Yarur et al., 2023, Wang et al., 2023) is critical for promoting a functional switch in population-level activity in response to threats that persists beyond acute threat exposure. Moreover, decreased correlated activity between the first tone and footshock and subsequent presentations was not different between vmPFC^Dyn^ knockdown and control mice (Fig. 7E,F). These results suggest that vmPFC networks lacking Dyn signaling are able to distinguish a novel tone that is benign and acquires salience with footshock pairing, as well as the on-going presentation of a footshock. These results are consistent with no changes in encoding of tone and footshock by individual vmPFC neurons. However, loss of vmPFC Dyn signaling decreased correlated activity across the first post-footshock ITI and subsequent ITIs, where vmPFC state transitions remain high in control mice (Fig. 7G). Collectively, these results suggest that PDyn transmission in the vmPFC promotes the engagement of a threat-associated network state that persists beyond the acute threat.

**Figure 7:**
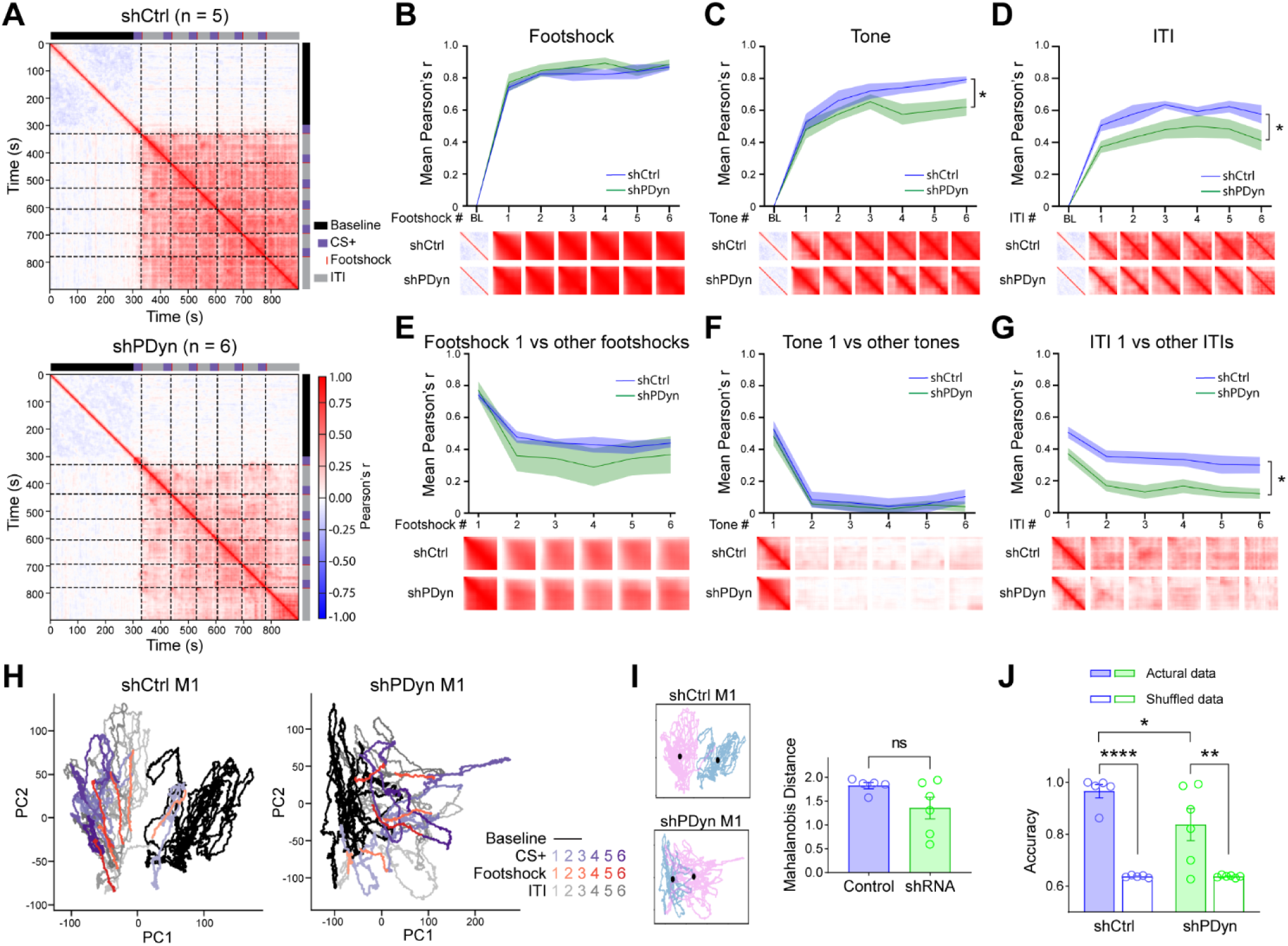
Dyn signaling promotes state transitions of vmPFC dynamics upon threat encounter. A) Correlation matrix of population activity in control and PDyn shRNA-expressing mice across all time points. Baseline (black), cue (purple), and footshock (red) are aligned to the time points on the matrix. The black dashed line is aligned to shock. B) Mean Peason’s correlation coefficient (PCC) in control and PDyn shRNA-expressing mice during the footshock periods across all trials. Insets consist of time windows used to derive the mean (PCC). C) Mean PCC during the tone period (Two-way ANOVA, Treatment Main Effect, \**p*<0.05). D) Mean PCC during the ITI (Two-way ANOVA, Treatment Main Effect, \**p*<0.05). E) Mean correlation of population dynamics between the first footshock delivery on trial 1 and footshock delivery on all subsequent trials. F) Same as above but for the cue period. G) Same as above but for the ITI period (Two-way ANOVA, Treatment Main Effect, \**p*<0.05). H) Representative neural trajectory plots from a control and a PDyn shRNA-expressing mouse. I) Manahalobis distance between the mean of the baseline period and the mean of the remaining portion of the threat conditioning session beyond the first tone. J) Results from stratified linear classifier demonstrating that the neural state space obtained from control mice has a significantly higher accuracy at correctly predicting whether the time series belonged to the baseline or post-shock epoch of the session (Two-way ANOVA with Bonferroni’s Post Hoc test, \**p*<0.05, \*\**p*<0.01, \*\*\*\**p*<0.0001).

To further explore the impact of vmPFC^Dyn^ peptidergic transmission on population-level vmPFC dynamics, we examined the trajectory of the neural state space of vmPFC networks during threat conditioning procedures in control and vmPFC^Dyn^ knockdown mice using Principal Component Analysis (PCA). In control mice, the first presentation of the tone, before it is associated with the footshock, did not cause a shift from the subspace of vmPFC activity associated with baseline exploratory behavior in the chamber (Fig. 7H). Interestingly, the trajectory of the neural state space shifted immediately away from the baseline state after the first footshock co-terminating with the first tone presentation. The neural trajectory did not, or minimally, overlapped with the baseline subspace during subsequent intertrial intervals (ITI) and tone presentations (Fig. 7H,I). This shift from baseline subspace upon threat exposure (the first footshock) was smaller in 4 out of 6 vmPFC PDyn knockdown mice, though this failed to reach significance at group level (Fig. 7H, I). However, a logistic regression model trained on randomly sampled time bins classified vmPFC subspace as belonging to baseline or post-shock periods with 96.6% accuracy in control mice (Fig. 7J). In contrast, in PDyn shRNA mice accuracy was 83.7%, which was significantly lower than control mice (Fig. 7J). Collectively, these results suggest that threats drive vmPFC networks state transitions and this state transition is impaired by vmPFC Dyn knockdown.

## Discussion

Here, we provide evidence that vmPFC^Dyn^ neurons utilize Dyn-mediated signaling to respond to threats and suppress a passive defensive behavior. We demonstrate that vmPFC^Dyn^ neurons are more responsive to threats relative to Dyn-negative mPFC neurons. Dyn is expressed in a subset of inhibitory SST-expressing neurons and excitatory neurons, which are differentially embedded within cortical circuits. Both excitatory and inhibitory vmPFC^Dyn^ neurons are recruited by threats and their predictors. Moreover, we characterized the genetically-encoded fluorescent KOR sensor, kLight, in the mPFC both *ex-vivo* and *in-vivo*. Using kLight in tandem with fiber photometry, we demonstrate that footshocks and restraint stress rapidly drive vmPFC Dyn release from locally-derived sources. PDyn knockdown in the vmPFC promotes passive defensive behaviors suggesting that Dyn-mediated signaling facilitates vmPFC circuit suppression of passive defensive behavioral states. Lastly, Dyn-mediated signaling modulates excitation and inhibition of vmPFC neurons in response to threat and facilitates transitions to threat-related vmPFC network states. Collectively, these results highlight a critical role for vmPFC^Dyn^ neurons embedded within vmPFC circuits in responding to threats and constraining passive defensive behaviors.

Dyn / KOR signaling has been implicated in mediating motivation and affective behavior (Bruchas et al., 2010, Crowley et al., 2016, Tejeda et al., 2012, Massaly et al., 2016, Cahill et al., 2022). Here, we observed that vmPFC^Dyn^ neurons, including excitatory and inhibitory mPFC^Dyn^ neurons, are more robustly activated by footshocks and footshock-predictive cues relative to the vmPFC pan-neuronal population. Moreover, using kLight we demonstrate that vmPFC^Dyn^ neuron activity results in rapid release of Dyn within vmPFC circuits in response to threats and threat-predictive cues. To our knowledge, this is the first demonstration that neuropeptide release in mPFC circuits can occur rapidly, in the order of seconds, and track the activity of cells that release it. Cue-evoked Dyn release developed rapidly with footshock-predictive cues and transients that last seconds are observed with restraint stress. These results highlight the utility of kLight as a sensor to monitor Dyn / KOR signaling on rapid time scales. kLight responses lasting almost an hour were previously documented in the mPFC during naloxone-precipitated morphine withdrawal (Abraham et al., 2021). Importantly, using the kLight sensor in combination with viral-mediated PDyn knockdown, we were able to demonstrate that vmPFC kLight responses rely on Dyn released from vmPFC neurons in response to threat predictive cues. Our results further demonstrate that vmPFC Dyn release occurs to other aversive modalities and, importantly, tracks associative learning, suggesting that Dyn signaling may not only be modulating the processing of unconditioned aversive stimuli but states of fear associated with aversive outcome anticipation. Interestingly, both excitatory and inhibitory classes of vmPFC^Dyn^ neurons decreased their activity during extinction when footshocks were omitted, and outcomes were updated, consistent with the hypothesis that Dyn neuron activity and release tracks states associated with active threats. Accordingly, vmPFC^Dyn^ knockdown potentiated passive defensive behavior (freezing) during initial threat conditioning acquisition, without interfering with long-term fear memory recall, locomotor function, working/short-term memory, and autonomic responses to threats. These results suggest that Dyn may exert its effects via specialized vmPFC circuit elements that limit expression of passive defensive behaviors that persists beyond the actual threat or its predictor. Together, these results suggest that Dyn acts within vmPFC circuits to limit fear-related behavior, which may serve to disengage from active threat interaction.

Expression of Dyn in subsets of mPFC cells provides a specific mechanism for distinct excitatory and inhibitory cell types to influence vmPFC circuits. Here, both excitatory and inhibitory vmPFC^Dyn^ neurons were excited in response to threats and threat-predictive cues, but decreased their activity rapidly once shocks were omitted. These results suggest vmPFC^Dyn^ neurons may be broadly sensitive to threat state. In primary sensory cortices, Dyn expression is restricted to SST interneurons (Sohn et al., 2014, Smith et al., 2019), whereas in the mPFC, PDyn expression is observed in both SST interneurons and excitatory neurons (present study) (Sohn et al., 2014). Putative excitatory neurons in the insular cortex are labeled in PDyn-Cre mice crossed with a reporter mouse line (Pina et al., 2020), consistent with the notion that in higher-order cortical structures Dyn may also be expressed in excitatory neurons. Expansion of Dyn expression to excitatory neurons may confer higher processing capabilities by allowing Dyn released from excitatory neurons to influence activity at synapses and somatodendritic compartments containing functional KORs which in other brain regions would only be accessible to neuromodulation by Dyn-positive interneurons. SST neurons in cortical circuits are heterogeneous and display a plethora of electrophysiological, morphological, and molecular properties (Nigro et al., 2018, Naka et al., 2019, Tasic et al., 2018, Kupferschmidt and Gordon, 2022). Dyn-negative SST neurons were enriched in more superficial layers than mPFC^Dyn/SST^ neurons, suggesting that mPFC^Dyn/SST^ neurons represent a bona fide sub-population of SST interneurons. Interestingly, vmPFC^Dyn/SST^ neurons displayed differential activity in response to threat conditioning and extinction procedures relative to Dyn-negative SST interneurons, emphasizing that vmPFC^Dyn/SST^ cells may be functionally distinct, including an increased propensity to track on-going threat state. Thus, Dyn may be utilized as a specialized neuromodulator by subsets of excitatory principal neurons and inhibitory SST interneurons to influence vmPFC networks.

Dyn signaling may exert differential effects on motivated behavior depending on the brain region this system is regulating. Here, vmPFC Dyn signaling limited passive defensive behavior, in contrast to its well-appreciated role in mediating mal-adaptive behavior via actions in other limbic regions, such as the extended amygdala (Al-Hasani et al., 2015, Shippenberg, 2009, Wigton et al., 2022). The present results are consistent with literature demonstrating that the vmPFC plays a critical role in decreasing affective and passive defensive reactions to threats. This includes the role of the vmPFC in promoting extinction of threats (Sierra-Mercado et al., 2011), generalization of fear memories (Bayer and Bertoglio, 2020), and anxiety-like behavior (Chen et al., 2021). Work from the Quirk group has demonstrated that vmPFC (infralimbic) inhibition increased freezing and suppressed reward-seeking in a conflict-based task (Bravo-Rivera et al., 2014). Moscarello and LeDoux (Moscarello and LeDoux, 2013) demonstrated that infralimbic lesions or inactivations impair avoidance behavior and promote freezing. The Dyn / KOR system may play a role in modulating vmPFC circuits and their control over defensive behaviors in response to acute threats. These results are consistent with a previous study demonstrating that KOR activation in the infralimbic cortex produces anxiolytic effects (Wall and Messier, 2002). Moreover, a translational study demonstrated that PDyn constitutive KO mice displayed impairments in contextual fear extinction and decreased extinction-induced vmPFC (infralimbic) cfos activation (Bilkei-Gorzo et al., 2012). In this study, humans with a polymorphism in the PDyn genes that presumably confers reduced PDyn expression had impaired fear extinction and coupling between the vmPFC and amygdala during fear extinction, a circuit that we have reported is regulated by endogenous Dyn (Tejeda et al., 2015, Yarur et al., 2023). Rewarding effects of endogenous Dyn release have also been observed in the dorsomedial NAcc shell (Al-Hasani et al., 2015), highlighting that Dyn signaling exerts different effects on motivated behavior depending on the brain region. These results suggest that Dyn signaling within vmPFC circuits shapes information processing to constrain persistent defensive behavior initiated by acute threats.

Our study demonstrates that cortical neuropeptidergic transmission can control shifts in cortical networks associated with abrupt shifts in behavioral state. Here, Dyn-mediated signaling is necessary for vmPFC network transitions to a “fear” state in response to threat. vmPFC Dyn knockdown prevents the activation of vmPFC cells that increase their activity not to the threat per se, but the behavioral state of fear persisting beyond the presence of the aversive footshock. vmPFC Dyn peptide release is also necessary for threat-associated state transitions in vmPFC networks that never return to basal levels. Since vmPFC^Dyn^ neurons have an increased propensity to respond to footshocks and rapidly adapted their activity when shocks were omitted, this would position this neuropeptidergic system to consistently release peptide with active on-going threats and predictors of threat. This Dyn release and subsequent KOR activation is, likely in part, responsible for shifting the activity of vmPFC ensembles and networks associated with a state of “on-going, active” threat during time periods where persistence of a defensive behavior must be regulated (e.g. the ITI). Slower Dyn-mediated signaling, delimited by Dyn volume transmission and long-term modifications in synaptic transmission after KOR activation (Yarur et al., 2023), may provide a mechanism for threats and their predictors to generate a brain state that persists even after the acute threatening stimuli are absent (e.g. during the ITI where Dyn knockdown effects on behavior and network dynamics are largest). It should be noted that viral mediated vmPFC^Dyn^ knockdown does not permit us to conclusively determine whether vmPFC Dyn / KOR transmission is regulating vmPFC state transitions and/or through an indirect circuit mechanism, such as regulation of KOR-expressing thalamo-cortical loops at the level of the thalamus (Yarur et al., 2023). Independent of its exact mechanism, by facilitating engagement of vmPFC during active threat exposure, Dyn signaling may be driving the suppression of defensive behaviors (Roelofs and Dayan, 2022, Alexandra Kredlow et al., 2022). vmPFC activity may operate in a push-pull manner, in conjunction with other circuits and/or cell types that suppress passive defensive behaviors, to oppose circuits and/or cell-types that promote passive defensive behaviors. Thus, vmPFC Dyn signaling may act locally to influence how behavioral states driven by threats are dynamically toggled through the various phases of threat evaluation, encounter, escape, and/or termination (Roelofs and Dayan, 2022, Moscarello and Penzo, 2022, Fanselow, 2018, Mobbs et al., 2020).

It is currently unclear how the Dyn / KOR system may be shaping mPFC circuit function to modify behavior. KOR activation inhibits inputs to the PFC via a presynaptic site of action, including those arising from the BLA, VTA, and PVT (Tejeda et al., 2013, Tejeda et al., 2015, Yarur et al., 2023). Incoming inputs from the VTA, BLA, and PVT are critical for various behaviors, including the processing of aversive events and extinction (Marcus et al., 2020, Senn et al., 2014, Burgos-Robles et al., 2017, Cummings and Clem, 2020, Vander Weele et al., 2018, Gao et al., 2020). Thus, Dyn release acting on KORs on these presynaptic terminals from afferent inputs may regulate extrinsic factors that control vmPFC circuit function. KOR mRNA expression is also observed within mPFC circuits (Meng et al., 1993, Peckys and Hurd, 2001, Yarur et al., 2023), and Dyn inhibits the release of glutamate and GABA from KOR-positive excitatory and inhibitory cells (Yarur et al., 2023). KOR-positive mPFC cells may respond to Dyn released from mPFC^Dyn^ neurons via KOR-containing local circuit axons, soma, and/or dendrites. Future work dissecting the role of Dyn / KOR signaling in filtering incoming afferents and the role of KOR expressing cells locally in shaping information processing will be of importance in dissecting how this system participates in motivationally-charged behaviors.

In conclusion, we provide evidence of threat-induced vmPFC^Dyn^ neuron activation and subsequent Dyn release in the vmPFC. vmPFC Dyn signaling within local circuits constrain passive defensive behavior in the face of active threats. This occurs via reorganization of vmPFC network states associated with fear that may facilitate the inhibitory function of the vmPFC in suppressing threat responsivity. The rapid release of PDyn-derived peptides, revealed by the employment of a genetically-encoded opioid sensor, and the impact of Dyn combination of neuropeptidergic manipulation on network states significantly advance our understanding of the capacity of neuropeptides in shaping rapid, dynamic cortical processing during behavior. Future work aimed at understanding whether adaptations in Dyn / KOR signaling in mPFC circuits mediate mal-adaptive behavior may elucidate novel therapeutic strategies to treat neuropsychiatric disorders.

## Extended Methods

### Animals

Male and female C57BL/6J WT, prodynorphin-Cre (PDyn-Cre), somatostatin-Flpo (SST-Flpo) were used. Mice were housed in humidity and temperature-controlled vivariums using a reverse light cycle with lights off at 8:00 hours and lights on at 20:00 hours. Animals had ad-libitum access to standard laboratory chow and water. All procedures were approved by the National Institute of Mental Health and National Institute of Dental and Craniofacial Research Animal Care and Use Committees.

### RNAscope in-situ hybridization

RNAscope procedures were conducted as previously described (Tejeda et al., 2017, Tejeda et al., 2013, Wang et al., 2012). For all RNAscope procedures, tools, and equipment were cleaned with 70% ethanol and RNAse inhibitors (RNaseZap, Invitrogen). Mice were euthanized by cervical dislocation; brains were rapidly dissected, and flash-frozen with 2-methylbutane chilled on dry ice for 20 sec. Subsequently, brains were stored at −80°C until sectioning. Brains were sliced utilizing a cryostat (Leica), and 16-18 µm sections were mounted on Superfrost Plus slides (Fisher Scientific). Slides containing sections were stored at −80°C until in-situ hybridization processing. For human post-mortem DLPFC and sgACC tissue, samples were collected from the NIMH Human Brain Collection Core.

RNAscope procedures were conducted according to the Advanced Cell Diagnostics (ACD) user manual. Slides were fixed with 4% paraformaldehyde for 20 min at 4°C. Slides were subsequently washed twice for 1 min with PBS, before dehydration with 50% ethanol (1 x 5 min), 70% ethanol (1 x 5 min), and 100% ethanol (2 x 5 min).

Slides were incubated in 100% ethanol at −20°C overnight. The next day, slides were dried at room temperature for 10 min. A hydrophobic pen was used to draw a hydrophobic barrier around the sections and allowed to dry for 10-15 min. Sections were then incubated with Protease Pretreat-4 solution for 20 min at RT. Slides were washed with PBS (2 x 1 min), before being incubated with the appropriate probes for 2 hr at 40°C in the HybEZ oven (Advanced Cell Diagnostics). Mouse mPFC sections were processed using RNAscope Fluorescent Multiplex Reagent Kit (320850, ACD). Probes used were: Mm-Pdyn (318771), Mm-Slc17a7-C2 (416631-C2), Mm-Slc32a1-C3 (319191-C3), Mm-Sst-C2 (404631-C2), and Mm-Pvalb-C3 (421931-C3). Human dmPFC sections were processed using RNAscope Multiplex Fluorescent Kit v2 (323120, ACD). Probes used were: Hs-PDYN (507161), Hs-SLC17A7-C2 (415611-C2), and Hs-SLC32A1-C3 (415681-C3).

Slides were scanned using Nikon C2 or Zeiss 780 confocal microscope using 20X objective. Images were processed with Fiji and RNAscope signals were quantified with CellProfiler 3.1.8 (McQuin et al., 2018, Erben and Buonanno, 2019). Histograms of SST mRNA-positive PDyn mRNA-positive cells relative to PDyn mRNA-negative cells were plotted by binning (25 µm bin) the depth of the cells from the midline.

### Surgeries

For Stereotaxic surgery, mice (8-12 weeks) were anesthetized with the cocktail of ketamine (100 mg/Kg) and xylazine (10 mg/kg). Viruses were bilaterally or ipsilaterally injected by a 32G metal needle connected to a hamilton syringe which mounted on a microinjection pump (UMP3, WPI). The coordinates for viral injections are: vmPFC, +1.7mm A/P, ±0.3mm M/L, −2.6 D/V; Volumes of virus ranges between 200-400 nl / site. For optic fiber implantation, a 400 μm fiber-optic cannula (NA 0.66, B280-4604-4.1, Doric Lenses) was placed 100 μm above the center of viral injection site, and cemented onto the skull using Metabound (C&B Metabond, Parkell). For gradient index (GRIN) lens implantation, one week after GCaMP viral injection a (GRIN) lens (0.6 × 7 mm or 1 × 4mm, Inscopix) was implanted in the mouse brain above vmPFC under isoflurane anesthesia. In brief, a craniotomy slightly larger than the GRIN lens diameter was made (A/P:+1.7mm, M/L:0.4mm). The brain tissue above the mPFC was removed by aspiration via a 30-Gauge blunt needle connected to the lab vacuum. GRIN lens was slowly (100 μm/min) lowered through the craniotomy until it reached −2.5mm D/V. The GRIN lens was then secured to skull using Metabound. Two weeks after GRIN lens implant, a baseplate was mounted on top of the lens. The location of baseplate was determined with miniscope camera attached, first focused on the edge of the GRIN lens, then moved baseplate up 300-350 μm and secured to the skull with Metabound. For PDyn shRNA / miniscope Ca2+ imaging experiments, integrated GRIN lenses with baseplates were used. For pupillometry recording, after viral injection a head bar 22.3 mm long and 3mm wide was fixed to the back of the head using metabond and covered with dental acrylic.

### Behavior

#### Threat conditioning

Mice were placed in threat (fear) conditioning chambers housed in sound-attenuating chambers (Coulbourn Instruments). During conditioning, auditory cues (5 kHz; 30 sec duration) co-terminating with a 0.6 mA, 2 sec footshock delivered through grid floors (Coulbourn Instruments). Mice were exposed to a total of 6 or 10 tone footshock pairings for one or 3 days for behavioral and in-vivo recording studies. During cued-fear recall and extinction, mice were placed in a novel context (context B) containing a plexiglass floor, distinct visual cues, chamber geometry, and odor of 5% acetic acid. Mice were exposed to the conditioned auditory cue 15-20 times during these sessions to determine cued-fear recall during early trials and threat extinction during later trials. During cue renewal, mice were placed in the same context (context A) as conditioning days. Mice were exposed to conditioned auditory cues 5 times without any footshock. Behavior was monitored and analyzed using FreezeFrame 4 (Actimetrics) behavioral tracking software.

Mice underwent cue discrimination testing were similar to procedures described by the Zweifel group (Fellinger et al., 2021, Baird et al., 2021). Mice were placed in threat (fear) conditioning chambers housed in sound-attenuating chambers (MedAssociates). Equipment was controlled using AnyMaze (Stoelting). On the first day of the task, mice were first exposed to context A consisting of solid floors, hexagonal steel walls, with 5% acetic acid odor. In context A, mice were exposed to two neutral, 10 second cues (4 and 12 kHz). For baseline, the CS+ and the CS-were presented three times each. During acquisition, mice are placed in a novel context consisting of shock grids, a square chamber with two stainless steel walls and two Plexiglass walls. During acquisition, the 4Hz auditory cue (CS+) was presented and co-terminated with a 0.5 sec footshock (0.4 mA). Trials where mice were presented with the 12kHz cue (CS-), which did not co-terminate with the footshock, were randomly interleaved. During acquisition, the CS+ and the CS-were presented ten times each, and the first cue was the CS+. Mice were subsequently tested for cued fear retrieval in context A where the CS+ and the CS-were presented three times each. All tests began with a 2-minute habituation. The ITI between trials was 60 sec. Behavior was monitored and analyzed using AnyMaze (Stoelting) behavioral tracking software.

#### Elevated Plus Maze

The elevated plus maze is a cross-shaped arena consisting of two open arms (33.02 L x 5.08 W cm) and two closed arms (33.02 L x 5.08 W x 15.24 H cm) that was elevated 62.23 cm above the ground. Mice were allowed to explore the maze for 10 min. Behavior was monitored and analyzed using Topscan behavioral tracking software.

#### Light/Dark Box

The light-dark box consisted of an open-lit compartment (43.815 L x 21.9 W cm) and a closed dark compartment (43.815 L x 21.9 W cm). The two compartments were separated by an entry (4 W x 5 H cm) in the middle. Mice were placed in the arena, and the latency to enter the dark and time spent in the light and dark were recorded. Behavior was monitored and analyzed using Topscan behavioral tracking software.

#### Novel Object Recognition Test

The novel object recognition test was conducted in a 43.815 L x 43.815 W cm chamber. During the baseline period, two identical objects (object 1, Legos) were placed in the chamber and time spent with both objects was monitored for 10 min. Mice were removed from the chamber for 60 min. Mice were reintroduced to the chamber that now contained a novel object (object 2) and the same object (object 1) used during baseline for another 10 min. Time of mice spent observing and sniffing on each object were analyzed using Topscan behavioral tracking software.

Difference index was calculated as: (Time with object 2 – Time with object 1) / (Time with object 2 + Time with object 1).

#### Spontaneous Alternation Task (Y-maze)

Working memory was assessed utilizing a Y-shaped maze with arms 55.88 L x 10.16 L x 31.75 H cm extending from the maze’s center. Mice were allowed to freely explore the three arms for 10 min. Number of arm entries and number of alternations are scored using Topscan behavioral tracking software. Alternation was counted when the mouse visited all three arms in three consecutive arm entries. The alternation percentage is calculated by dividing the number of alterations by the number of arm entries x 100.

#### Restrained Stress

Mice were subjected to restrained stress by placement in 50-mL conical tubes for 30min. Holes were drilled at the front of the tube to allow proper ventilation during the restrain. The tube has a 5mm opening slot to allow fiber optic cannula connections to patch cord during the restrained stress.

#### Pupillometry

Head-fixed pupillometry was recorded as described by Gao et al 2020 (Gao et al., 2020). Briefly, mice were first acclimated to a custom-built head fixation device for 5 days, 30 min per day. A monochromatic CMOS camera equipped with a macro zoom lens (MVL7000, ThorLabs) and an infrared LED lamp (LIU850A, ThorLabs) were used to collect images from the left pupil at 5 frames per second. The start of camera recording and the delivery of mild aversive stimulus were triggered by TTL signal controlled by Clampex. 5 consecutive air puffs (100 ms, 60 psi) or tail shocks (1s, ∼250 μA) were started 5 min after the baseline recording, with the interval between stimulation range from 30-60s. The 29G tubing for air puff was placed 2mm under the tip of the snout so that the air puff won’t cause eye blink of the mouse. The recorded videos were subsequently analyzed using Bonsai 2.4, and changes in pupil size (area) were plotted.

#### Hargreaves test

Mice were placed in plexiglass testing enclosures on an elevated transparent platform for at least 60 min prior to testing. The plantar surfaces of the hindpaws were stimulated using a movable IR heat source (Ugo Basile), and the time taken for the withdrawal of the stimulated paw was recorded. The heat intensity of the heat source was set to 30% and a cutoff time of 20s was used to prevent tissue damage. At least 5 min was given between stimulating the same paw again to allow for recovery. The surface of the transparent platform was kept clean to ensure that the heat from the IR heat source was not perturbed. Four to six measurements were taken from each hindpaw and an average value was reported as the withdrawal latency.

#### von Frey test

Mice were placed in plexiglass enclosures on a wire mesh frame and were allowed to acclimate for at least 60 min prior to experiments. The plantar surfaces of the hindpaws were stimulated with calibrated von Frey filaments starting from a filament weight of 0.4 g. Filaments were gently pressed against the hind paw plantar surface until the filament bent slightly, and was held in place for 3-5s. For each trial mice were tested with the same filament 3-5 times, and the number of paw withdrawals was recorded. The weight of the next filament was heavier or lighter than the previous, with higher weight filaments chosen when <50% of responses failed to produce a paw withdrawal, and were lower when >50% of trials resulted in a withdrawal. This process was repeated five times to determine the 50% withdrawal threshold.

#### Hot plate test

Mice were placed in a plexiglass container on a heated plate set to 52°C (IITC Life Science, PE34), and the time taken for the first nocifensive response (flicking, shaking, or licking the paw) was measured. In addition, the time taken for escape-like behaviors was also recorded (time to first jump). Mice were removed from the hot plate either after the first escape attempt, or after 60 s if no attempts were made to escape. An average result from 3-4 trials performed on separate days is reported.

For jumping behavior, it was noted that few mice attempted to escape the hotplate during the first trial, but subsequent exposures resulted in more consistent escaping behaviors. Therefore, the first trial was not included in the calculation for escape behavior.

#### CFA inflammation

To induce paw inflammation, the glabrous surface of the left hind paw was injected with 20 µl of Complete Freund’s Adjuvant (CFA) (Sigma F5881). The needle was left in place for over 5 s to reduce the leak from the injection site. Sensory testing was performed between 24- and 48-hours post-CFA injection.

### Fiber Photometry

*In-vivo* fiber photometry experiments were conducted as previously described (Pignatelli et al., 2021). For recordings, light-emitting diodes (LEDs) were modulated by DAC ports on a real-time signal processor (RZ5P; Tucker Davis Technologies) that controlled a light-emitting diode LED driver (Thorlabs) connected to 405 nm (isosbestic control; modulated at 530 Hz) and 470 nm (calcium-dependent GCaMP7 or Dyn-dependent kLight signal; modulated at 211 Hz). Light was fed into a fluorescent minicube (Doric Lenses) connected to a femtowatt photoreceiver (Model 2151; Newport) or a minicube with integrated detectors and lock-in amplifiers (Doric Lenses). Signals were then collected, digitized at 6 kHz, and recorded using the RZ5P processor. Analysis of recorded signals was performed using custom-written Python scripts. Signals in the 405 nm channel were corrected for drift by calculating the linear regression of the signal and subsequently subtracted from GCaMP7 or kLight signals to remove bleaching and artifacts introduced by fiber bending and/or motion. Full session GCaMP7 or kLight signals were normalized to the baseline period during behavioral testing prior to the presentation of any stimuli. Peri-event signals were normalized to the 20 sec window preceding the onset of the auditory cue.

### Miniscope recording

Single cell Ca^2+^ recordings through a GRIN lens with a head-mounted miniaturized microscope (nVoke, Inscopix) were conducted 1-2 weeks after baseplate implant in the data presented in Figure 1 and approximately one month after implantation with an integrated GRIN lens for data presented in figures 6 and 7. Each mouse was given 3 days of habituation, in which the miniscope camera was attached to the baseplate and mice were put in an open field for 20 min. During the habituation, miniscope settings (frequency, LED power, gain, and focus position) for each mouse were determined, and these settings were kept for each mouse throughout the behavioral recording.

To identify individual regions of interest (ROIs) Ca^2+^ dynamics from raw miniscope recordings, imaging data were first processed using the Inscopix Data Processing Software (IDPS, Inscopix). Videos were spatially downsized by a factor of 4 and temporally downsampled to 10 Hz (only for experiments in figures 6 and 7), bandpass filtered (0.005-0.5 pixels) and motion corrected. Neuronal signals from miniscope imaging data were extracted using constrained nonnegative matrix factorization-extended (CNMFe) (Zhou et al., 2018) using the following parameters: spatial down sample factor = 1, temporal down sample = 1, Gaussian filter size = 2, maximum neuron diameter = 5, minCorr=.8, min pnr=10, merge threshold = 0.8, ring size factor = 1.4, patch size = 80, patch overlap = 20). After CNMFE, probable neurons were visually inspected by experimenters for soma-like morphology with an appropriate size and shape as well as dynamic activity consistent with that of Ca^2+^ dynamics. Cells with abnormal morphology or Ca^2+^ dynamic activity were rejected from the final cell set. The data were then exported to CSV files and custom-written Python scripts were used from there onwards.

### Multiple Linear Regression (MLR) with Time Kernel

To determine how the activity of single cells in the vmPFC were influenced during threat conditioning procedures and changes in locomotor activity we employed a multiple linear regression model.

Each neuron was fitted to the the following MLR model:

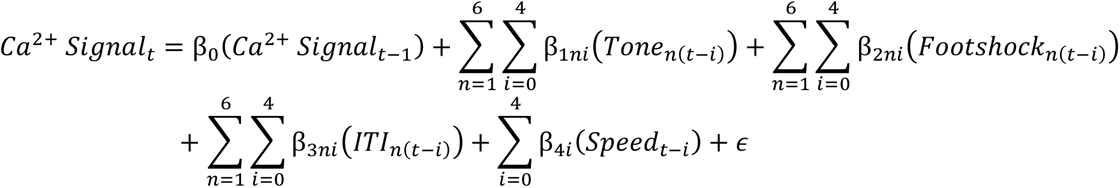

The dependent variable in this model was the ΔF/F_noise_ value (*Ca*^2+^ *Signal_t_*) of each neuron, and the independent variables were the task-related variables and speed and their interaction with a time kernel. The data were aggregated into 500 ms bins, and the time kernel consisted of 5 time bins (i), equivalent to 2.5 seconds. This allowed us to investigate how past stimuli or events influenced current neural activity, which is crucial for understanding encoding of information by individual neurons. β_0_(*Ca*^2+^ *Signal_t_*_-1_) is an autoregressive term to account for the influence of past calcium signals on the current events. This term included the calcium signal lagged by one time bin (500 ms). Each β is the coefficient of this model capturing the strength and direction of the relationship between each task variable and the neuronal activity. *Tone_n_*, *Shock_n_*, and *ITI_n_*are stimuli/task phases on trial n.

To analyze the encoding of stimuli/task phases (tone, shock, ITI) on a trial-by-trial basis, a design matrix was constructed with dummy-coded variables to represent the presence or absence of each stimulus. Each stimulus was allocated as a separate column, with time points as rows. Specifically, when the tone, shock, or ITI was present during a particular time point, it was coded as 1; conversely, when these stimuli were absent during that time point, it was coded as 0. Speed of mice was treated as a continuous variable and was not subdivided into individual trials.

To assess the significance of variables in influencing neural activity, we compared the variance explained by the full model, which included the variable of interest, to that of a reduced model that excluded a specific variable, allowing us to test whether the difference in explained variance was statistically significant (p<0.05). A variable that significantly modulated neural activity was considered “encoded” in the neural signal of a given neuron (Costa and Averbeck, 2015, Engelhard et al., 2019).

We employed diagnostic analyses to assess potential multicollinearity issues in our regression model, including assessing the Variance Inflation Factor (VIF), where values exceeding 10 suggest the presence of multicollinearity, and the Condition Index, where values greater than 30 indicate multicollinearity. A minimum eigenvalue plot that incorporates the newly added predictors to gain insights into how the multicollinearity changes as more predictors are introduced was also examined. If the minimum eigenvalues start to drop significantly, indicating increasing multicollinearity, reconsideration of choice of predictors are necessary. Based on the diagnostic analyses we chose speed instead of freezing in our predictor because freezing as an independent variable caused very high multicollinearity in the model.

### Population-level analyses of single-cell Ca^2+^ imaging

For Ca^2+^ transients from the experiment comparing scrambled ShRNA vs PDyn ShRNA, the whole session recording of each cell was standardized (Z-Scored) relative to the first 5-minutes (baseline test period) of the recording. We analyzed dynamic changes at the population level (ΔF/F_noise_) using Principal Component Analysis (PCA) to generate the trajectory of the network through the neural state space for each mouse. First, we derived the ΔF/F_noise_ of all neurons at each sampled time point. The matrix dimensions consisted of number of time points x number of neurons. The first and second components were plotted for each time point and color coded according to epoch during the session (i.e., baseline, cue, shock, intertrial interval). To quantify the change in neural activity induced during our fear conditioning procedure, we first labeled each mouse’s PC values at each timestep as being either “baseline” or “post-baseline”. Baseline was defined as the set of all time points up to, but not including, the first shock. Post-baseline was defined as the set of all time points from the onset of the first shock until the end of the task. We then identified the centroids of derived PC coordinates for baseline and post-baseline periods and computed the Mahalanobis distance between them as follows:

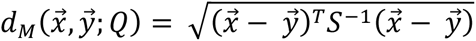

where 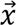 and 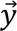 designate the centroids at baseline and post-baseline, respectively; *Q* represents the probability distribution of 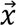 and 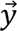 in *R^N^*; and *S*^−1^ designates the inverse covariance matrix. To determine whether the state space shift caused by threat onset was significantly different between control and PDyn shRNA groups, we trained a stratified 10-fold logistic regression model (StratifiedKFold, SciKitLearn) (Pedregosa et al., 2011) on 75% of the shuffled dataset during the baseline session or post-shock epoch of the session (all time points beyond baseline) to classify the given PC1 and PC2 coordinates as belonging to the baseline or post-shock period. The test dataset (25% of the data) was used to determine the accuracy of the model in correctly assigning new pairs of PC1 and PC2 coordinates for random time points to either the baseline or post-shock period. For each animal this process was iterated 10-fold and the mean of all iterations within a given subject was computed.

To monitor correlated activity across different behavioral epochs we generated a correlation matrix where each n-dimensional (e.g. number of neurons) population vector at time t was correlated (mean Pearson’s correlation coefficient (PCC)) with the population vector of the same neurons at every other time point within the session, yielding a t x t matrix which represents the “moment by moment” changes in correlated activity across the population throughout the entire session. We then calculated the mean PCC within or across specific trial epochs of defined windows (28 sec of the cue, 5 sec during the shock and during the ITI). Because the PCC matrix is symmetric and contains values of 1 along the diagonal, we factored out the diagonal from our computations of the mean PCC at each window.

### PDyn shRNA

pAV-U6-scrambled shRNA-GFP and pAV-U6-PDyn shRNA-tdTomato cloning and AAV packaging were conducted by Vigene Biosciences using the following sequence, GGTTGCTTTGGAAGAAGGCTACA. AAV-U6-PDyn shRNA-GFP was injected into the mPFC, and knockdown of the virus was verified utilizing PDyn immunohistochemistry after kLight fiber photometry recordings.

### PDyn immunohistochemistry

Mice were perfused with PBS and 4% PFA after behavioral testing. 30 μm mPFC sections were collected using a cryostat. Sections were incubated in blocking buffer (4% BSA + 0.3% Triton X-100 in PBS) for 1 hour at room temperature then transferred into primary antibody buffer (Anti-ProDynorphin antibody, ab10280, Abcam; 1:200 dilution in blocking buffer). Sections were incubated in primary antibody for 48 hours at 4 °C, then rinsed in PBS and transferred to secondary antibody (Alexa Fluor 594 AffiniPure Donkey Anti-Guinea Pig IgG, 706-585-148, Jackson ImmunoResearch; 1:200 dilution in blocking butter) for 2 hours at room temperature. Sections were then rinsed in PBS and mounted on slides. Fluorescence images were taken on Zeiss 780 confocal microscope with 20X objective. Data were quantified using Fiji (ImageJ) by analyzing fluorescence intensity in fixed-size regions of interest with intense GFP expression.

### Acute brain slice preparation

Mice were anesthetized with euthanasia (NIH Veterinarian Services) and subsequently decapitated. Brains were removed rapidly and placed in ice-cold NMDG-based cutting solution (Ting et al., 2014) containing (in mM): 92 NMDG, 20 HEPES, 25 glucose, 30 NaHCO_3_, 2.5 KCl, 1.2 NaPO_4_ saturated with 95% O_2_/5% CO_2_ with an osmolarity of 303-306 mOsm (Wescorp). The brain was rapidly blocked, dried on filter paper, and glued to a platform containing ice-cold NMDG-based cutting solution within a Leica VT1200 Vibratome. Coronal slices, 250 µm thick, containing the mPFC were cut at a speed of 0.07 mm/s. Following slicing, sections were incubated in a chamber containing NMDG-based cutting solution for 5-10 min at 34°C. Slices were subsequently transferred to a chamber filled with modified holding aCSF saturated with 95% O_2_/5% CO_2_ containing (in mM): 92 NaCl, 20 HEPES, 25 glucose, 30 NaHCO_3_, 2.5 KCl, 1.2 NaPO_4_ (303-306 mOsm) at room temperature for at least 1 hr. Slices remained in this solution until being transferred to the recording chamber.

### Ex-vivo whole cell electrophysiology

Whole-cell patch-clamp electrophysiology studies were performed as previously described (Tejeda et al., 2017, Pignatelli et al., 2021, Yarur et al., 2023). The recording chamber was perfused by a pump (World Precision Instruments) at a flow rate of 1.5-2.0 ml per minute with aCSF containing (in mM): 126 NaCl, 2.5 KCl, 1.4 NaH_2_PO_4_, 1.2 MgCl_2_, 2.4 CaCl_2_, 25 NaHCO_3_, and 11 glucose (303-305 mOsm). For whole-cell recordings of intrinsic excitability, we utilized glass microelectrodes (3-5 MΩ) containing (in mM): 135 K-gluconate, 10 HEPES, 4 KCl, 4 Mg-ATP, and 0.3 Na-GTP and in some cases 0.1% Biocytin. For oEPSC and oIPSC recording, we utilized glass microelectrodes (3-5 MΩ) containing (in mM): 117 cesium methanesulfonate, 20 HEPES, 0.4 EGTA, 2.8 NaCl, 5 TEA-Cl, 4 Mg-ATP, 0.4 Na-GTP, and 5 QX-314 (280-285 mOsm). Monosynaptic oEPSCs and oIPSCs evoked by Dyn+ cell stimulation were isolated with TTX (1 µM) and 4-AP (50 µM). For monosynaptic oEPSCs and oIPSCs, layer V pyramidal neurons were targeted for recordings. oEPSC and oIPSC charge transfer (pC) was calculated as the area under the curve (pA/ms). Cells were visualized using IR-DIC optics on an inverted Olympus BX5iWI microscope. Neurons were recorded utilizing a Multiclamp 700B amplifier (Molecular Devices). Data were filtered at 2 kHz and digitized at 20 kHz using a 1440A Digidata Digitizer (Molecular Devices). Series-resistance (10-20 MΩ) was monitored using a −5 mV voltage step. Cells with >20% change in series resistance were discarded from further analysis. Intrinsic excitability was assessed by applying hyperpolarizing and depolarizing current steps (25 pA steps; 1-sec duration) and measuring the change in voltage and action potential firing. Following electrophysiological recording, the pipette was slowly retracted from the cell, and the slice was transferred to 4% paraformaldehyde overnight.

### Morphological reconstruction of electrophysiologically-recorded Dyn^+^ neurons

Slices containing biocytin-filled Dyn+ neurons were washed with PBS three times for ten minutes before incubation with PBS containing 4% BSA and 0.3% Triton-X 100 for two hours. Slices were then incubated with Alexa Fluor 594 conjugated streptavidin (1:500, S11227, Thermo Fisher Scientific) in PBS containing 4% BSA and 0.3% Triton-X 100 overnight. Slices were washed with PBS at least three times for ten minutes each, coverslipped with Fluoromount G mounting media containing DAPI (Southern Biotech) and stored at 4°C in a light-protective box. High-magnification 40X tiled images (1 µm stack size) of biocytin-filled Dyn+ neurons were acquired using a confocal microscope (Nikon A1R). Morphology of biocytin-filled Dyn+ neurons was reconstructed, and morphological data were extracted from images using Neurolucida 360 (MBF Bioscience).

### kLight slice imaging

Mice were injected with AAV1-hSyn-kLight 1.2a. Imaging was performed on BX5Wi epifluorescent microscope utilizing a 0.8 NA 40X objective (Olympus), equipped with bandpass dichroic filters to direct excitation 470 nm light (50-100 µW at the focal plane of the objective) at the brain slice and collect fluorescence time-lapse images utilizing a Q-Imaging camera and Ocular Software (Q-Imaging). kLight fluorescence was evoked utilizing 50 ms light pulses delivered at 4 Hz with a fixed camera exposure time of 100 ms. Movies were analyzed using ImageJ. Integrated intensity across frames was derived and normalized to baseline prior to drug application.

#### Drugs

Drugs were dissolved in saline or water. Drugs were purchased from Sigma Aldrich, Tocris, or generously provided by the NIDA Drug Supply Program.

#### Statistics

Statistics were computed using Graphpad Prism. Detailed statistics including t- and F statistics and degrees of freedom can be found in supplemental Table 1, while *p*-values associated with symbols are noted in the figure legends. Data are presented as mean ± s.e.m. Code used in the manuscript is available upon reasonable request. The datasets generated during and/or analyzed during the current study are available from the corresponding author on reasonable request.

## Supporting information

Statistics Table

Key Resources Table

## Acknowledgements

This work was supported by the National Institute of Mental Health Intramural Research Program (ZIA MH002970-04), a Brain and Behavior Research Foundation NARSAD Young Investigator Award to HAT, NIH Center for Compulsive Behavior Fellowships to RFG and HEY, and a NIH Post-Doctoral Research Associate Training Fellowship to RFG. We would like to thank Dr. Candice Contet (Scripps Research Institute) for advice on the development of the PDyn-shRNA virus. The authors would like to thank Drs. Ted Usdin, Sarah Williams and Jonathan Kuo of the NIMH Systems Neuroscience Imaging Resource and Dr. Samer Hattar for their microscopy support. The authors would like to thank the NIMH Dr. Yogita Chudasama and members of the Rodent Behavioral Core for behavioral equipment support. Drs. Barbara Lipska and Stefano Marenco and members of the NIMH Human Brain Collection Core for providing post-mortem samples. We would like to thank members of the Unit on Neuromodulation and Synaptic Integration and Drs. Mario Penzo and Michael Krashes for their inputs. We would like to thank Drs. Yogita Chudasama and Claire Le Pichon for generously sharing their cryostats. We would like to thank Dr. Bruno Averbeck for helping our laboratory develop and implement the multivariate linear regression model for calcium imaging analysis.

---

**Figure S1:**
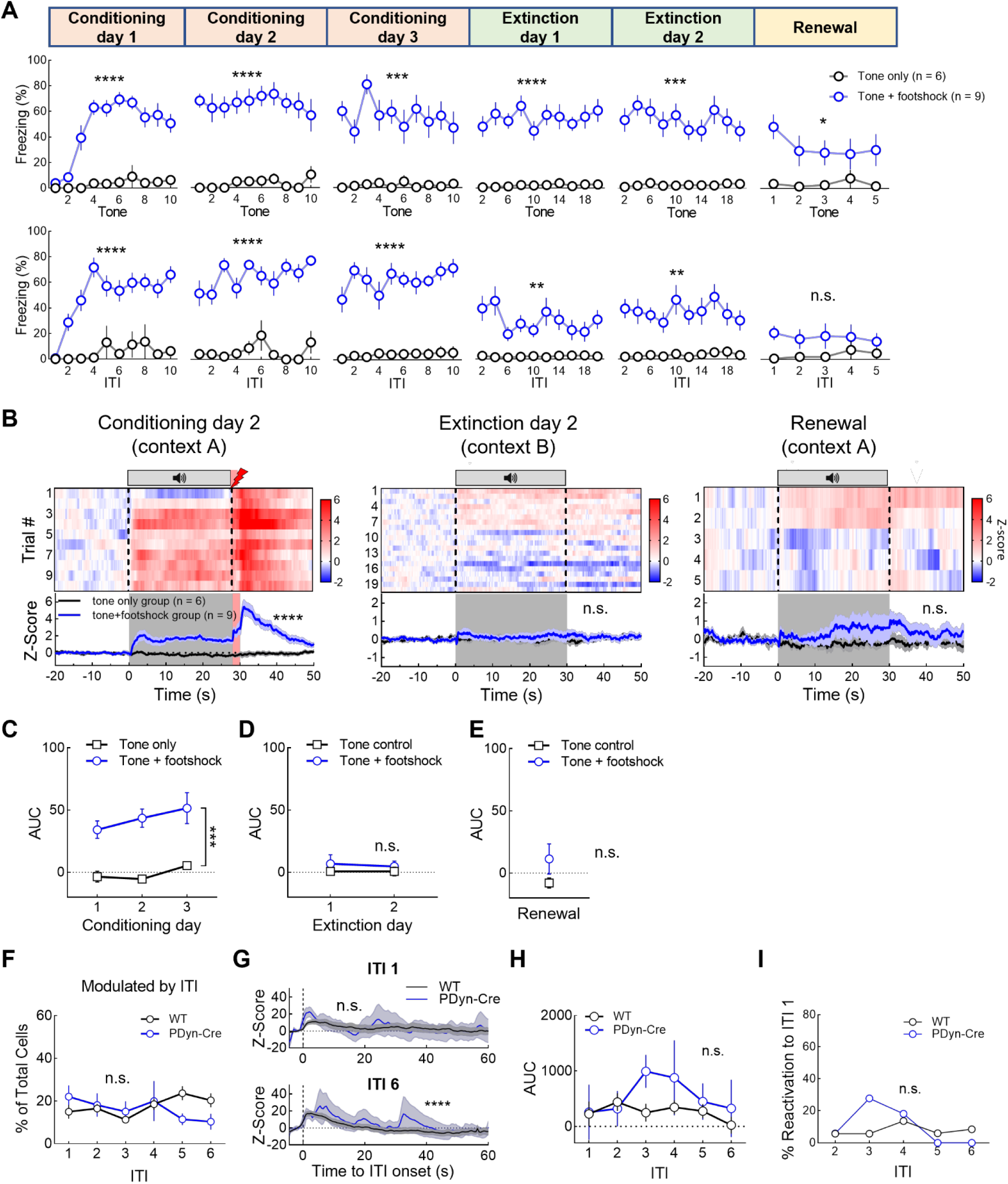
Relevant to Figure 1 In-vivo monitoring of mPFC^Dyn^ cells during fear conditioning. A) Freezing during tone and ITI period across threat conditioning, threat recall/extinction day 1 and extinction day 2, and renewal. Data is binned every 2 trials for extinction days (See Statistics Table for statistics). B) Heatmaps of fiber photometry GCaMP activity across the session during day 2 of threat conditioning, day 2 of threat extinction, and renewal of threat responsivity in the training context after extinction (RM two-way ANOVA, group main effect ****p<0.0001). C-E) AUC analysis of Ca^2+^ activity evoked during conditioning (C), extinction (D), and renewal (E) (Two-way ANOVA for C and D, group main effect, \*\*\**p*=0.0006; unpaired t-test for E). F) Percentage of neurons significantly modulated by the intertrial interval (Two-way ANOVA, cell type main effect, *p*=0.6202). G) Ca^2+^ responses aligned to ITI onset in intertrial interval-modulated neurons (Two-way ANOVA, cell type main effect, \*\*\*\**p*<0.0001). H) AUC of Ca^2+^ responses during the intertrial intervals (Two-way ANOVA, cell type main effect, *p*=0.0734). I) The percentage of footshock encoding neurons during intertrial interval 1 that were also significantly activated during all subsequent intertrial intervals (paired t-test, *p*=0.6738).

**Figure S2:**
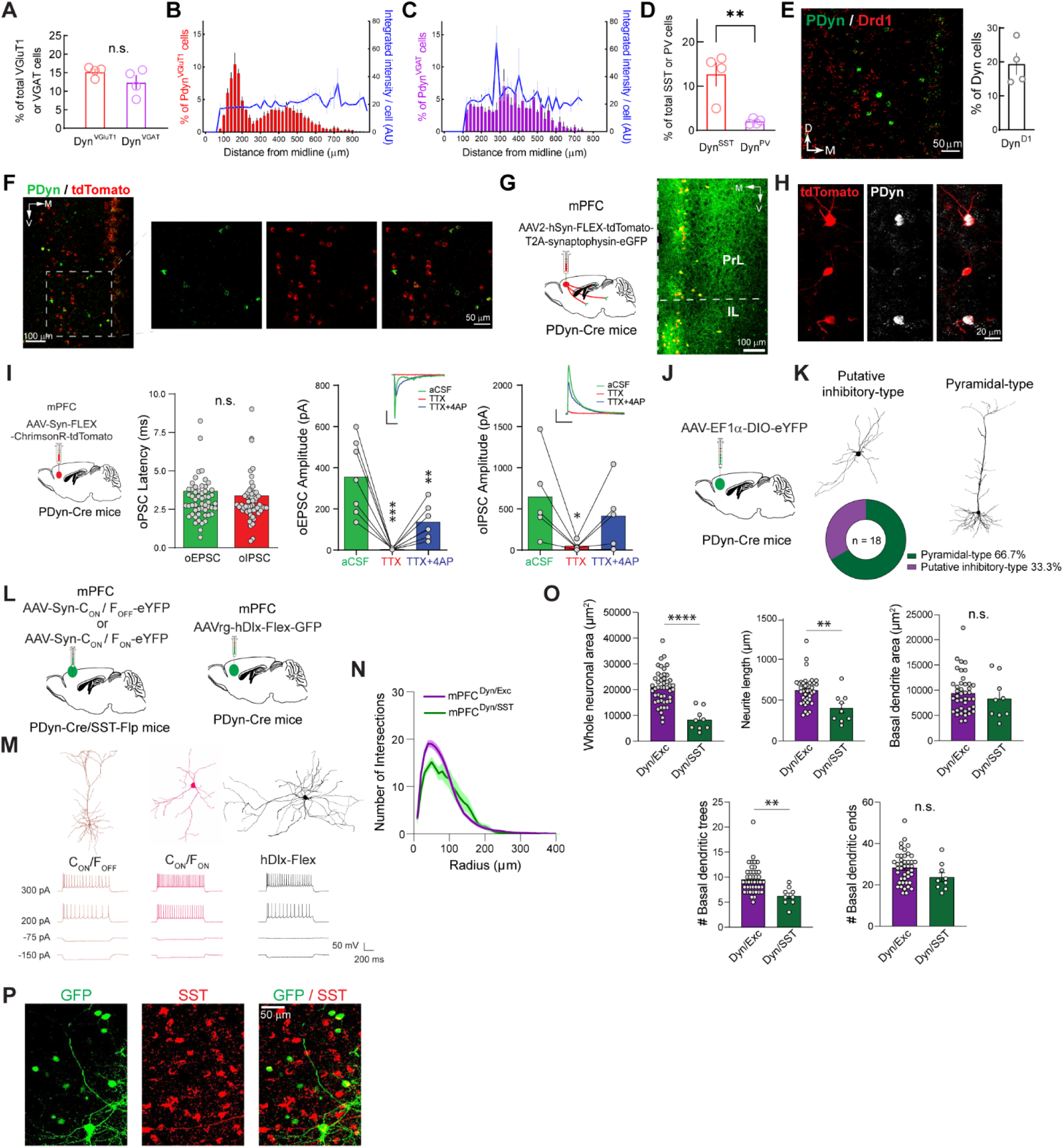
Relevant to Figure 2 Anatomical characterization of mPFC dynorphin-expressing neurons. A) The proportion of PDyn mRNA-containing neurons does not differ between mPFC excitatory and inhibitory neuron populations (unpaired t-test, *p*=0.1858). B) Percentage of VGluT1-positive mPFC^Dyn^ neurons and their levels of PDyn mRNA expression across the medial-lateral gradient of the mPFC. C) Same as above but for VGAT-positive neurons that co-express PDyn mRNA. D) Percentage of mPFC SST- and PV-positive interneurons that co-express PDyn mRNA (unpaired t-test, \*\**p*=0.0066). E) Representative image of PDyn (green) and Drd1a (red) mRNA expression in the mPFC of WT mice. The bar graph shows quantification of the co-expression of Drd1a mRNA in mPFC^Dyn^ cells. F) Tomato-positive cells from PDyn-iCre mice crossed with Ai14 tdTomato reporter mice do not reliably label cells that express PDyn mRNA during adulthood. Representative confocal image of 20X RNAscope in-situ hybridization showing that PDyn mRNA (green) is largely absent in the majority of tdTomato mRNA-positive cells (red), suggesting ectopic expression of tdTomato. G) Schematic and representative image of AAV-hSyn-FLEX-tdTomato-T2A-Synaptophysin-eGFP expression in the mPFC of PDyn-Cre mice. H) PDyn immunoreactivity in mPFC neurons expressing tdTomato-2A-Synaptophysin-eGFP in a Cre-dependent manner. I) Mean onset latency of mPFC^Dyn^ neurons evoked oEPSCs and oIPSCs consistent with direct monosynaptic excitatory and inhibitory connections, respectively (unpaired t-test, *p*=0.6032). The monosynaptic connections were also confirmed by TTX and 4-AP application (\**p*=0.0143, \*\**p*=0.0063, \*\*\**p*=0.0001). J) Schematic AAV-EF1α-DIO-eYFP expression in the mPFC of PDyn-Cre mice. K) Representative reconstructions of mPFC^Dyn^ cells exhibiting pyramidal and non-pyramidal morphologies indicate putative excitatory and inhibitory neurons, respectively. Pie chart shows percentage of pyramidal and putative inhibitory mPFC^Dyn^ cells. L) INTRSECT and hDlx promoter approaches to label mPFC^Dyn/SST^ neurons. M) Non-pyramidal mPFC^Dyn/SST^ cells labeled with C_ON_/F_ON_ INTRSECT and hDlx promoter approaches and excitatory pyramidal neurons labeled with C_ON_/F_OFF_ INTRSECT. I/V curve in current clamp recordings from these mPFC^Dyn/SST^ and mPFC^Dyn/Exc^ cells. N) Sholl analysis of mPFC^Dyn/SST^ neuron morphology. O) Summary of whole neuronal area, neurite length, basal dendrite area, number of basal dendritic trees, and number of dendritic ends of mPFC^Dyn/Exc^ and mPFC^Dyn/SST^ cells (\*\**p*<0.01, \*\*\*\**p*<0.0001). P) Additional representative images of SST immunoreactivity (red) on hDlx-Flex-GFP expression (green) mPFC cells as in Fig. 2L.

**Figure S3:**
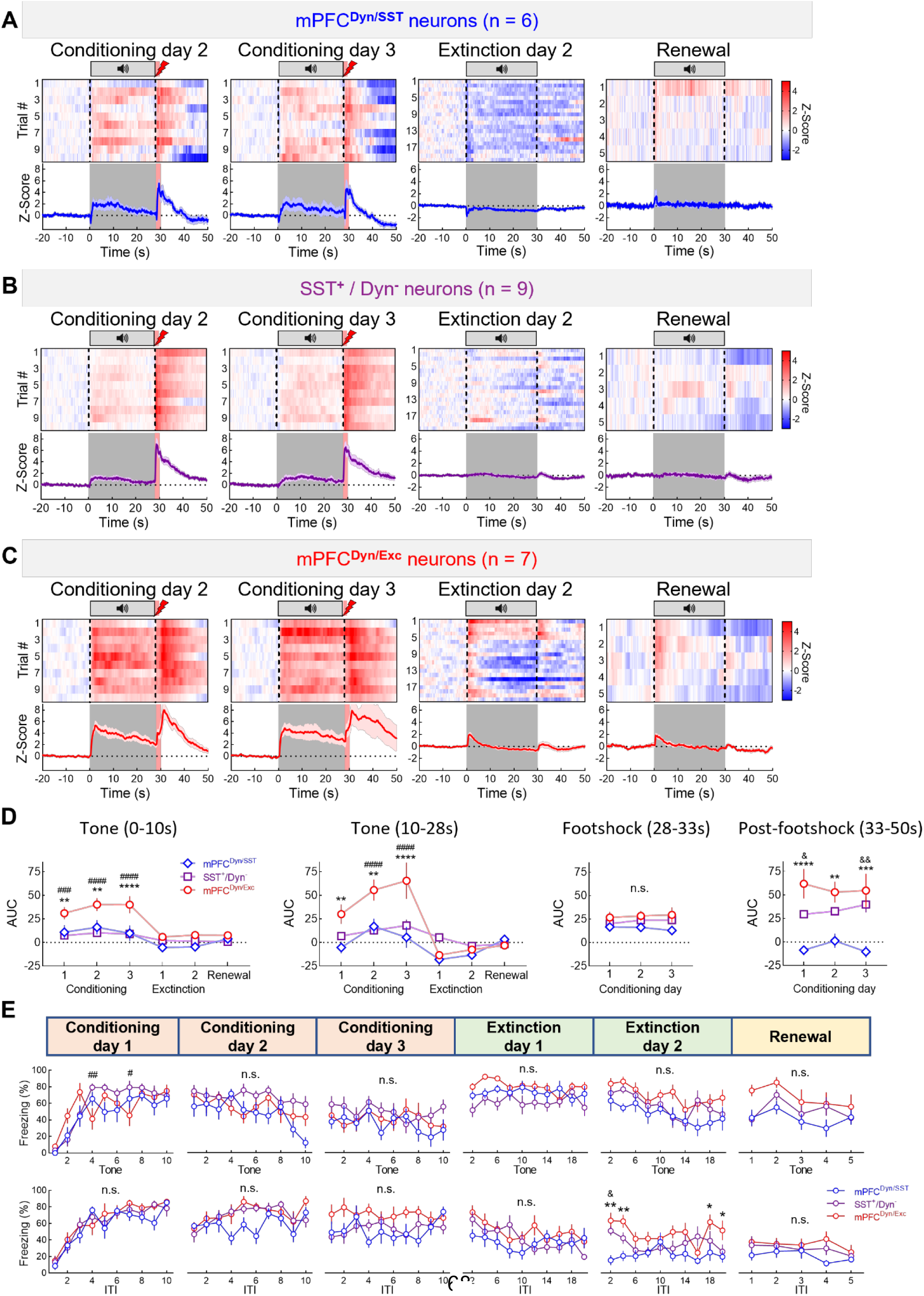
Relevant to Figure 3 Activity of excitatory and inhibitory vmPFC^Dyn^ cells during threat conditioning, extinction and renewal. A) Heatmaps and traces of GCaMP activity in mPFC^Dyn/SST^ neurons across the session during day 2 and day 3 of threat conditioning, day 2 of threat extinction, and renewal of threat responsitivity in the training context after extinction. B) Heatmaps and traces of GCaMP activity in SST^+^ /Dyn^-^ neurons across the session during day 2 and day 3 of threat conditioning, day 2 of threat extinction, and renewal of threat responsitivity in the training context after extinction. C) Heatmaps and traces of GCaMP activity in mPFC^Dyn/Exc^ neurons across the session during day 2 and day 3 of threat conditioning, day 2 of threat extinction, and renewal of threat responsitivity in the training context after extinction. D) AUC of Ca^2+^ activity during early tone (0-10s, Two-way ANOVA, cell-type main effect with Bonferroni’s post-hoc test, \*\**p*<0.01 \*\*\*\**p*<0.0001 between mPFC^Dyn/SST^ and mPFC^Dyn/Exc^ neurons, ^###^*p*<0.001 ^####^*p*<0.0001 between SST^+^/Dyn^-^ and mPFC^Dyn/Exc^ neurons), late tone (10-28s, Two-way ANOVA, cell-type main effect with Bonferroni’s post-hoc test, \*\**p*<0.01 \*\*\*\**p*<0.0001 between mPFC^Dyn/SST^ and mPFC^Dyn/Exc^ neurons, ^####^*p*<0.0001 between SST^+^/Dyn^-^ and mPFC^Dyn/Exc^ neurons), footshock (28-33s, Two-way ANOVA), and post footshock periods (33-50s, Two-way ANOVA, cell-type main effect with Bonferroni’s post-hoc test, \*\**p*=0.0049 \*\*\**p*=0.0003 \*\*\*\**p*<0.0001 between mPFC^Dyn/SST^ and mPFC^Dyn/Exc^ neurons, ^&^*p*=0.0386 ^&&^*p*=0.0041 between mPFC^Dyn/SST^ and SST^+^/Dyn^-^ neurons). E) Freezing during tone and ITI period across threat conditioning, threat recall/extinction day 1 and extinction day 2, and renewal in 3 INTRSECT groups of mice. Data is binned every 2 trials for extinction days (Two-way ANOVA, \**p*<0.05 \*\**p*<0.01 between mPFC^Dyn/SST^ and mPFC^Dyn/Exc^ group, ^#^*p*=0.0172 ^##^*p*=0.006 between SST^+^/Dyn^-^ and mPFC^Dyn/Exc^ group, ^&^*p*=0.0151 between mPFC^Dyn/SST^ and SST^+^/Dyn^-^ group).

**Figure S4:**
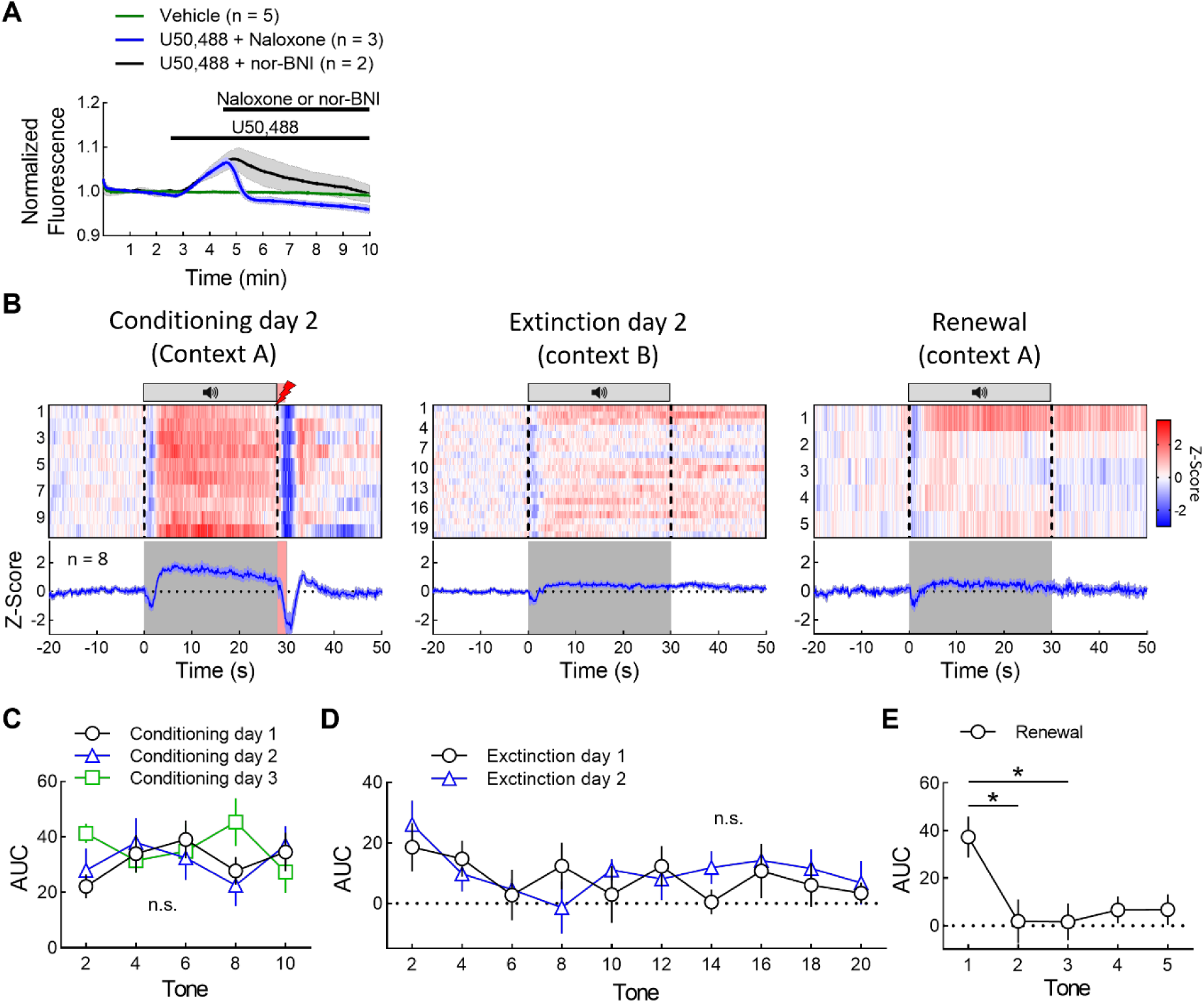
Relevant to Figure 4 In-vivo monitoring of putative/ dynorphin/KOR signaling. A) Naloxone or nor-BNI blocks increase of mPFC kLight1.2 fluorescence by U50,488 in slices. B) Heatmaps and traces of kLight activity in mPFC across the session during day 2 of threat conditioning, day 2 of threat extinction, and renewal of threat responsitivity. C-E) AUC of kLight signal during threat conditioning (C), threat extinction (D), and renewal of threat responsitivity (E). Data is binned every 2 trials in C and D (Two-way ANOVA for C and D, ANOVA with Tukey’s Post Hoc test for E, \**p*<0.05).

**Figure S5:**
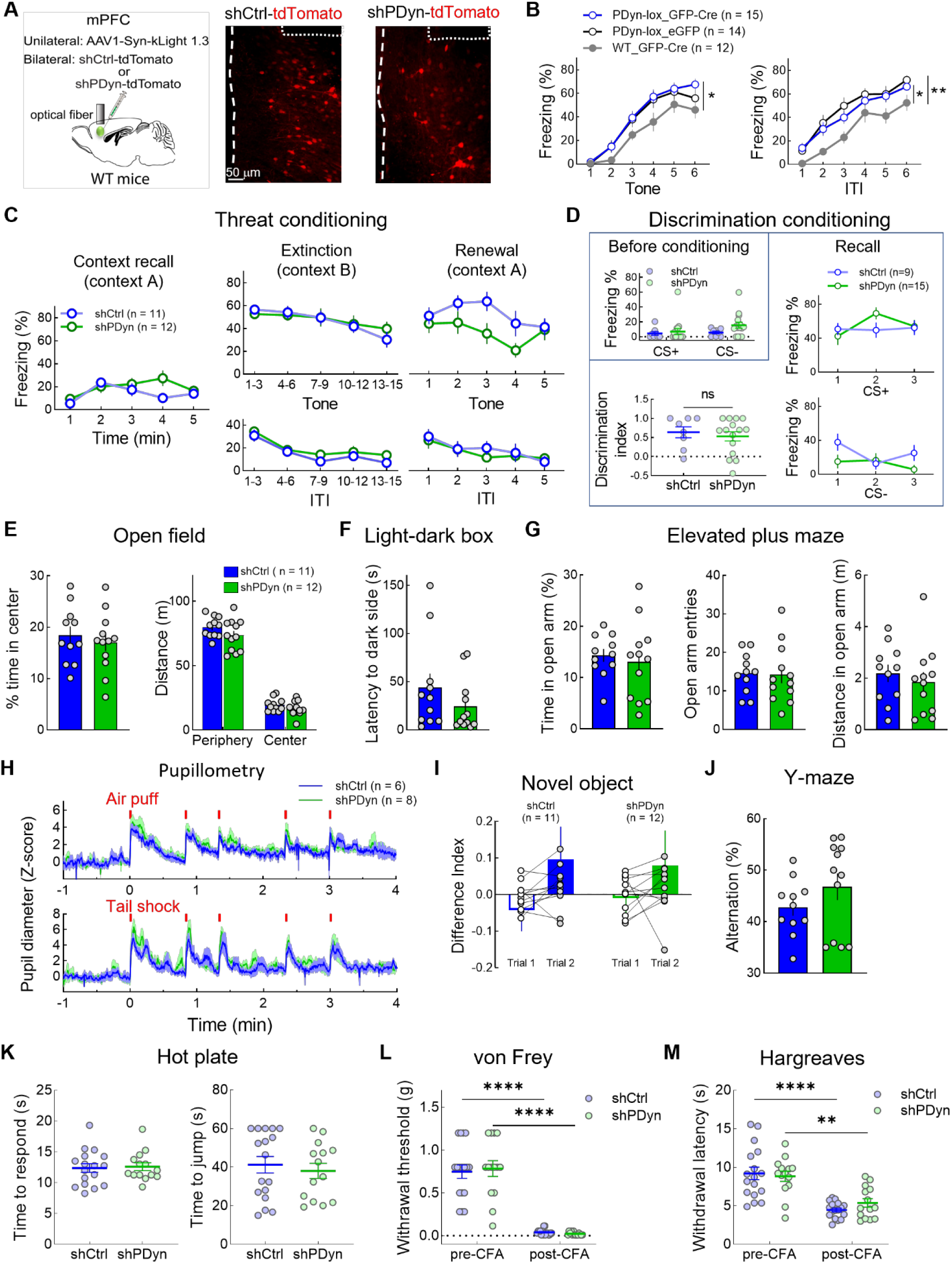
Relevant to Figure 5 PDyn knockdown in the vmPFC does not impact on recall or extinction of fear memory, recall of discriminative cues, locomotion, anxiety, autonomic response, and working memory. A) Representative images of shCtrl-tdTomato or shPDyn-tdTomato expression in WT mice of kLight fiber photometry recording. B) Freezing during tone and ITI in PDyn-loxP mice expressing AAV-Cre-GFP or AAV-eGFP, and in WT mice expressing AAV-Cre-GFP in the vmPFC (Two-way ANOVA with Bonferroni’s Post Hoc test, Tone: \**p*=0.0304 between PDyn-lox_GFP-Cre and WT_GFP-Cre; ITI: \**p*=0.0351 between PDyn-lox_GFP-Cre and WT_GFP-Cre, \*\**p*=0.0055 between PDyn-lox_eGFP and WT_GFP-Cre). C) Contextual threat recall, cued threat recall/extinction, and renewal of conditioned freezing. D) vmPFC Dyn knockdown does not modify basal freezing during the baseline of the cue discrimination task or during the cued recall of the CS+ and CS-. E) vmPFC PDyn does not regulate time spent in the center of the open field or total distance moved in the periphery or center of the arena. F) Latency to the dark side in the light dark-box test. G) No significant difference in elevated plus maze test. H) No significant differences in air-puff or shock-evoked pupil dilation. I) No significant difference in novel object recognition memory. J) No significant differences in working memory as assessed by the spontaneous alternation task in the Y-maze. K) No significant difference in time to respond and time to jump at the hot plate test. L) No significant difference in von Frey test before or after CFA treatment (Two-way ANOVA with Bonferroni’s Post Hoc test, Treatment Main Effect, \*\*\*\**p*<0.0001). M) No significant difference in Hargreaves test before or after CFA treatment (Two-way ANOVA with Bonferroni’s Post Hoc test, Treatment Main Effect, \*\**p*=0.0011, \*\*\*\**p*<0.0001). Same group of mice were used in Fig. 5F,H as in Fig. S5C, E-G, and I,J.

**Figure S6:**
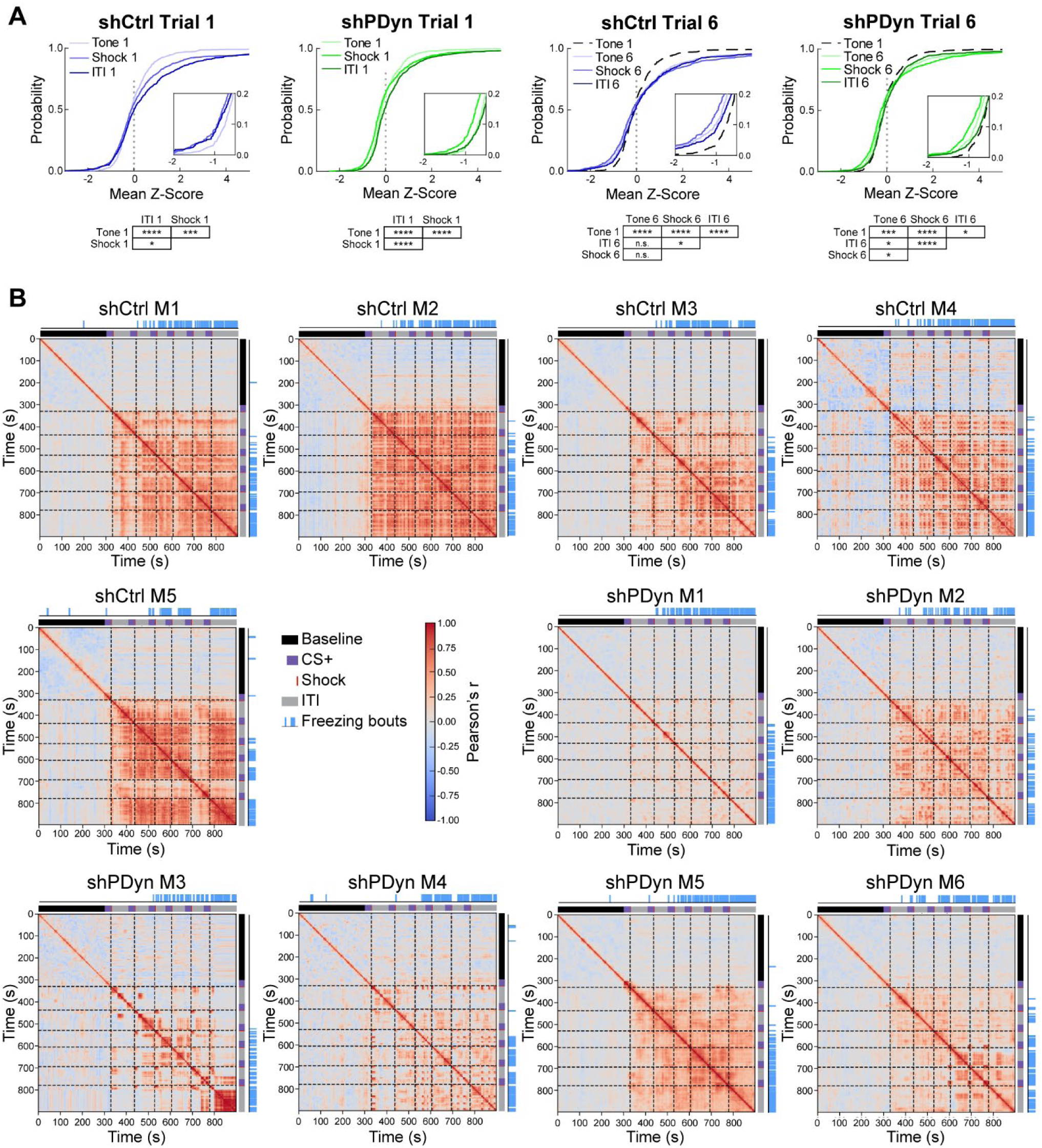
Relevant to Figure 6 and Figure 7. PCA trajectory and PCC data in individual mice. A) Cumulative probability of mean Z-Score in control (blue) and PDyn-shRNA (green) mice during the Tone periods, shock periods, and ITI from trial 1 and trial 6. Insets are zoomed in between Mean Z-score −2 to −0.5. Statistical significance from pair-wise Kolmogorov-Smirnov test is shown under each plot (\**p*<0.05, \*\*\**p*<0.001, \*\*\*\**p*<0.0001). B) Correlation matrices from individual mice in control-shRNA and PDyn-shRNA groups overlayed with freezing bouts.

