## Supplementary material for "Prefrontal cortical dynorphin peptidergic transmission constrains threat-driven behavioral and network states": Statistics Table

### Table 1

| Figure | Sub-figure | Statistical Test | One tailed or two tailed? | t, D, or F value | P value | If there is signifance, where the signifciance occurred (multiple comparison)? |
|---|---|---|---|---|---|---|
| **Fig.1B** | **Freezing in conditioning** | RM two-way ANOVA with Bonferroni's Post Hoc test | | Group Main Effect, F (1, 13) = 99.65 | p<0.0001 | Difference between tone only and tone + footshock group from Tone 3 to Tone10 |
| | | | | Trial Main Effect, F (9, 117) = 9.665 | p<0.0001 | |
| | | | | Group x Trial interaction, F (9, 117) = 6.607 | p<0.0001 | Difference between tone only and tone + footshock mice during specific epochs |
| **Fig.1C** | **Freezing in exctinction** | RM two-way ANOVA with Bonferroni's Post Hoc test | | Group Main Effect, F (1, 13) = 39.92 | p<0.0001 | Difference between tone only and tone + footshock group in all trials |
| | | | | Trial Main Effect, F (9, 117) = 0.9754 | p=0.4641 | |
| | | | | Group x Trial interaction, F (9, 117) = 0.7390 | p=0.6725 | |
| **Fig.1D** | **Conditioning day 1** | RM two-way ANOVA with Bonferroni's Post Hoc test | | Group Main Effect, F (1, 13) = 43.05 | P<0.0001 | Difference between tone only and tone + footshock mice |
| | | | | Time Main Effect, F (560, 7280) = 8.380 | P<0.0001 | |
| | | | | Group x Time interaction, F (560, 7280) = 10.86 | p<0.0001 | Difference between tone only and tone + footshock mice during specific epochs |
| **Fig.1D** | **Conditioning day 3** | RM two-way ANOVA with Bonferroni's Post Hoc test | | Group Main Effect, F(1,13)=11.12 | P=0.0054 | Difference between tone only and tone + footshock mice |
| | | | | Time Main Effect, F (560, 7280) = 5.692 | p<0.0001 | |
| | | | | Group x Time interaction, F (560, 7280) = 5.014 | p<0.0001 | Difference between tone only and tone + footshock mice during specific epochs |
| **Fig.1D** | **Extinction** | RM two-way ANOVA | | Group Main Effect, F(1,13)=0.3375 | p=0.5712 | |
| | | | | Time Main Effect, F(560,7280)=0.5750 | p>0.9999 | |
| | | | | Group x Time interaction, F(560,7280)=0.4342 | p>0.9999 | |
| **Fig.1E** | **AUC in conditioning day 1&3** | RM three-way ANOVA with Bonferroni's Post Hoc test | | Group Main Effect, F (1, 13) = 15.78 | p=0.0016 | Tone 1-3:tone+footshock day3 vs. Tone 1-3:tone only day1 (p=0.0017); Tone 1-3:tone+footshock day3 vs. Tone 1-3:tone only day3 (p=0.0389); |
| | | | | Days Main Effect, F (1, 13) = 4.180 | p=0.0617 | |
| | | | | Trial Main Effect, F (2, 26) = 0.09033 | p=0.9139 | |
| **Fig.1E** | **AUC in extinction day 1** | RM two-way ANOVA | | Group Main Effect, F (1, 13) = 0.3314 | p=0.5747 | |
| | | | | Trial Main Effect, F (9, 117) = 1.737 | P=0.0880 | |
| | | | | Group x Trial interaction, F (9, 117) = 1.453 | P=0.1736 | |
| **Fig.1I** | | RM wo-way ANOVA with Bonferroni's Post Hoc test | | Cell Main Effect, F (1, 10) = 1.965 | p=0.1912 | |
| | | | | Trial Main Effect, F (5, 50) = 2.854 | p=0.0241 | |
| | | | | Cell x Trial interaction, F (5, 50) = 3.811 | p=0.0053 | Footshock 1: WT vs. PDyn-Cre (p=0.0019) |
| **Fig.1J** | **Footshock 1** | RM two-way ANOVA with Bonferroni's Post Hoc test | | Cell Main Effect, F (1, 2184) = 184.2 | P<0.0001 | WT vs. PDyn-Cre: significant different from 1s to 7s. |
| | | | | Time Main Effect, F (23, 2184) = 14.93 | P<0.0001 | |
| | | | | Cell x Time interaction, F (23, 2184) = 7.227 | P<0.0001 | |
| **Fig.1J** | **Footshock 6** | RM two-way ANOVA with Bonferroni's Post Hoc test | | Cell Main Effect, F (1, 1848) = 89.13 | P<0.0001 | WT vs. PDyn-Cre: significant different from 3.5s to 7s. |
| | | | | Time Main Effect, F (23, 1848) = 30.45 | P<0.0001 | |
| | | | | Cell x Time interaction, F (23, 1848) = 6.980 | P<0.0001 | |
| **Fig.1K** | | RM two-way ANOVA with Bonferroni's Post Hoc test | | Cell Main Effect, F (1, 510) = 75.33 | p<0.0001 | Footshock 1: WT vs. PDyn-Cre (p<0.0001); Footshock 2: WT vs. PDyn-Cre (p=0.0002); Footshock 3: WT vs. PDyn-Cre (p=0.0098); Footshock 4: WT vs. PDyn-Cre (p=0.0309) |
| | | | | Trial Main Effect, F (5, 510) = 6.484 | p<0.0001 | |
| | | | | Cell x Trial interaction, F (5, 510) = 4.907 | p=0.0002 | |
| **Fig.1L** | | Paired t-test | two tailed | t=2.785 | p=0.0496 | |
| **Fig.1M** | | RM two-way ANOVA | | Cell Main Effect, F (1, 10) = 4.018 | p=0.0728 | |
| | | | | Trial Main Effect, F (5, 50) = 1.284 | p=0.2854 | |
| | | | | Cell x Trial interaction, F (5, 50) = 1.761 | p=0.1382 | |

| Figure | Sub-figure | Statistical Test | One tailed or two tailed? | t, D, or F value | P value | If there is signifance, where the signifciance occurred (multiple comparison)? |
|---|---|---|---|---|---|---|
| **Fig.1N** | **Tone 1** | RM two-way ANOVA with Bonferroni's Post Hoc test | | Cell Main Effect, F (1, 4320) = 240.9 | P<0.0001 | WT vs. PDyn-Cre: significant different from 32s to 35s. |
| | | | | Time Main Effect, F (79, 4320) = 1.706 | P=0.0001 | |
| | | | | Cell x Time interaction, F (79, 4320) = 1.487 | P=0.0036 | |
| **Fig.1N** | **Tone 6** | RM two-way ANOVA | | Cell Main Effect, F (1, 6400) = 7.352e-030 | P>0.9999 | |
| | | | | Time Main Effect, F (79, 6400) = 1.523 | P=0.0021 | |
| | | | | Cell x Time interaction, F (79, 6400) = 9.307e-032 | P>0.9999 | |
| **Fig.1O** | | RM two-way ANOVA with Bonferroni's Post Hoc test | | Cell Main Effect, F (1, 360) = 13.91 | p=0.0002 | **Tone 3:** WT vs. PDyn-Cre (p=0.0074) |
| | | | | Trial Main Effect, F (5, 360) = 2.856 | p=0.0152 | |
| | | | | Cell x Trial interaction, F (5, 360) = 1.678 | p=0.139 | |
| **Fig.1P** | | Paired t-test | two tailed | t=2.246 | p=0.0880 | |
| **Fig.1Q** | | Unpaired t-test | two tailed | t=1.260 | p=0.2364 | |
| **Fig.1S** | | Chi-square test | | Chi-square=4.342, df=8 | p=0.8251 | |
| **Fig. 1T** | | RM two-way ANOVA with Bonferroni's Post Hoc test | | Cell Main Effect, F (1, 155) = 2.328 | p=0.1291 | **Speed bin 48:** WT vs. PDyn-Cre (p=0.0382) |
| | | | | Speed Main Effect, F (16, 2480) = 4.088 | p<0.0001 | |
| | | | | Cell x Speed interaction, F (16, 2480) = 1.720 | p=0.0367 | |
| **Fig.2D** | | Two-way ANOVA with Bonferroni's Post Hoc test | | Sub-region Main Effect, F (1, 12) = 4.460e-013 | P>0.9999 | |
| | | | | Cell-Type Main Effect, F (1, 12) = 1011 | P<0.0001 | **PrL:**VGluT1 vs. PrL:VGAT (P<0.0001); **IL:**VGluT1 vs. IL:VGAT (P<0.0001); |
| | | | | Sub-region x Cell-Type interaction, F (1, 12) = 2.746 | P=0.1234 | |
| **Fig.2F** | | Unpaired t-test | two tailed | t=4.740 | p=0.0032 | |
| **Fig.2G** | | Kolmogorov-Smirnov test | | D=0.2677 | p<0.0001 | |
| **Fig.2I** | | Unpaired t-test | two tailed | t=3.513 | P=0.0007 | |
| **Fig.2N** | | RM two-way ANOVA | | AP frequency Main Effect, F (12, 234) = 8.509 | P<0.0001 | |
| | | | | Cell-Type Main Effect, F (1, 234) = 13.89 | P=0.0002 | |
| | | | | Sub-region x Cell-Type interaction, F (12, 234) = 0.2115 | P=0.9979 | |
| **Fig.3J** | **Tone (0-10s)** | RM two-way ANOVA with Bonferroni's Post Hoc test | | Days Main Effect, F (1, 18) = 28.80 | P<0.0001 | |
| | | | | Cell-type Main Effect, F (2, 18) = 8.611 | P=0.0024 | **Conditioning day1:** Con/Foff vs. Con/Fon (p = 0.0043); Con/Foff vs. Coff/Fon (p = 0.0001) |
| | | | | Days x Cell-type interaction, F (2, 18) = 4.623 | P=0.0240 | |
| **Fig.3J** | **Tone (10-28s)** | RM two-way ANOVA with Bonferroni's Post Hoc test | | Days Main Effect, F (1, 18) = 17.13 | P=0.0006 | |
| | | | | Cell-type Main Effect, F (2, 18) = 5.823 | P=0.0112 | **Conditioning day1:** Con/Foff vs. Con/Fon (p = 0.0008); Con/Foff vs. Coff/Fon (p = 0.0121). **Extinction day1:** Con/Fon vs. Coff/Fon (p = 0.0242) |
| | | | | Days x Cell-type interaction, F (2, 18) = 8.311 | P=0.0028 | |
| **Fig.3J** | **Footshock (28-33s)** | One-way ANOVA with Bonferroni's Post Hoc test | | F (2, 19) = 1.551 | P=0.2377 | |
| **Fig.3J** | **Post-footshock (33-50s)** | One-way ANOVA with Bonferroni's Post Hoc test | | F (2, 19) = 13.48 | P=0.0002 | Con/Foff vs. Con/Fon (p = 0.0002); Con/Fon vs. Coff/Fon (p = 0.0192) |
| **Fig.4D** | | RM two-way ANOVA | | Time Main Effect, F(2399,14394)=9.426 | p<0.0001 | |
| | | | | Treatment Main Effect, F(1,6)=10.98 | p=0.0161 | |
| | | | | Time x Treatment interaction, F(2399,14394)=11.57 | p<0.0001 | |
| **Fig.4K** | | RM two-way ANOVA with Bonferroni's Post Hoc test | | Treatment Main Effect, F (1, 16) = 7.690 | P=0.0136 | **kLight 1.2a:** Tone control vs. Conditioning (p = 0.0046) |
| | | | | Sensor Main Effect, F (1, 16) = 3.824 | P=0.0682 | **Conditioning:** kLight1.2 vs. kLight0 (P = 0.034) |
| | | | | Treatment x Sensor interaction, F (1, 16) = 3.277 | P=0.0891 | |

| Figure | Sub-figure | Statistical Test | One tailed or two tailed? | t, D, or F value | P value | If there is signifance, where the significiance occurred (multiple comparison)? |
|---|---|---|---|---|---|---|
| **Fig.4M** | **Number of Events** | RM two-way ANOVA with Bonferroni's Post Hoc test | | Trial Main Effect, F (1, 8) = 19.30 | P=0.0023 | **kLight 1.2a:** Baseline vs. Restrained (p = 0.001) |
| | | | | Sensor Main Effect, F (1, 8) = 7.292 | P=0.0271 | **Restrained:** kLight1.2 vs. kLight0 (P = 0.003) |
| | | | | Trial x Sensor interaction, F (1, 8) = 7.292 | P=0.0271 | |
| **Fig.4M** | **AUC** | RM two-way ANOVA with Bonferroni's Post Hoc test | | Trial Main Effect, F (1, 8) = 3.083 | P=0.1172 | **kLight 1.2a:** Baseline vs. Restrained (p = 0.0087) |
| | | | | Sensor Main Effect, F (1, 8) = 11.37 | P=0.0097 | **Restrained:** kLight1.2 vs. kLight0 (P = 0.0005) |
| | | | | Trial x Sensor interaction, F (1, 8) = 10.32 | P=0.0124 | |
| **Fig.5B** | **Z-Score** | RM two-way ANOVA with Bonferroni's Post Hoc test | | shRNA Main Effect, F(1, 9) = 0.03346 | P=0.8589 | |
| | | | | Time Main Effect, F (560.0, 5040) = 4.996 | p<0.0001 | |
| | | | | shRNA x Time interaction,F (560, 5040) = 2.245 | p<0.0001 | |
| **Fig.5C** | **Z-Score** | RM two-way ANOVA with Bonferroni's Post Hoc test | | shRNA Main Effect, F (1, 8) = 0.2007 | P=0.6660 | |
| | | | | Time Main Effect, F (560.0, 4480) = 4.429 | p<0.0001 | |
| | | | | shRNA x Time interaction,F (560, 4480) = 2.669 | p<0.0001 | |
| **Fig.5D** | **Conditioning** | RM two-way ANOVA | | shRNA Main Effect, F (1, 9) = 2.360 | P=0.1589 | |
| | | | | Trial Main Effect, F (4, 36) = 0.7895 | P=0.5397 | |
| | | | | shRNA x Trial interaction, F (4, 36) = 0.5907 | P=0.6715 | |
| **Fig.5D** | **Extinction** | RM two-way ANOVA | | shRNA Main Effect, F (1, 9) =0.05216 | P=0.8245 | |
| | | | | Trial Main Effect, F (9, 81) = 1.898 | P=0.0638 | |
| | | | | shRNA x Trial interaction, F (9, 81) = 3.504 | P=0.0011 | **Tone 2:** shCtrl vs shPDyn (p=0.0012) |
| **Fig.5F** | **Mean Intensity** | Unpaired t-test | two tailed | t=4.582 | P=0.0010 | |
| **Fig.5F** | **# of cells / mm$^2$** | Unpaired t-test | two tailed | t=4.459 | p=0.0012 | |
| **Fig.5H** | **Tone** | RM two-way ANOVA with Bonferroni's Post Hoc test | | Trial Main Effect, F (5, 105) = 51.75 | P<0.0001 | |
| | | | | shRNA Main Effect, F (1, 21) = 2.828 | P=0.1075 | |
| | | | | Trial x shRNA interaction, F (5, 105) = 2.759 | P=0.0221 | Tone3:shCtrl vs. shPDyn (P=0.0357) |
| **Fig.5H** | **ITI** | RM two-way ANOVA with Bonferroni's Post Hoc test | | Trial Main Effect, F (5, 105) = 63.82 | P<0.0001 | |
| | | | | shRNA Main Effect, F (1, 21) = 12.82 | P=0.0018 | Tone2:shCtrl vs. shPDyn (P=0.0157); Tone3:shCtrl vs. shPDyn (P=0.0082); Tone5:shCtrl vs. shPDyn (P=0.0013) |
| | | | | Trial x shRNA interaction, F (5, 105) = 2.125 | P=0.0682 | |
| **Fig.5J** | **CS+** | RM two-way ANOVA | | Trial Main Effect, F (9, 198) = 28.20 | P<0.0001 | |
| | | | | shRNA Main Effect, F (1, 22) = 2.249 | P=0.1479 | |
| | | | | Trial x shRNA interaction, F (9, 198) = 1.157 | P=0.3245 | |
| **Fig.5J** | **CS-** | RM two-way ANOVA with Bonferroni's Post Hoc test | | Trial Main Effect, F (9, 198) = 20.42 | P<0.0001 | |
| | | | | shRNA Main Effect, F (1, 22) = 6.453 | P=0.0187 | **Trial7**:shCtrl vs. shPDyn (P=0.0229); **Trial9:**shCtrl vs. shPDyn (P=0.0483); **Trial10:**shCtrl vs. shPDyn (P=0.0452) |
| | | | | Trial x shRNA interaction, F (9, 198) = 2.928 | P=0.0028 | |
| **Fig.5K** | | Unpaired t-test | two tailed | t=2.473 | p=0.0216 | |
| **Fig.6C** | | Kolmogorov-Smirnov test | | D=0.001389 | P>0.9999 | |
| **Fig.6E** | **Tone** | Kolmogorov-Smirnov test | | D=0.04697 | P=0.3108 | |
| **Fig.6E** | **Footshock** | Kolmogorov-Smirnov test | | D=0.09809 | P=0.0006 | |
| **Fig.6E** | **ITI** | Kolmogorov-Smirnov test | | D=0.1187 | P<0.0001 | |
| **Fig.6G** | **Tone** | Kolmogorov-Smirnov test | | D=0.07091 | P=0.0290 | |
| **Fig.6G** | **Footshock** | Kolmogorov-Smirnov test | | D=0.08962 | P=0.0023 | |
| **Fig.6G** | **ITI** | Kolmogorov-Smirnov test | | D=0.1141 | P<0.0001 | |

| Figure | Sub-figure | Statistical Test | One tailed or two tailed? | t, D, or F value | P value | If there is signifance, where the signifciance occurred (multiple comparison)? |
|---|---|---|---|---|---|---|
| **Fig.6H** | | Two-way ANOVA | | Trial Main Effect, F (5, 45) = 4.294 | P=0.0028 | |
| | | | | shRNA Main Effect, F (1, 9) = 0.5528 | P=0.4761 | |
| | | | | Trial x shRNA interaction, F (5, 45) = 1.025 | P=0.4143 | |
| **Fig.6J** | | Two-way ANOVA | | Trial Main Effect, F (5, 894) = 0.2352 | P=0.9471 | |
| | | | | shRNA Main Effect, F (1, 894) = 3.210 | P=0.0735 | |
| | | | | Trial x shRNA interaction, F (5, 894) = 0.1597 | P=0.9771 | |
| **Fig.6K** | | Two-way ANOVA | | Trial Main Effect, F (5, 45) = 3.767 | P=0.0062 | |
| | | | | shRNA Main Effect, F (1, 9) = 0.2084 | P=0.6588 | |
| | | | | Trial x shRNA interaction, F (5, 45) = 0.6519 | P=0.6615 | |
| **Fig.6M** | | Two-way ANOVA | | Trial Main Effect, F (5, 808) = 0.5240 | P=0.7583 | |
| | | | | shRNA Main Effect, F (1, 808) = 0.4076 | P=0.5234 | |
| | | | | Trial x shRNA interaction, F (5, 808) = 0.8868 | P=0.4893 | |
| **Fig.6N** | | Two-way ANOVA | | Trial Main Effect, F (5, 45) = 3.525 | P=0.0090 | |
| | | | | shRNA Main Effect, F (1, 9) = 3.450 | P=0.0962 | |
| | | | | Trial x shRNA interaction, F (5, 45) = 0.7599 | P=0.5835 | |
| **Fig.6P** | | Two-way ANOVA | | Trial Main Effect, F (5, 745) = 0.2647 | P=0.9323 | |
| | | | | shRNA Main Effect, F (1, 745) = 2.122 | P=0.1456 | |
| | | | | Trial x shRNA interaction, F (5, 745) = 2.380 | P=0.0372 | |
| **Fig.6Q** | | Unpaired t-test | two tailed | t=2.255 | p=0.0506 | |
| **Fig.6S** | | Chi-square test | | Chi-square=10.99, df=7 | p=0.1392 | |
| **Fig.6T** | | Two-way ANOVA with Bonferroni's Post Hoc test | | Speed Main Effect, F (16, 5696) = 10.28 | P<0.0001 | |
| | | | | shRNA Main Effect, F (1, 356) = 1.337 | P=0.2484 | |
| | | | | Speed x shRNA interaction, F (16, 5696) = 2.061 | P=0.0076 | |
| **Fig.7B** | | RM two-way ANOVA with Bonferroni's Post Hoc test | | Trial Main Effect, F (5, 30) = 5.566 | P=0.0010 | |
| | | | | shRNA Main Effect, F (1, 6) = 0.3320 | P=0.5855 | |
| | | | | Trial x shRNA interaction, F (5, 30) = 1.047 | P=0.4087 | |
| **Fig.7C** | | RM two-way ANOVA with Bonferroni's Post Hoc test | | Trial Main Effect, F (5, 30) = 6.156 | P=0.0005 | |
| | | | | shRNA Main Effect, F (1, 6) = 6.828 | P=0.0400 | |
| | | | | Trial x shRNA interaction, F (5, 30) = 1.040 | P=0.4124 | |
| **Fig.7D** | | RM two-way ANOVA with Bonferroni's Post Hoc test | | Trial Main Effect, F (5, 30) = 2.758 | P=0.0364 | |
| | | | | shRNA Main Effect, F (1, 6) = 10.82 | P=0.0166 | shCtrl_trial3 vs. shPDyn_trial3 (P=0.0250); shCtrl_trial5 vs. shPDyn_trial5 (P=0.0176) |
| | | | | Trial x shRNA interaction, F (5, 30) = 0.5040 | P=0.7708 | |
| **Fig.7E** | | RM two-way ANOVA with Bonferroni's Post Hoc test | | Trial Main Effect, F (5, 30) = 40.05 | P<0.0001 | |
| | | | | shRNA Main Effect, F (1, 6) = 0.1791 | P=0.6869 | |
| | | | | Trial x shRNA interaction, F (5, 30) = 0.4782 | P=0.7896 | |
| **Fig.7F** | | RM two-way ANOVA with Bonferroni's Post Hoc test | | Trial Main Effect, F (5, 30) = 12.02 | P<0.0001 | |
| | | | | shRNA Main Effect, F (1, 6) = 0.1079 | P=0.7537 | |
| | | | | Trial x shRNA interaction, F (5, 30) = 0.4053 | P=0.8413 | |

| Figure | Sub-figure | Statistical Test | One tailed or two tailed? | t, D, or F value | P value | If there is signifance, where the signifciance occurred (multiple comparison)? |
|---|---|---|---|---|---|---|
| **Fig.7G** | | RM two-way ANOVA with Bonferroni's Post Hoc test | | Trial Main Effect, F (5, 30) = 27.19 | P<0.0001 | |
| | | | | shRNA Main Effect, F (1, 6) = 9.971 | P=0.0196 | shCtrl_trial2 vs. shPDyn_trial2 (P=0.0255); shCtrl_trial3 vs. shPDyn_trial3 (P=0.0046); shCtrl_trial6 vs. shPDyn_trial6 (P=0.0367) |
| | | | | Trial x shRNA interaction, F (5, 30) = 1.070 | P=0.3965 | |
| **Fig.7I** | | Unpaired t-test | two tailed | t=1.817 | P=0.1027 | |
| **Fig.7J** | | Two-way ANOVA with Bonferroni's Post Hoc test | | Data Main Effect, F (1, 18) = 53.86 | P<0.0001 | **shCtrl:** Actural data vs. Shuffled data (P<0.0001);**shPDyn:** Actural data vs. Suffled data (P=0.0013) |
| | | | | shRNA Main Effect, F (1, 18) = 3.197 | P=0.0906 | **Actural data:** shCtrl vs. shPDyn (p=0.0419) |
| | | | | Data x shRNA interaction, F (1, 18) = 3.206 | P=0.0902 | |
| **Supp Fig.1A** | **Conditioning day1, Tone** | RM two-way ANOVA with Bonferroni's Post Hoc test | | Group Main Effect, F (1, 13) = 99.65 | p<0.0001 | Difference between tone only and tone + footshock group from Tone 3 to Tone10 |
| | | | | Trial Main Effect, F (9, 117) = 9.665 | p<0.0001 | |
| | | | | Group x Trial interaction, F (9, 117) = 6.607 | p<0.0001 | |
| **Supp Fig.1A** | **Conditioning day2, Tone** | RM two-way ANOVA with Bonferroni's Post Hoc test | | Group Main Effect, F (1, 13) = 53.64 | p<0.0001 | Difference between tone only and tone + footshock group in all trials |
| | | | | Trial Main Effect, F (9, 117) = 0.4923 | p=0.8773 | |
| | | | | Group x Trial interaction, F (9, 117) = 0.5008 | p=0.8714 | |
| **Supp Fig.1A** | **Conditioning day3, Tone** | RM two-way ANOVA with Bonferroni's Post Hoc test | | Group Main Effect, F (1, 13) = 25.87 | p=0.0002 | Difference between tone only and tone + footshock group in all trials |
| | | | | Trial Main Effect, F (9, 117) = 1.321 | p=0.2334 | |
| | | | | Group x Trial interaction, F (9, 117) = 1.515 | p=0.1505 | |
| **Supp Fig.1A** | **Extinction day1, Tone** | RM two-way ANOVA with Bonferroni's Post Hoc test | | Group Main Effect, F (1, 13) = 39.92 | p<0.0001 | Difference between tone only and tone + footshock group in all trials |
| | | | | Trial Main Effect, F (9, 117) = 0.9754 | p=0.4641 | |
| | | | | Group x Trial interaction, F (9, 117) = 0.7390 | p=0.6725 | |
| **Supp Fig.1A** | **Extinction day2, Tone** | RM two-way ANOVA with Bonferroni's Post Hoc test | | Group Main Effect, F (1, 13) = 26.25 | p=0.0002 | Difference between tone only and tone + footshock group in all trials |
| | | | | Trial Main Effect, F (9, 117) = 1.246 | p=0.2742 | |
| | | | | Group x Trial interaction, F (9, 117) = 1.286 | p=0.2517 | |
| **Supp Fig.1A** | **Renewal, Tone** | RM two-way ANOVA with Bonferroni's Post Hoc test | | Group Main Effect, F (1, 13) = 5.048 | p=0.0427 | Tone1: Tone only vs. Tone+footshock (p=0.0137) |
| | | | | Trial Main Effect, F (4, 52) = 1.707 | p=0.1625 | |
| | | | | Group x Trial interaction, F (4, 52) = 1.867 | p=0.1303 | |
| **Supp Fig.1A** | **Conditioning day1, ITI** | RM two-way ANOVA with Bonferroni's Post Hoc test | | Group Main Effect, F (1, 13) = 64.64 | p<0.0001 | Difference between tone only and tone + footshock group from ITI 3 to ITI 10 |
| | | | | Trial Main Effect, F (9, 117) = 6.319 | p<0.0001 | |
| | | | | Group x Trial interaction, F (9, 117) = 3.856 | p=0.0003 | |
| **Supp Fig.1A** | **Conditioning day2, ITI** | RM two-way ANOVA with Bonferroni's Post Hoc test | | Group Main Effect, F (1, 13) = 105.1 | p<0.0001 | Difference between tone only and tone + footshock group in all trials |
| | | | | Trial Main Effect, F (9, 117) = 2.115 | p=0.0335 | |
| | | | | Group x Trial interaction, F (9, 117) = 1.482 | p=0.1626 | |
| **Supp Fig.1A** | **Conditioning day3, ITI** | RM two-way ANOVA with Bonferroni's Post Hoc test | | Group Main Effect, F (1, 13) = 81.16 | p<0.0001 | Difference between tone only and tone + footshock group in all trials |
| | | | | Trial Main Effect, F (9, 117) = 1.048 | p=0.4069 | |
| | | | | Group x Trial interaction, F (9, 117) = 0.7321 | p=0.6788 | |

| Figure | Sub-figure | Statistical Test | One tailed or two tailed? | t, D, or F value | P value | If there is signifance, where the signifciance occurred (multiple comparison)? |
|---|---|---|---|---|---|---|
| **Supp Fig.1A** | **Extinction day1, ITI** | RM two-way ANOVA with Bonferroni's Post Hoc test | | Group Main Effect, F (1, 13) = 15.18 | p=0.0018 | Difference between tone only and tone + footshock group in ITI2, ITI4, ITI10, ITI12, ITI14, and ITI16 |
| | | | | Trial Main Effect, F (9, 117) = 1.098 | p=0.3699 | |
| | | | | Group x Trial interaction, F (9, 117) = 1.190 | p=0.3077 | |
| **Supp Fig.1A** | **Extinction day2, ITI** | RM two-way ANOVA with Bonferroni's Post Hoc test | | Group Main Effect, F (1, 13) = 16.80 | p=0.0013 | Difference between tone only and tone + footshock group in ITI2, ITI4, ITI10, and ITI12 |
| | | | | Trial Main Effect, F (9, 117) = 0.7721 | p=0.6424 | |
| | | | | Group x Trial interaction, F (9, 117) = 0.5106 | p=0.8645 | |
| **Supp Fig.1A** | **Renewal, ITI** | RM two-way ANOVA with Bonferroni's Post Hoc test | | Group Main Effect, F (1, 13) = 2.535 | p=0.1354 | |
| | | | | Trial Main Effect, F (4, 52) = 0.1675 | p=0.9539 | |
| | | | | Group x Trial interaction, F (4, 52) = 0.4318 | p=0.7850 | |
| **Supp Fig.1B** | **Conditioning day2** | RM two-way ANOVA with Bonferroni's Post Hoc test | | Group Main Effect, F (1, 13) = 51.86 | P<0.0001 | Difference between tone only and tone + footshock mice in multiple epochs |
| | | | | Time Main Effect, F (560, 7280) = 8.394 | P<0.0001 | |
| | | | | Group x Time interaction, F (560, 7280) = 9.578 | p<0.0001 | |
| **Supp Fig.1B** | **Extinction day2** | RM two-way ANOVA | | Group Main Effect, F (1, 13) = 0.4369 | P=0.5202 | |
| | | | | Time Main Effect, F (560, 7280) = 0.5223 | P>0.9999 | |
| | | | | Group x Time interaction, F (560, 7280) = 0.5495 | P>0.9999 | |
| **Supp Fig.1B** | **Renewal** | RM two-way ANOVA | | Group Main Effect, F (1, 13) = 1.473 | P=0.2464 | |
| | | | | Time Main Effect, F (560, 7280) = 0.6172 | P>0.9999 | |
| | | | | Group x Time interaction, F (560, 7280) = 0.9297 | p=0.8741 | |
| **Supp Fig.1C** | | RM two-way ANOVA | | Group Main Effect, F (1, 13) = 19.98 | P=0.0006 | Day1: Tone only vs. Tone+footshock (p=0.0075); Day2: Tone only vs. Tone+footshock (p=0.0005); Day3: Tone only vs. Tone+footshock (p=0.0009) |
| | | | | Days Main Effect, F (2, 26) = 3.190 | P=0.0577 | |
| | | | | Group x Days interaction, F (2, 26) = 0.5919 | P=0.5605 | |
| **Supp Fig.1D** | | RM two-way ANOVA | | Group Main Effect, F (1, 13) = 0.5352 | P=0.4774 | |
| | | | | Days Main Effect, F (1, 13) = 0.1050 | P=0.7511 | |
| | | | | Group x Days interaction, F (1, 13) = 0.09517 | P=0.7626 | |
| **Supp Fig.1E** | | Unpaired t-test | two tailed | t=1.257 | P=0.2309 | |
| **Supp Fig.1F** | | RM two-way ANOVA | | Cell Main Effect, F (1, 10) = 0.2615 | p=0.6202 | |
| | | | | Trial Main Effect, F (5, 50) = 0.5202 | p=0.7598 | |
| | | | | Cell x Trial interaction, F (5, 50) = 1.573 | p=0.1848 | |
| **Supp Fig.1G** | **ITI 1** | RM two-way ANOVA | | Cell Main Effect, F (1, 6890) = 0.2456 | P=0.6202 | |
| | | | | Time Main Effect, F (129, 6890) = 0.9264 | P=0.7124 | |
| | | | | Cell x Time interaction, F (129, 6890) = 0.2632 | P>0.9999 | |
| **Supp Fig.1G** | **ITI 6** | RM two-way ANOVA with Bonferroni's Post Hoc test | | Cell Main Effect, F (1, 6853) = 41.72 | P<0.0001 | |
| | | | | Time Main Effect, F (129, 6853) = 2.899 | P<0.0001 | |
| | | | | Cell x Time interaction, F (129, 6853) = 0.3288 | P>0.9999 | |
| **Supp Fig.1H** | | RM two-way ANOVA | | Cell Main Effect, F (1, 328) = 3.227 | p=0.0734 | |
| | | | | Trial Main Effect, F (5, 328) = 0.9110 | p=0.474 | |
| | | | | Cell x Trial interaction, F (5, 328) = 0.7953 | p=0.5536 | |

| Figure | Sub-figure | Statistical Test | One tailed or two tailed? | t, D, or F value | P value | If there is signifance, where the significiance occurred (multiple comparison)? |
|---|---|---|---|---|---|---|
| **Supp Fig.1I** | | Paired t-test | two tailed | t=0.4533 | p=0.6738 | |
| **Supp Fig.2A** | | Unpaired t-test | two tailed | t=1.494 | P=0.1858 | |
| **Supp Fig.2D** | | Unpaired t-test | two tailed | t=4.062 | P=0.0066 | |
| **Supp Fig.2I** | **oPSC Latency** | Unpaired t-test | two tailed | t=0.5213 | P=0.6032 | |
| **Supp Fig.2I** | **oEPSC Amplitude** | RM one-way ANOVA with Tukey's Post Hoc test | | Treatment, F (2, 12) = 19.12 | P=0.0002 | aCSF vs. TTX (P=0.0001); aCSF vs. TTX+4AP (P=0.0063) |
| **Supp Fig.2I** | **oIPSC Amplitude** | RM one-way ANOVA with Tukey's Post Hoc test | | Treatment, F (2, 8) = 7.054 | P=0.0171 | aCSF vs. TTX (P=0.0143) |
| **Supp Fig.2O** | **Whole neuronal area** | Unpaired t-test | two tailed | t=5.520 | P<0.0001 | |
| **Supp Fig.2O** | **Neurite length** | Unpaired t-test | two tailed | t=3.362 | P=0.0015 | |
| **Supp Fig.2O** | **Basal dendrite area** | Unpaired t-test | two tailed | t=0.7566 | P=0.4530 | |
| **Supp Fig.2O** | **# Basal dendritic trees** | Unpaired t-test | two tailed | t=3.376 | P=0.0014 | |
| **Supp Fig.2O** | **# Basal dendritic ends** | Unpaired t-test | two tailed | t=1.680 | P=0.0992 | |
| **Supp Fig.3D** | **Tone (0-10s)** | RM two-way ANOVA with Bonferroni's Post Hoc test | | Cell Type Main Effect, F (2, 18) = 11.48 | P=0.0006 | **Conditioning day1:** Con/Foff vs. Con/Fon (p=0.0097); **Conditioning day1:** Con/Foff vs. Coff/Fon (p=0.0002); **Conditioning day2:** Con/Foff vs. Con/Fon (p=0.0015); **Conditioning day2:** Con/Foff vs. Coff/Fon (p<0.0001); **Conditioning day3:** Con/Foff vs. Con/Fon (p<0.0001); **Conditioning day3**: Con/Foff vs. Coff/Fon (p<0.0001). |
| | | | | Days Main Effect, F (5, 90) = 21.50 | P<0.0001 | **Con/Foff:** Conditioning day1 vs. Extinction day1 (p<0.0001); **Con/Foff:** Conditioning day1 vs. Extinction Day2 (p=0.0002); **Con/Foff:** Conditioning day1 vs. Renewal (p=0.0001); **Con/Foff:** Conditioning day2 vs. Extinction day1 (p<0.0001); **Con/Foff:** Conditioning day2 vs. Extinction day2 (p<0.0001); **Con/Foff:** Conditioning day2 vs. Renewal (p<0.0001); **Con/Foff:** Conditioning day3 vs. Extinction day1 (p<0.0001); **Con/Foff:** Conditioning day3 vs. Extinction day2 (p<0.0001); **Con/Foff:** Conditioning day3 vs. Renewal (p<0.0001); **Con/Fon:** Conditioning day2 vs. Extinction day1 (p=0.0067); **Con/Fon:** Conditioning day2 vs. Extinction day2 (p=0.0098). |
| | | | | Cell Type x Days interaction, F (10, 90) = 4.024 | P=0.0001 | |
| **Supp Fig.3D** | **Tone (10-28s)** | RM two-way ANOVA with Bonferroni's Post Hoc test | | Cell Type Main Effect, F (2, 18) = 6.638 | P=0.0069 | **Conditioning day1:** Con/Foff vs. Con/Fon (p=0.0069); **Conditioning day2:** Con/Foff vs. Con/Fon (p=0.0027); **Conditioning day2:** Con/Foff vs. Coff/Fon (p<0.0001); **Conditioning day3:** Con/Foff vs. Con/Fon (p<0.0001); **Conditioning day3**: Con/Foff vs. Coff/Fon (p<0.0001). |
| | | | | Days Main Effect, F (5, 90) = 20.07 | P<0.0001 | **Con/Foff:** Conditioning day1 vs. Conditioning day3 (p=0.0034); **Con/Foff:** Conditioning day1 vs. Extinction day1 (p=0.0001); **Con/Foff:** Conditioning day1 vs. Extinction Day2 (p=0.0015); **Con/Foff:** Conditioning day1 vs. Renewal (p=0.0077); **Con/Foff:** Conditioning day2 vs. Extinction day1 (p<0.0001); **Con/Foff:** Conditioning day2 vs. Extinction day2 (p<0.0001); **Con/Foff:** Conditioning day2 vs. Renewal (p<0.0001); **Con/Foff:** Conditioning day3 vs. Extinction day1 (p<0.0001); **Con/Foff:** Conditioning day3 vs. Extinction day2 (p<0.0001); **Con/Foff:** Conditioning day3 vs. Renewal (p<0.0001); **Con/Fon:** Conditioning day2 vs. Extinction day1 (p=0.0272). |
| | | | | Cell Type x Days interaction, F (10, 90) = 5.955 | P<0.0001 | |
| **Supp Fig.3D** | **Footshock (28-33s)** | RM two-way ANOVA | | Cell Type Main Effect, F (2, 18) = 1.984 | P=0.1665 | |
| | | | | Days Main Effect, F (2, 36) = 0.4397 | P=0.6476 | |
| | | | | Cell Type x Days interaction, F (4, 36) = 0.7836 | P=0.5433 | |
| **Supp Fig.3D** | **Post-footshock (33-50s)** | RM two-way ANOVA with Bonferroni's Post Hoc test | | Cell Type Main Effect, F (2, 18) = 10.24 | P=0.0011 | **Conditioning day1:** Con/Foff vs. Con/Fon (p<0.0001); **Conditioning day1:** Con/Fon vs. Coff/Fon (p=0.0386); **Conditioning day2:** Con/Foff vs. Con/Fon (p=0.0049); **Conditioning day3:** Con/Foff vs. Con/Fon (p=0.0003); **Conditioning day3**: Con/Fon vs. Coff/Fon (p=0.0041). |
| | | | | Days Main Effect, F (2, 36) = 0.04810 | P=0.9531 | |
| | | | | Cell Type x Days interaction, F (4, 36) = 1.311 | P=0.2844 | |
| **Supp Fig.3E** | **Conditioning day1, Tone** | RM two-way ANOVA with Bonferroni's Post Hoc test | | Group Main Effect, F (2, 19) = 1.081 | P=0.3591 | |
| | | | | Trial Main Effect, F (9, 171) = 23.48 | P<0.0001 | |
| | | | | Group x Trial interaction, F (18, 171) = 2.505 | P=0.0012 | **Tone4:** Coff/Fon vs. Con/Foff (p=0.006); **Tone7:** Coff/Fon vs. Con/Foff (p=0.0172) |

| Figure | Sub-figure | Statistical Test | One tailed or two tailed? | t, D, or F value | P value | If there is signifance, where the signifciance occurred (multiple comparison)? |
|---|---|---|---|---|---|---|
| **Supp Fig.3E** | **Conditioning day2, Tone** | RM two-way ANOVA | | Group Main Effect, F (2, 19) = 3.198 | P=0.0635 | |
| | | | | Trial Main Effect, F (9, 171) = 3.677 | P=0.0003 | |
| | | | | Group x Trial interaction, F (18, 171) = 1.273 | P=0.2112 | |
| **Supp Fig.3E** | **Conditioning day3, Tone** | RM two-way ANOVA | | Group Main Effect, F (2, 19) = 1.178 | P=0.3295 | |
| | | | | Trial Main Effect, F (9, 171) = 1.378 | P=0.2016 | |
| | | | | Group x Trial interaction, F (18, 171) = 0.8264 | P=0.6675 | |
| **Supp Fig.3E** | **Extinction day1, Tone** | RM two-way ANOVA | | Group Main Effect, F (2, 19) = 2.904 | P=0.0794 | |
| | | | | Trial Main Effect, F (9, 171) = 1.038 | P=0.4121 | |
| | | | | Group x Trial interaction, F (18, 171) = 1.406 | P=0.1338 | |
| **Supp Fig.3E** | **Extinction day2, Tone** | RM two-way ANOVA | | Group Main Effect, F (2, 19) = 2.346 | P=0.1229 | |
| | | | | Trial Main Effect, F (9, 171) = 6.339 | P<0.0001 | |
| | | | | Group x Trial interaction, F (18, 171) = 0.8439 | P=0.6467 | |
| **Supp Fig.3E** | **Renewal, Tone** | RM two-way ANOVA | | Group Main Effect, F (2, 19) = 2.993 | P=0.0742 | |
| | | | | Trial Main Effect, F (4, 76) = 3.927 | P=0.0060 | |
| | | | | Group x Trial interaction, F (8, 76) = 0.8558 | P=0.5573 | |
| **Supp Fig.3E** | **Conditioning day1, ITI** | RM two-way ANOVA | | Group Main Effect, F (2, 19) = 0.4370 | P=0.6523 | |
| | | | | Trial Main Effect, F (9, 171) = 32.53 | P<0.0001 | |
| | | | | Group x Trial interaction, F (18, 171) = 0.9072 | P=0.5707 | |
| **Supp Fig.3E** | **Conditioning day2, ITI** | RM two-way ANOVA | | Group Main Effect, F (2, 19) = 2.790 | P=0.0866 | |
| | | | | Trial Main Effect, F (9, 171) = 3.650 | P=0.0003 | |
| | | | | Group x Trial interaction, F (18, 171) = 1.436 | P=0.1204 | |
| **Supp Fig.3E** | **Conditioning day3, ITI** | RM two-way ANOVA | | Group Main Effect, F (2, 19) = 3.090 | P=0.0689 | |
| | | | | Trial Main Effect, F (9, 171) = 0.6076 | P=0.7895 | |
| | | | | Group x Trial interaction, F (18, 171) = 1.079 | P=0.3768 | |
| **Supp Fig.3E** | **Extinction day1, ITI** | RM two-way ANOVA | | Group Main Effect, F (2, 19) = 1.581 | P=0.2317 | |
| | | | | Trial Main Effect, F (9, 171) = 3.053 | P=0.0020 | |
| | | | | Group x Trial interaction, F (18, 171) = 1.344 | P=0.1663 | |
| **Supp Fig.3E** | **Extinction day2, ITI** | RM two-way ANOVA with Bonferroni's Post Hoc test | | Group Main Effect, F (2, 19) = 3.851 | P=0.0394 | **ITI2:** Con/Fon vs. Coff/Fon (p=0.0151); **ITI2:** Con/Fon vs. Con/Foff (p=0.0012); **ITI4:** Con/Fon vs. Con/Foff (p=0.0055); **ITI18:** Con/Fon vs. Con/Foff (p=0.0185); **ITI20:** Con/Fon vs. Con/Foff (p=0.0495) |
| | | | | Trial Main Effect, F (9, 171) = 2.173 | P=0.0261 | |
| | | | | Group x Trial interaction, F (18, 171) = 2.074 | P=0.0086 | |
| **Supp Fig.3E** | **Renewal, ITI** | RM two-way ANOVA | | Group Main Effect, F (2, 19) = 1.220 | P=0.3173 | |
| | | | | Trial Main Effect, F (4, 76) = 1.508 | P=0.2082 | |
| | | | | Group x Trial interaction, F (8, 76) = 0.5764 | P=0.7942 | |

| Figure | Sub-figure | Statistical Test | One tailed or two tailed? | t, D, or F value | P value | If there is signifance, where the signifciance occurred (multiple comparison)? |
|---|---|---|---|---|---|---|
| **Supp Fig.4C** | | RM two-way ANOVA | | Tone Main Effect, F (4, 28) = 0.3200 | P=0.8622 | |
| | | | | Days Main Effect, F (2, 14) = 0.6314 | P=0.5463 | |
| | | | | Tone x Days interaction, F (8, 56) = 1.959 | P=0.0690 | |
| **Supp Fig.4D** | | RM two-way ANOVA | | Tone Main Effect, F (9, 63) = 1.775 | P=0.0908 | |
| | | | | Days Main Effect, F (1, 7) = 0.2860 | P=0.6093 | |
| | | | | Tone x Days interaction, F (9, 63) = 0.8102 | P=0.6086 | |
| **Supp Fig.4E** | | RM one-way ANOVA with Tukey's Post Hoc test | | Tone, F (4, 28) = 3.870 | P=0.0126 | Tone1 vs. Tone2 (P=0.0211); Tone1 vs. Tone3 (p=0.0200). |
| **Supp Fig.5B** | **Tone** | RM two-way ANOVA with Bonferroni's Post Hoc test | | Trial Main Effect, F (5, 190) = 87.02 | P<0.0001 | |
| | | | | Group Main Effect, F (2, 38) = 3.891 | P=0.0290 | PDyn-lox_GFP-Cre vs. WT_GFP-Cre (p=0.0304) |
| | | | | Trial x Group interaction, F (10, 190) = 0.9070 | P=0.5278 | |
| **Supp Fig.5B** | **ITI** | RM two-way ANOVA with Bonferroni's Post Hoc test | | Trial Main Effect, F (5, 190) = 81.99 | P<0.0001 | |
| | | | | Group Main Effect, F (2, 38) = 6.096 | P=0.0051 | PDyn-lox_GFP-Cre vs. WT_GFP-Cre (p=0.0351); PDyn-lox_eGFP vs. WT_GFP-Cre (p=0.0055) |
| | | | | Trial x Group interaction, F (10, 190) = 0.6769 | P=0.7451 | |
| **Supp Fig.5C** | **Context recall** | RM two-way ANOVA | | Time Main Effect, F (4, 84) = 4.272 | P=0.0034 | |
| | | | | shRNA Main Effect, F (1, 21) = 1.315 | P=0.2645 | |
| | | | | Time x shRNA interaction, F (4, 84) = 1.922 | P=0.1142 | |
| **Supp Fig.5C** | **Extinction Tone** | RM two-way ANOVA | | Trial Main Effect, F (4, 84) = 8.982 | P<0.0001 | |
| | | | | shRNA Main Effect, F (1, 21) = 0.02167 | P=0.8844 | |
| | | | | Trial x shRNA interaction, F (4, 84) = 0.9908 | P=0.4172 | |
| **Supp Fig.5C** | **Extinction ITI** | RM two-way ANOVA | | Trial Main Effect, F (4, 84) = 22.04 | P<0.0001 | |
| | | | | shRNA Main Effect, F (1, 21) = 0.9771 | P=0.3342 | |
| | | | | Trial x shRNA interaction, F (4, 84) = 0.5602 | P=0.6922 | |
| **Supp Fig.5C** | **Renewal Tone** | RM two-way ANOVA | | Trial Main Effect, F (4, 68) = 3.371 | P=0.0141 | |
| | | | | shRNA Main Effect, F (1, 17) = 3.101 | P=0.0962 | |
| | | | | Trial x shRNA interaction, F (4, 68) = 1.404 | P=0.2418 | |
| **Supp Fig.5C** | **Renewal ITI** | RM two-way ANOVA | | Trial Main Effect, F (4, 68) = 6.740 | P=0.0001 | |
| | | | | shRNA Main Effect, F (1, 17) = 0.1004 | P=0.7552 | |
| | | | | Trial x shRNA interaction, F (4, 68) = 0.6114 | P=0.6559 | |
| **Supp Fig.5D** | **Freezing before conditioning** | RM two-way ANOVA | | Cue Main Effect, F (1, 21) = 1.996 | P=0.1724 | |
| | | | | shRNA Main Effect, F (1, 21) = 1.702 | P=0.2061 | |
| | | | | Cue x shRNA interaction, F (1, 21) = 1.159 | P=0.2939 | |
| **Supp Fig.5D** | **Recall CS+** | RM two-way ANOVA | | Trial Main Effect, F (2, 44) = 2.079 | P=0.1372 | |
| | | | | shRNA Main Effect, F (1, 22) = 0.1810 | P=0.6747 | |
| | | | | Trial x shRNA interaction, F (2, 44) = 2.575 | P=0.0876 | |
| **Supp Fig. 5D** | **Recall CS-** | RM two-way ANOVA | | Trial Main Effect, F (2, 44) = 2.283 | P=0.1139 | |
| | | | | shRNA Main Effect, F (1, 22) = 1.448 | P=0.2416 | |
| | | | | Trial x shRNA interaction, F (2, 44) = 2.762 | P=0.0741 | |
| **Supp Fig. 5D** | **Recall discrimination index** | Unpaired t-test | two tailed | t=0.5406 | P=0.5945 | |
| **Supp Fig.5E** | **% time in center** | Unpaired t-test | two tailed | t=0.5794 | P=0.5685 | |

| Figure | Sub-figure | Statistical Test | One tailed or two tailed? | t, D, or F value | P value | If there is signifance, where the signifciance occurred (multiple comparison)? |
|---|---|---|---|---|---|---|
| **Supp Fig.5E** | **Distance** | RM two-way ANOVA | | Zone Main Effect, F (1, 21) = 758.9 | P<0.0001 | |
| | | | | shRNA Main Effect, F (1, 21) = 3.039 | P=0.0959 | |
| | | | | Zone x shRNA interaction, F (1, 21) = 0.6209 | P=0.4395 | |
| **Supp Fig.5F** | | Unpaired t-test | two tailed | t=1.233 | P=0.2313 | |
| **Supp Fig.5G** | **Time in open arm (%)** | Unpaired t-test | two tailed | t=0.4763 | P=0.6388 | |
| **Supp Fig.5G** | **Open arm entries** | Unpaired t-test | two tailed | t=0.1390 | P=0.8908 | |
| **Supp Fig.5G** | **Distance in open arm** | Unpaired t-test | two tailed | t=0.6522 | P=0.5213 | |
| **Supp Fig.5H** | **Air puff** | RM two-way ANOVA | | Time Main Effect, F(2711,32532)=11.14 | p<0.0001 | |
| | | | | shRNA Main Effect, F(1,12)=0.7521 | p=0.4028 | |
| | | | | Time x shRNA interaction, F(2711,32532)=0.8162 | p>0.9999 | |
| **Supp Fig.5H** | **Tail shock** | RM two-way ANOVA | | Time Main Effect, F(2711,29821)=10.57 | p<0.0001 | |
| | | | | shRNA Main Effect, F(1,11)=0.3917 | p=0.9298 | |
| | | | | Time x shRNA interaction, F(2711,29821)=0.8468 | p>0.999 | |
| **Supp Fig.5I** | | RM two-way ANOVA | | Test Main Effect, F (1, 21) = 2.343 | P=0.1408 | |
| | | | | shRNA Main Effect, F (1, 21) = 0.007956 | P=0.9298 | |
| | | | | Test x shRNA interaction, F (1, 21) = 0.1164 | P=0.7363 | |
| **Supp Fig.5J** | | Unpaired t-test | two tailed | t=1.285 | P=0.213 | |
| **Supp Fig.5K** | **Time to respond** | Unpaired t-test | two tailed | t=0.2221 | P=0.8258 | |
| **Supp Fig.5K** | **Time to jump** | Unpaired t-test | two tailed | t=0.5450 | P=0.5899 | |
| **Supp Fig.5L** | | RM two-way ANOVA with Bonferroni's Post Hoc test | | Treatment Main Effect, F (1, 29) = 152.8 | P<0.0001 | **shCtrl:** pre-CFA vs. post-CFA (p<0.0001); **shPDyn:** pre-CFA vs. post-CFA (p<0.0001). |
| | | | | shRNA Main Effect, F (1, 29) = 0.02259 | P=0.8816 | |
| | | | | Treatment x shRNA interaction, F (1, 29) = 0.2013 | P=0.6570 | |
| **Supp Fig.5M** | | RM two-way ANOVA with Bonferroni's Post Hoc test | | Treatment Main Effect, F (1, 29) = 46.26 | P<0.0001 | **shCtrl:** pre-CFA vs. post-CFA (p<0.0001); **shPDyn:** pre-CFA vs. post-CFA (p=0.0011). |
| | | | | shRNA Main Effect, F (1, 29) = 0.2041 | P=0.6548 | |
| | | | | Treatment x shRNA interaction, F (1, 29) = 1.134 | P=0.2957 | |
| **Supp Fig.6A** | **shCtrl Trial 1** | Kolmogorov-Smirnov test | | D=0.1085 | P=0.0004 | Tone 1 vs. Shock 1 |
| | | | | D=0.1864 | P<0.0001 | Tone 1 vs. ITI 1 |
| | | | | D=0.08345 | P=0.0134 | Shock 1 vs. ITI 1 |
| **Supp Fig.6A** | **shPDyn Trial 1** | Kolmogorov-Smirnov test | | D=0.1102 | P<0.0001 | Tone 1 vs. Shock 1 |
| | | | | D=0.1070 | P<0.0001 | Tone 1 vs. ITI 1 |
| | | | | D=0.1587 | P<0.0001 | Shock 1 vs. ITI 1 |
| **Supp Fig.6A** | **shCtrl Trial 6** | Kolmogorov-Smirnov test | | D=0.1335 | P<0.0001 | Tone 1 vs. Tone 6 |
| | | | | D=0.1544 | P<0.0001 | Tone 1 vs. Shock 6 |
| | | | | D=0.1446 | P<0.0001 | Tone 1 vs. ITI 6 |
| | | | | D=0.05007 | P=0.3283 | Tone 6 vs. Shock 6 |
| | | | | D=0.05563 | P=0.2158 | Tone 6 vs. ITI 6 |
| | | | | D=0.07371 | P=0.0402 | Shock 6 vs. ITI 6 |
| **Supp Fig.6A** | **shPDyn Trial 6** | Kolmogorov-Smirnov test | | D=0.08661 | P=0.0010 | Tone 1 vs. Tone 6 |
| | | | | D=0.1309 | P<0.0001 | Tone 1 vs. Shock 6 |
| | | | | D=0.06947 | P=0.0148 | Tone 1 vs. ITI 6 |
| | | | | D=0.06791 | P=0.0184 | Tone 6 vs. Shock 6 |
| | | | | D=0.06922 | P=0.0153 | Tone 6 vs. ITI 6 |
| | | | | D=0.1184 | P<0.0001 | Shock 6 vs. ITI 6 |
