## Supplementary material for "Prefrontal cortical dynorphin peptidergic transmission constrains threat-driven behavioral and network states": Key Resources Table

| REAGENT or RESOURCE | SOURCE | IDENTIFIER |
|---|---|---|
| Antibodies | | |
| Anti-ProDynorphin antibody | Abcam | Cat# ab10280 |
| Anti-GFP antibody | Abcam | Cat# ab13970 |
| Anti-Somatostatin Antibody, clone YC7 | MilliporeSigma | Cat# MAB354 |
| Alexa Fluor® 488 AffiniPure Donkey Anti-Guinea Pig IgG (H+L) | Jackson ImmunoResearch | Cat# 706-545-148 |
| Alexa Fluor® 594 AffiniPure Donkey Anti-Guinea Pig IgG (H+L) | Jackson ImmunoResearch | Cat# 706-585-148 |
| Alexa Fluor® 594 AffiniPure Donkey Anti-Rat IgG (H+L) | Jackson ImmunoResearch | Cat# 712-585-153 |
| Goat Anti-Chicken IgY H&L (Alexa Fluor® 488) preabsorbed | Abcam | Cat# ab150173 |
| Bacterial and Virus Strains | | |
| AAV2/9-phSyn1(S)-Flex-tdTomato-T2A-SypEGFP-WPRE | Boston Children's Hospital Viral Core | NA |
| AAV1-Syn-FLEX-ChrimsonR-tdTomato | UNC vector core | Lot# AV6554B |
| AAVrg-EF1α-DIO-eYFP | Addgene | Cat# 27056 |
| AAV5-EF1α-DIO-ChR2-eYFP | UNC vector core | Lot# AV4313-2A |
| AAVrg-FLEX-tdTomato | Addgene | Cat# 28306 |
| AAV-Syn-Con/Foff-eYFP | UNC vector core | Lot# AV6151 |
| AAV-Syn-Con/Fon-eYFP | UNC vector core | Lot# AV6148B |
| AAVrg-hDlx-Flex-GFP | Addgene | Cat# 83895 |
| AAV-EF1α-DIO-eYFP | UNC vector core | Lot# AV4310K |
| AAV9-syn-FLEX-jGCaMP7f-WPRE | Addgene | Cat# 104492 |
| AAV9-syn-jGCaMP7f-WPRE | Addgene | Car# 104488 |
| AAV8-EF1a-Con/Fon-GCaMP6m | Dr. Karl Deisseroth, Stanford | NA |
| AAV8-EF1a-Coff/Fon-GCaMP6m | Dr. Karl Deisseroth, Stanford | NA |
| AAV8-EF1a-Con/Foff-GCaMP6m | Dr. Karl Deisseroth, Stanford | NA |

| | | |
|---|---|---|
| AAV1-Syn-kLight 1.2 | Dr. Lin Tian, UC Davis | NA |
| AAV1-Syn-kLight 1.3 | Dr. Lin Tian, UC Davis | NA |
| AAV1-Syn-kLight 0 | Dr. Lin Tian, UC Davis | NA |
| AAV5-U6-PDyn-shRNA-GFP | Custom packaged by Vigene | NA |
| AAV5-U6-scrambled-shRNA-GFP | Custom packaged by Vigene | NA |
| AAV5-U6-PDyn-shRNA-tdTomato | Custom packaged by Vigene | NA |
| AAV5-U6-scrambled-shRNA-tdTomato | Custom packaged by Vigene | NA |
| Chemicals, Peptides, and Recombinant Proteins | | |
| (-)-U-50488 hydrochloride | Tocris | Cat# 0496 |
| Naloxone hydrochloride | Tocris | Cat# 0599 |
| Dyn A 1-17 | NIDA | Cat# MPSP-015 |
| Dyn A 2-17 | | |
| nor-Binaltorphimine dihydrochloride (nor-BNI) | Tocris | Cat# 0347 |
| Tetrodotoxin citrate (TTX) | Tocris | Cat# 1069 |
| 4-Aminopyridine (4AP) | Tocris | Cat# 0940 |
| Experimental Models: Organisms/Strains | | |
| Mouse: WT: C57BL/6J | The Jackson Laboratory | Strain #:000664 |
| Mouse: PDyn-Cre: B6;129S-*Pdyn*$^{tm1.1(cre)Mjkr}$/LowlJ | The Jackson Laboratory | Strain #:027958 |
| Mouse: Ai14: B6.Cg-*Gt(ROSA)26Sor*$^{tm14(CAG-tdTomato)Hze}$/J | The Jackson Laboratory | Strain #:007914 |
| Mouse: SST-FlpO: B6J.Cg-*Sst*$^{tm3.1(flpo)Zjh}$/AreckJ | The Jackson Laboratory | Strain #:031629 |
| Oligonucleotides | | |
| RNAscope™ Probe-Mm-Pdyn | Advanced Cell Diagnostics | Cat# 318771 |
| RNAscope™ Probe-Mm-Slc17a7-C2 | Advanced Cell Diagnostics | Cat# 416631-C2 |
| RNAscope™ Probe-Mm-Slc32a1-C3 | Advanced Cell Diagnostics | Cat# 319191-C3 |

| | | |
|---|---|---|
| RNAscope™ Probe-Mm-Sst-C2 | Advanced Cell Diagnostics | Cat# 404631-C2 |
| RNAscope™ Probe-Mm-Pvalb-C3 | Advanced Cell Diagnostics | Cat# 421931-C3 |
| RNAscope™ Probe- Hs-PDYN | Advanced Cell Diagnostics | Cat# 507161 |
| RNAscope™ Probe- Hs-SLC17A7-C2 | Advanced Cell Diagnostics | Cat# 415611-C2 |
| RNAscope™ Probe- Hs-SLC32A1-C3 | Advanced Cell Diagnostics | Cat# 415681-C3 |
| Software and Algorithms | | |
| Fiji (ImageJ) | Schneider et al., 2012 | https://imagej.net/software/fiji/ |
| GraphPad Prism 9 | GraphPad software | https://www.graphpad.com/scientific-software/prism/ |
| Clampex and Clampfit 11 | Molecular Devices | https://www.moleculardevices.com/products/axon-patch-clamp-system |
| Adobe Illustrator | Adobe | https://www.adobe.com/products/illustrator.html |
| FreezeFrame 4 and FreezeFrame 5 | Actimetrics | https://actimetrics.com/products/freezeframe/ |
| TopScan | Clever Sys Inc. | http://cleversysinc.com/CleverSysInc/csi_products/topscan-suite/ |
| ANY-maze | Stoelting | http://www.anymaze.co.uk/index.htm |
| Synapse | Tucker-Davis Technologies | https://www.tdt.com/component/synapse-software/ |
| Inscopix Data Acquisition Software | Inscopix | https://www.inscopix.com/nvoke |
| Inscopix Data Processing Software | Inscopix | https://www.inscopix.com/nvoke |
| RStudio | RStudio | https://www.rstudio.com/ |
| Python | Python Software Foundation | https://www.python.org/ |
| Bonsai | Lopes et al., 2015 | https://bonsai-rx.org/ |
